# TMED9 coordinates the clearance of misfolded GPI-anchored proteins out of the endoplasmic reticulum and into the Golgi

**DOI:** 10.1101/2024.09.27.615420

**Authors:** Elsa Ronzier, Prasanna Satpute-Krishnan

## Abstract

The p24-family member, TMED9, has recently emerged as a player in secretory pathway protein quality control (PQC) that influences the trafficking and degradation of misfolded proteins. Here we show that TMED9 plays a central role in the PQC of GPI-anchored proteins (GPI-APs). Typically, upon release from the endoplasmic reticulum (ER)-resident chaperone calnexin, misfolded GPI-APs traffic to the Golgi by an ER-export pathway called Rapid ER stress-induced Export (RESET). From the Golgi, they access the plasma membrane where they are rapidly internalized for lysosomal degradation. We used biochemical and imaging approaches in cultured cells to demonstrate that at steady-state, the majority of misfolded GPI-APs reside in the ER in association with calnexin and TMED9. During RESET, they dissociate from calnexin and increase their association with TMED9. Inhibition of TMED9’s function through siRNA-induced depletion or chemical inhibitor, BRD4780, blocked ER-export of misfolded GPI-APs. By contrast, TMED9-inhibition did not prevent ER-export of wild type GPI-APs, indicating a specific role for TMED9 in GPI-AP PQC. Intriguingly, we discovered that acute treatment with BRD4780 induced a shift in TMED9 localization away from the ER to the downstream Golgi cisternae and blocked the RESET pathway. Upon removal of BRD4780 following acute treatment, TMED9 regained access to the ER where TMED9 was able to associate with the RESET substrate and restore the RESET pathway. These results suggest that TMED9 plays a requisite role in RESET by capturing misfolded GPI-APs that are released by calnexin within the ER and conveying them to the Golgi.

## Introduction

Protein folding is an error-prone process that often generates misfolded and potentially toxic byproducts [1, 2]. To avoid the accumulation and aggregation of misfolded proteins, cells employ “protein quality control (PQC)” systems. PQC systems comprise elaborate networks of chaperones and enzymes that work together to fold and refold, sequester or destroy misfolded proteins [3, 4].

In the case of secretory pathway proteins, PQC systems monitor folding within the early secretory pathway [i.e. the endoplasmic reticulum (ER), ER-Golgi intermediate compartment (ERGIC) and Golgi] before releasing properly folded proteins to their final destination [5]. To manage the diverse topologies of secretory pathway proteins, eukaryotic cells have evolved numerous distinct PQC pathways to capture and clear misfolded products from the early secretory pathway [6]. Misfolded transmembrane or soluble secretory proteins are typically retained in the early secretory pathway and retrotranslocated to the cytosol for proteasomal degradation by the ER-associated degradation (ERAD) pathways [7, 8] or are directly transferred to lysosomes by the ER-to-lysosome-associated degradation (ERLAD) pathways, including selective ER autophagy [9–11]. By contrast, trafficking studies of misfolded GPI-APs, including naturally occurring mutants of prion protein (PrP) and artificial misfolding mutants PrP (C179A) “PrP*” and CD59 (C94S), revealed that misfolded GPI-APs are poor substrates for ERAD and instead exit the ER and access the cell surface prior to being internalized for lysosomal degradation [12, 13].

During steady-state conditions, cells constitutively release misfolded GPI-APs from the ER to the Golgi for eventual turnover in lysosomes [12, 14]. However, the flux of ER-to-Golgi clearance is dramatically enhanced by the chemical ER stressors, thapsigargin or dithiothreitol, or overexpression of other ER PQC substrates [12, 14]. Thus, we referred to the ER-clearance pathway as “RESET” for “Rapid ER Stress-induced ExporT” [12]. RESET involves dissociation from the ER-resident chaperone calnexin, and association with p24-family members, including TMP21 [12, 14, 15]. The p24-family is composed of type I transmembrane proteins that are conserved between yeast and mammals, form hetero-oligomers, and associate with COPI and COPII coat proteins to mediate vesicular trafficking between the ER and Golgi [16–19]. SiRNA knockdown of p24-family member TMP21 or TMED2, or treatment with chemical inhibitor, brefeldin A, completely blocks ER to Golgi export of misfolded GPI-APs [12, 20], indicating a role for p24-family hetero-oligomers and the conventional secretory pathway in RESET, respectively.

Several recent findings have provided compelling evidence for the involvement of the p24-family member, TMED9, in regulating diverse aspects of PQC in the early secretory pathway [21]. In particular, Dvela-Levitt *et al.* discovered that TMED9 traps the disease mutant of the secretory protein mucin-1, called MUC1-fs, in TMED9-enriched compartments [22]. Importantly, they identified a small molecule, BRD4780, that induces the acute degradation of TMED9 and concurrently allows for the specific degradation of MUC1-fs but not wild type mucin-1 [22]. Follow up work by Xiao *et al.* identified closely interacting regions of the TMED9 GOLD domain and MUC1-fs through AlphaFold modeling and disulfide crosslinking experiments [23]. Additionally, Roberts *et al.* found TMED9, along with TMP21, preferentially associates with the misfolding Z mutant of alpha-1-antitrypsin (ATZ) over the wild type variant, and is required for ATZ clearance from the ER [24]. Furthermore, Zavodszky and Hegde discovered that PrP* co-immunoprecipitates with TMED9, along with TMP21 and TMED2, and that partial knockdown of TMED9 partially reduced the ER-export of PrP* [20].

In this study, we sought to elucidate TMED9’s role in ER-clearance of misfolded GPI-APs by using a combination of biochemical and imaging approaches in cells. We demonstrated that properly folded GPI-APs are able to traffic from the ER to the cell surface regardless of TMED9 expression or knock down. Conversely, inhibiting TMED9 with siRNA or BRD4780 specifically blocked ER-export of misfolded GPI-APs. Critically, we provide evidence that in order to execute its role in RESET, TMED9 must be localized in the ER where it can bind to the RESET substrates. Complete degradation of TMED9 blocks RESET. Similarly, acute mislocalization of TMED9 from the ER to the Golgi blocks RESET. However, relocalization of TMED9 to the ER restores the RESET pathway. Altogether, the results we present here strongly suggest that TMED9, in association with the known RESET factors TMP21 and TMED2, participates in RESET by capturing misfolded GPI-APs that are released by calnexin in the ER and coordinating their transport to the Golgi.

## Results

### Rationale behind our model substrate, system and strategy to dissect the role of TMED9 in RESET

To dissect the role of TMED9 in RESET, we used a mutant variant of YFP-PrP WT, called “YFP-PrP*” as our primary misfolding model substrate. YFP-PrP WT includes the N-terminal prolactin signal sequence, which drives efficient translocation into the ER [25], and yields a homogenous population of PrP that efficiently folds and traffics to the cell surface [13, 26]. YFP-PrP* is derived from YFP-PrP WT but contains a C179A mutation that disrupts the single disulfide bond in PrP. YFP-PrP* was stably expressed in an NRK cell line “YFP-PrP* NRK” at levels similar to endogenous PrP in mouse brain lysates [12, 14]. Here, we verified our previous observations that in YFP-PrP* NRK cells, YFP-PrP* proteins were strongly localized to the ER and poorly localized on the plasma membrane at steady-state (Supp Figure S1A-B and [12]).

GFP-CD59 (C94S) is another misfolded GPI-AP model substrate that contains a single cysteine mutation that disrupts 1 of 5 possible disulfide bonds [15, 27]. Previously, we tested for the universality of the major findings that we made using YFP-PrP* as the model substrate by repeating the critical experiments with the unrelated, misfolded GPI-AP GFP-CD59 (C94S) [12, 14]. Here we show that GFP-CD59 (C94S) NRK cells expressed a heterogenous population of GFP-CD59 (C94S) that included clearly detectable ER and plasma membrane-localized populations (Supp Figure S1C-D). Thus, we opted to use the more homogenous population of YFP-PrP* NRK cells as our primary model system to study RESET, and GFP-CD59 (C94S) as a secondary model system.

Misfolded GPI-APs are exported out of the ER via RESET for subsequent lysosomal turnover during steady-state conditions [12]. However, various ER-stressors including the induction of new glycoprotein expression or chemical ER-stressors enhance the flux of misfolded GPI-APs through RESET [12, 14, 20]. Thapsigargin (TG)-treatment, in particular, induces the immediate and near-synchronized release of YFP-PrP* or GFP-CD59 (C94S) through the RESET pathway [12]. The misfolded GPI-APs traffic from the ER to the Golgi within 30 min, and to lysosomes for degradation within 90 min in NRK cells [12]. Thus, for this study, we used TG-treatment in our YFP-PrP* or GFP-CD59 (C94S) NRK cell lines as a tractable and facile system to assay for the integrity of the RESET pathway.

### During RESET, misfolded GPI-APs release calnexin and increase binding to TMED9 and TMP21

We set out to validate and expand upon the previously reported finding that TMED9 associates with GFP-PrP* at steady-state and may be involved in its clearance from the ER via the RESET pathway in TG-treated cells [20]. We confirmed that during steady-state conditions, YFP-PrP* co-immunoprecipitates with TMED9, along with calnexin, TMP21, and TMED2 (Supp Figure S1E), as previously published [12, 20]. Similarly, we found that the alternate model RESET substrate, GFP-CD59 (C94S), co-immunoprecipitated with TMED9, calnexin, TMP21, and TMED2 at steady-state (Supp Figure S1F).

To dissect YFP-PrP* protein-interactions as YFP-PrP* undergoes RESET, we synchronized the release of YFP-PrP* for RESET with TG-treatment and correlated live-cell time-lapse imaging with co-immunoprecipitation time courses. The majority of YFP-PrP* molecules was previously shown to have been cleared from the ER to the Golgi via the RESET pathway in under 30 min of TG-treatment [12]. Here, we reproduced this finding using YFP-PrP* NRK cells that were either co-transfected with CER-CNX, a Cerulean (CER)-tagged version of the ER-resident chaperone calnexin to mark the ER or CER-GalT to mark the Golgi. Within the first 20 min of TG-treatment, YFP-PrP* actively trafficked out of the ER into the Golgi leading to a significant decrease in the colocalization of YFP-PrP* with CER-CNX and significant increase in the colocalization of YFP-PrP* with CER-GalT by 20 min (Figure 1A-D). Additionally, we imaged the trafficking of YFP-PrP* during RESET in cells co-transfected with CER-TMED9 and Golgi marker, FusionRed (FusRed)-SiT. Again, within 20 min of TG-treatment, YFP-PrP* moved into the FusRed-SiT-marked Golgi (Figure 1E-F, Video 1). Intriguingly, TG-treatment did not alter the overall distribution of CER-TMED9 molecules over the course of the 60 min time-lapse videos. Moreover, the Pearson’s colocalization coefficient between CER-TMED9 and FusRed-SiT did not significantly change from the 0 to the 20 min time point after TG-treatment, despite the dramatic shift in YFP-PrP* localization from the ER to the Golgi within that time frame (Figure 1E-F; Video 1).

**Figure 1:**
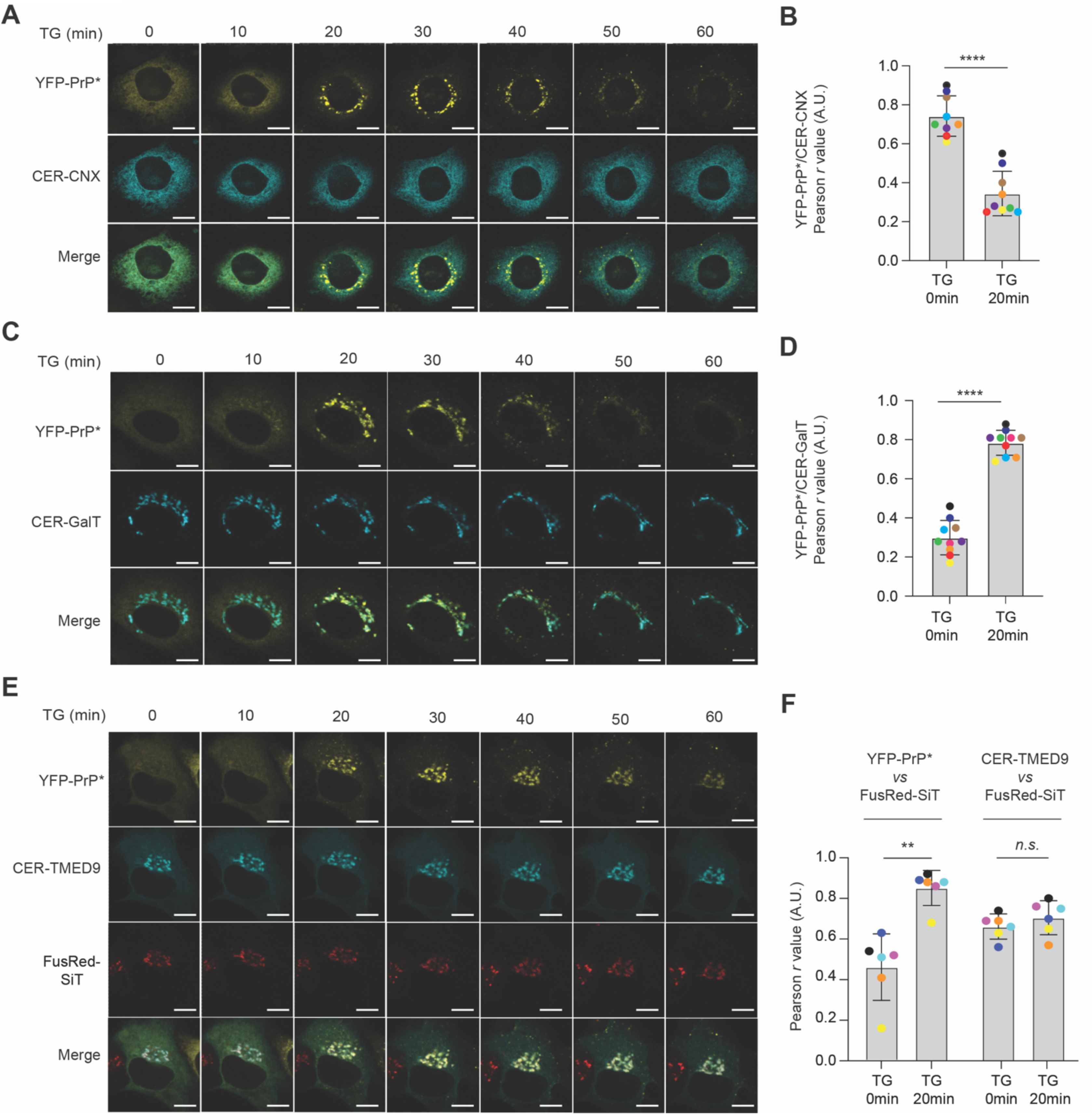
During RESET, misfolded GPI-APs traffic from the ER to the Golgi. **(A)** Time-lapse imaging sequence collected immediately after thapsigargin (TG)-treatment of a typical YFP-PrP* NRK cell that was co-expressing ER-marker, Cerulean-calnexin (CER-CNX). Scale bar represents 10 µm. **(B)** Plot of the average Pearson’s r values between YFP-PrP* and CER-CNX for 9 individual cells across the 0 and 20 min time points of time-lapses collected after addition of TG, as shown in (A). Pearson’s colocalization coefficients, r, were measured between YFP-PrP* and CER-CNX within the boundaries of the cell. For each image, the boundary of the cell was revealed by temporarily increasing the gain for CER-CNX. The Pearson’s r values are color-coded to signify each individual cell at different time points. The r values for 0 min and 20 min time points were 0.742 ± 0.098 and 0.344 ± 0.108, respectively. **(C)** Time-lapse imaging sequence collected immediately after TG-treatment of a typical YFP-PrP* NRK cell that was co-expressing the Golgi marker, CER-GalT. Scale bar represents 10 µm. **(D)** Plot of the average Pearson’s r values between YFP-PrP* and CER-GalT for 10 individual cells across the 0 and 20min time points of time-lapses collected after addition of TG, as shown in (C). Pearson’s colocalization coefficients, r, were measured between YFP-PrP* and CER-GalT within the boundaries of the cell. For each image, the boundary of the cell was revealed by temporarily maximizing the gain for YFP-PrP*. The Pearson’s r values are color-coded to signify each individual cell at different time points. The r values for 0 min and 20 min time points were 0.300 ± 0.084 and 0.785 ± 0.060, respectively. **(E)** Time-lapse images collected immediately after TG-treatment of a typical YFP-PrP* NRK cell that was co-expressing CER-TMED9 and FusionRed (FusRed)-SiT as a Golgi-marker. Scale bar represents 10 µm. This figure panel is associated with Video 1. **(F)** Plot of the average Pearson’s r values between YFP-PrP* and FusRed-SiT or CER-TMED9 and FusRed-SiT, as indicated, for 6 individual cells across the 0 and 20 min time points of time-lapses collected after addition of TG, as shown in (E). Pearson’s colocalization coefficients, r, were measured between YFP-PrP* and FusRed-SiT or CER-TMED9 and FusRed-SiT, within the boundaries of the cell. For each data point at 0 and 20 min time points, the boundary of the cell was revealed by temporarily maximizing the gain for YFP-PrP*. The Pearson’s r values are color-coded to signify each individual cell at different time points. The r values between YFP-PrP* and FusRed-SiT for 0 min and 20 min time points were 0.462 ± 0.149 and 0.852 ± 0.079, respectively. The r values between CER-TMED9 and FusRed-SiT for 0 min and 20 min time points were 0.662 ± 0.057 and 0.705 ± 0.077, respectively.

Next we examined the dynamics of misfolded GPI-AP and TMED9 protein-protein interactions during RESET by treating YFP-PrP* NRK cells with TG, co-immunoprecipitating YFP-PrP* and analyzing the changes in YFP-PrP*’s associations with TMED9, TMP21 and calnexin over time. During steady-state conditions, YFP-PrP* associates with calnexin, TMED9 and TMP21. However, within 20 min of TG-treatment, densitometry measurements of the western blot bands revealed that YFP-PrP*’s association with calnexin declined by 57±9% (Stdev.p, n=3) and concurrently YFP-PrP*’s association with TMED9 and TMP21 increased by 335±25% (Stdev.p, n=3) and 267±91% (Stdev.p, n=3), respectively (Figure 2A-B). From 20 min to 40 min after addition of TG, YFP-PrP*’s association with TMED9 and Tmp21 declined.

**Figure 2:**
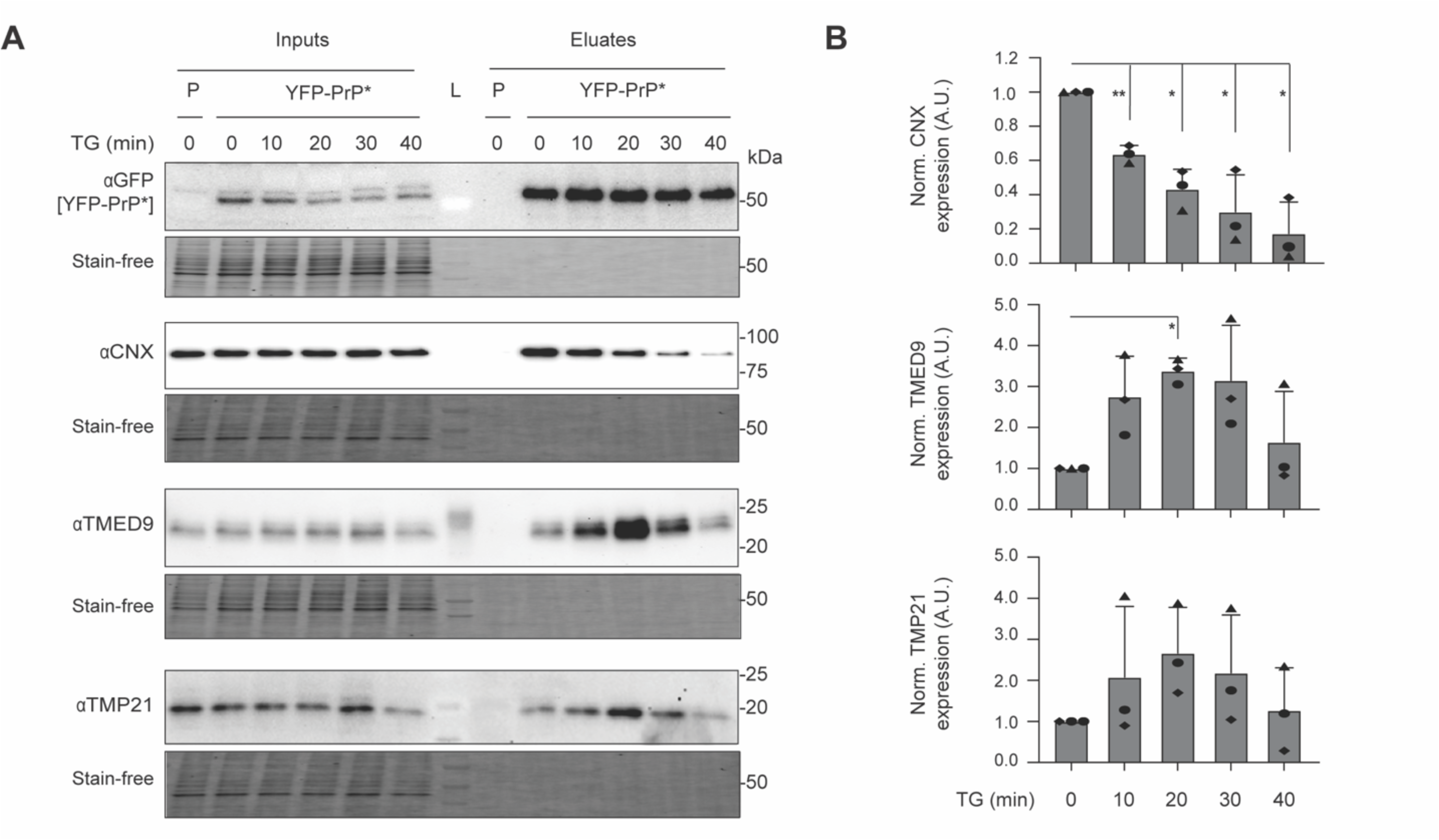
During RESET, misfolded GPI-APs dissociate from calnexin and increase association with TMED9 and TMP21. **(A)** Representative western blots of GFP-tag co-immunoprecipitations from the parental untransfected NRK cells (P) or stably transfected YFP-PrP* NRK cells (n=3 biological replicates). YFP-PrP* NRK cells were treated with thapsigargin (TG) and collected for co-immunoprecipitation at the indicated time points. In addition to using anti-GFP antibody to detect co-immunoprecipitation of YFP-PrP*, blots were probed for calnexin (CNX), TMED9 and TMP21. CNX and TMP21 were blotted on the same membrane. “L” indicates the lane that the ladder was loaded in. Under each western blot is depicted a “Stain-Free” image of the total protein in the gel. **(B)** Bar graph representing the mean band intensity for CNX, TMED9 and TMP21 in the eluates from 3 independently performed experiments as shown in (A). For each time point, CNX, TMED9 and TMP21 eluate band intensities were double normalized. First, they were normalized by the band intensity of eluate YFP-PrP*. Second, they were normalized against eluate time 0 band intensity of each protein probed. Error bars represent standard deviations of the mean of 3 independent experiments. Symbols were coded for each experiment. Statistics were calculated from unpaired t-test with Welch’s correction with * indicated p<0.05 and ** indicated p<0.01.

Synthesis of the data from Figure 1, Video 1 and Figure 2 demonstrate the following points. Within the first 20 min after TG-treatment when YFP-PrP* was actively undergoing RESET (i.e. clearing out of the ER to the Golgi), YFP-PrP* dissociated from calnexin and increased its associations with TMED9 and TMP21. After 20 min of TG-treatment, during the time when most of the YFP-PrP* was moving through and out of the Golgi towards the plasma membrane, YFP-PrP*’s associations with TMED9 and TMP21 declined.

Based on the steady-state association of TMED9 with diverse unrelated misfolded proteins shown in Supp Figure S1E-F and in other reports [20, 22, 24], we hypothesized that TMED9 may be a PQC factor. In particular, the dynamic association between TMED9 and YFP-PrP* that is heightened during the period that YFP-PrP* was actively undergoing RESET shown in Figures 1 and 2 implicates TMED9’s involvement in the RESET pathway. Therefore, we set out to test whether TMED9 plays an essential role in coordinating RESET of misfolded GPI-APs.

### siRNA depletion of TMED9 inhibits RESET and consequent access to the cell surface and degradation of misfolded GPI-APs

To test for a role of TMED9 in RESET, we used siRNA to deplete TMED9 levels. We examined the impact of TMED9 siRNA (siTMED9)-depletion on RESET at the population level using western blotting and live-cell imaging techniques, and at the individual cell level using immunofluorescence. For the below siTMED9 experiments, we exploited our previous observation that TG-treatment of YFP-PrP* NRK cells induces the rapid and synchronized ER-to-Golgi clearance of YFP-PrP*, followed by traffic to the cell surface where YFP-PrP* is internalized for near complete lysosomal degradation within 90 min [12]. Critical to the interpretation of the population-level TMED9-depletion experiments, immunofluorescence revealed that siTMED9-treatment effectively depleted TMED9 in 65-70% of the cells, but a minority of cells continued to express TMED9 (Supp Figure S2A).

First, we examined the effect of TMED9-depletion on TG-induced clearance of YFP-PrP* at the population level by western blot. In control cells (CTL) that were treated with scrambled siRNA, TG induced a 86.3±6.3% (Stdev.p, n=3) decrease in the levels of YFP-PrP* within 90 min (Figure 3A-B, compare lanes 1 and 2). On the other hand, in the siTMED9-treated cells, TG induced a 34.2±10.52% (Stdev.p, n=3) decrease in the levels of YFP-PrP* (Figure 3A-B, compare lanes 3 and 4), indicating a role for TMED9 in the efficient degradation of YFP-PrP*. We observed a slight and insignificant increase to 113.88±17.28% (Stdev.p, n=3) in the steady-state levels of YFP-PrP* in siTMED9 cells when compared to CTL cells (Figure 3A-B, compare lanes 1 and 3). These data show that at a population level, TMED9-depletion reduces the efficiency of YFP-PrP* clearance.

**Figure 3:**
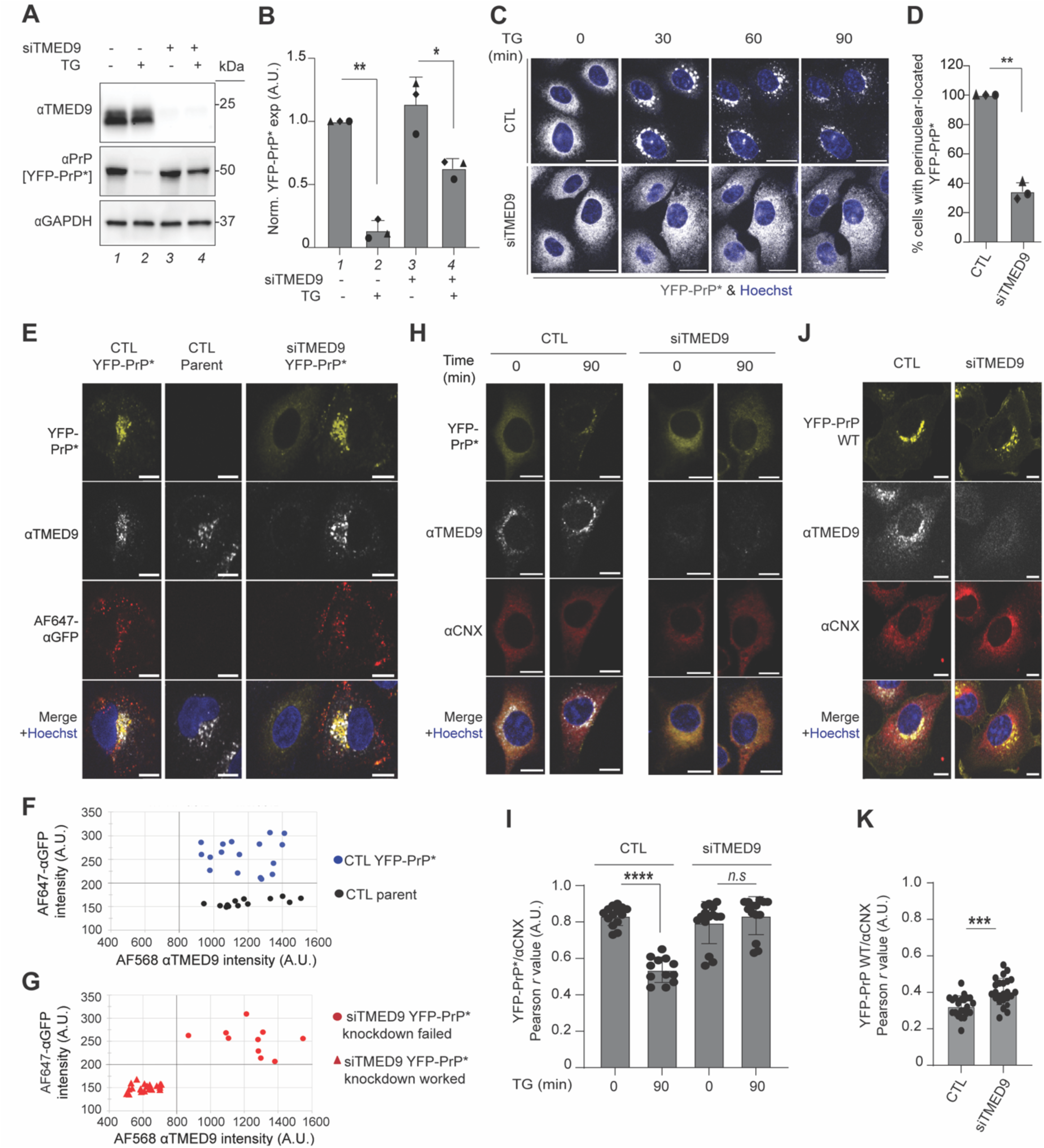
siRNA depletion of TMED9 blocks ER-export of misfolded GPI-APs but not wild type GPI-APs. **(A)** Representative western blots of control (siTMED9 –) or siTMED9-treated (siTMED9 +) YFP-PrP* NRK cells that were either untreated (TG –) or treated with TG for 90 min (TG +) (n=3 biological replicates). Blots were probed for TMED9, PrP to detect YFP-PrP* and GAPDH. Each lane was numbered to facilitate cross-referencing with the quantification shown in (B). **(B)** Bar graph representing the mean band intensity for YFP-PrP* from 3 biological replicates as shown in (A). For each condition, YFP-PrP* band intensities were double normalized. First, they were normalized by the band intensity of GAPDH. Second, they were normalized against the untreated control (“siTMED9 – TG – “) band intensity. Error bars represent standard deviation of the mean. Symbols were coded for each independently performed experiment. Statistics were calculated from unpaired t-test with Welch’s correction with * indicated p<0.05 and ** indicated p<0.01. The bars were numbered 1-4 to facilitate cross-referencing with the representative blot in (A). **(C)** Time-lapse images of control (CTL) or TMED9 siRNA (siTMED9)-treated YFP-PrP* NRK cells. Prior to starting the time-lapses, nuclei were stained with Hoechst. Image-collection was started immediately after the addition of thapsigargin (TG). Scale bar represents 20 µm. **(D)** Percentage of cells as represented in (C) with a perinuclear Golgi-pattern of YFP-PrP* after 30 min of TG treatment. Only cells which were flat and adherent were counted. Rounded up cells undergoing mitosis were not counted. The bar graph represents 3 biological replicates. Symbols were coded for each independently executed experiment. The number of cells analyzed for each biological replicate are as follows (CTL: triangle 29, diamond 56, circle 51; siTMED9: triangle 69, diamond 58, circle 61). Error bars represent standard deviation of the mean. Statistics were calculated from unpaired t-test with ** indicated p<0.01. **(E)** Confocal images of control scrambled siRNA (CTL) treated YFP-PrP* NRK cells or parental NRK cells, or siTMED9-treated YFP-PrP* NRK cells that were incubated with thapsigargin (TG), Alexa Fluor 647-conjugated anti-GFP antibodies and bafilomycin A1 (BAF) for 90 minutes, fixed and stained for TMED9. An Alexa Fluor^TM^ (AF) 568-conjugated secondary was used for the TMED9 immunofluorescence. Nuclei were stained with Hoechst. All images were taken with identical imaging parameters. Scale bar represents 10 µm. **(F)** Plot of background-subtracted mean intensities measured in arbitrary units (A.U.) for TMED9 immunofluorescence and AF647-αGFP Ab fluorescence of scrambled siRNA-treated control (CTL) parental NRK (n=14) or CTL YFP-PrP* NRK cells (n=17), as shown in (E). Cells were handled and images collected as shown and described in (E). ROIs around the periphery of each cell was drawn by temporarily increasing the gain in the 568 channel (TMED9-staining). We observed that in addition to strong specific TMED9 binding, a small fraction of the primary antibody against TMED9 appeared to non-specifically bind the entire cytoplasm and we took advantage of that to trace the outline of the cell and measure the mean intensities for both 568 nm (αTMED9) and 647 nm (AF647-αGFP) channels. Both the scrambled siRNA control (CTL)-treated parental NRK cells and CTL YFP-PrP* NRK cells served as positive controls for wild type TMED9 expression. A threshold (shown as bolded vertical grid line) for wild type TMED9 levels was placed to the left of the lowest expressing scrambled siRNA-treated (CTL) cell. CTL parental NRK served as a negative control for AF647-αGFP Ab uptake. The CTL YFP-PrP* NRK cells served as positive control for antibody uptake. A threshold (shown as bolded horizontal grid line) for antibody uptake was placed below the lowest level of AF647-αGFP Ab mean fluorescence intensity of the positive control CTL YFP-PrP* NRK, but above the highest level of AF647-αGFP Ab mean fluorescence intensity in the negative control CTL untransfected parental NRK. **(G)** Plot of background-subtracted mean intensities measured in arbitrary units (A.U.) for TMED9 immunofluorescence and AF647-αGFP Ab fluorescence inside of siTMED9-treated YFP-PrP* NRK cells (n=33), as shown in (E). The data points were plotted on an identically proportioned graph to that in (F) so that direct comparisons of the data points could be made with the controls plotted in (F). Cells were handled and images collected as shown and described in (E). ROIs around the periphery of each cell was drawn and background-subtracted mean intensities measured for the 568 and 647 channels exactly as described in (F). 10 of the data points had mean TMED9 intensities within the wild type levels and were coded with a circle symbol to represent “siTMED9 YFP-PrP* NRK knockdown failed.” 23 of the data points had mean TMED9 below the wild type levels and were coded with a triangle symbol to represent “siTMED9 YFP-PrP* NRK knockdown worked.” As shown in (E), in siTMED9-treated cells where knockdown worked and YFP-PrP* ER-export was blocked, the bright Golgi-patterned perinuclear staining was lost, but the low level of non-specific cytoplasmic binding persisted. **(H)** Immunofluorescence images of endogenous TMED9 in control siRNA or siTMED9-treated YFP-PrP* NRK cells that were depleted for TMED9. For each condition, cells were treated with thapsigargin (TG) for 0 min or 90 min before fixation and staining for TMED9 and ER-resident chaperone, calnexin (CNX). Nuclei were stained with Hoechst. Larger fields of views and additional data for these experiments are presented in Supp. Figure S3A. Scale bar represents 10 µm **(I)** Plot of the average Pearson’s r values between YFP-PrP* and endogenous calnexin (CNX) for control (CTL) scrambled siRNA or siTMED9-treated cells at 0 min or 90 min after thapsigargin addition, as shown in (H). The number of cells analyzed for each condition are listed as follows: CTL siRNA 0 min post-TG-treatment (n=14), CTL siRNA 90 min post-TG-treatment (n=12), siTMED9 0 min post-TG-treatment (n=15), siTMED9 90 min post-TG-treatment (n=14). TMED9-depleted cells were identified by their lack of perinuclear Golgi-patterned TMED9 staining. Pearson’s colocalization coefficients, r, were measured between YFP-PrP* and CNX, within the boundaries of the cell as defined by the outline of the CNX staining. In CTL cells, the r values between YFP-PrP* and CNX for 0 min and 90 min time points were 0.834 ± 0.051 and 0.538 ± 0.066, respectively. In siTMED9-treated cells, the r values between YFP-PrP* and CNX for 0 min and 90 min time points were 0.797 ± 0.112 and 0.835 ± 0.100, respectively. **(J)** Immunofluorescence images of endogenous TMED9 in control (CTL) or TMED9 siRNA (siTMED9)-treated YFP-PrP WT NRK cells that were depleted for TMED9. Cells were fixed and stained for TMED9 and ER-resident chaperone, calnexin (CNX) at steady-state conditions. No drugs were included. Nuclei were stained with Hoechst. Scale bar represents 10 µm. **(K)** Plot of the average Pearson’s *r* values between YFP-PrP WT and endogenous calnexin (CNX) for control (CTL) scrambled siRNA or siTMED-treated cells at steady-state, as shown in (J). The number of cells analyzed for each condition are listed as follows: scrambled siRNA (CTL) (n=22) and siTMED9 (n=24). TMED9-depleted cells were identified by their lack of perinuclear Golgi-patterned TMED9 staining. Pearson’s colocalization coefficients, *r*, were measured between YFP-PrP* and CNX, within the boundaries of the cell as defined by the outline of the CNX staining. The *r* values between YFP-PrP WT and CNX in CTL cells and in siTMED9-treated cells were 0.325 ± 0.056 and 0.414 ± 0.066, respectively.

Second, we examined the impact of TMED9-depletion on RESET at the population level by live-cell time-lapse imaging. To do this, we counted the number of YFP-PrP* NRK cells that demonstrated TG-induced ER-export of YFP-PrP* by deducing the ER and Golgi localizations of YFP-PrP* based on the YFP-fluorescent patterns. The ER is well established to be the largest contiguous membrane-bound organelle that fills the cytoplasmic space and includes the nuclear envelope in mammalian cell culture [28]. In adherent cultured cells such as the NRK cells used here, the Golgi is composed of perinuclear stacks of cisternae with a characteristic appearance of a crescent-shaped ribbon that hugs part of the nucleus [29–32]. As shown previously for YFP-PrP* NRK cells in the context of organelle markers, ER localized YFP-PrP* displays a cytoplasm-filling reticulate pattern that surrounds the nucleus, while Golgi localized YFP-PrP* displays a juxtanuclear, crescent-shaped pattern that we refer to as a “perinuclear Golgi-pattern” (Supp Figure S1A, Figure 1 and [12]). In control (CTL) scrambled siRNA-treated YFP-PrP* NRK cells, addition of TG resulted in the rapid relocalization of YFP-PrP* from the ER to the Golgi within 30 min in 100% of the cells (n=136, Figure 3C-D). On the other hand, in siTMED9-treated YFP-PrP* NRK cells, relocalization of YFP-PrP* from the ER to the Golgi only occurred in a minor fraction of the cells (34.4%, n=180). In the majority (65.6%, n=180) of the siTMED9-treated cells, YFP-PrP* remained in the ER for at least 90 min after TG-treatment (Figure 3C-D). This demonstrates a block in RESET in the majority of siTMED9-treated cells.

As an orthogonal approach, we examined the impact of TMED9-depletion on RESET at the population level by antibody uptake assay. Consequent to export out of the ER to the Golgi via RESET, misfolded GPI-APs transiently access the cell surface prior to being internalized for lysosomal degradation [12, 14, 20]. This phenomenon occurs at steady-state or during ER stress [12, 14, 20]. To exploit antibody (Ab)-uptake as a visible readout for traffic of YFP-PrP* through RESET, we used Alexa Fluor^TM^ 647-conjugated anti-GFP antibodies (AF647-αGFP Abs), which bind specifically to cell-surface exposed YFP-tagged misfolded GPI-APs, as published previously [12]. The AF647 continues to fluoresce inside of cells that are treated with Bafilomycin A1 (BAF), a chemical inhibitor of lysosomal acidification and degradation [12, 33]. Incidentally, BAF also inhibits lysosomal degradation of YFP-PrP*, allowing it to be detected in lysosomes after RESET [12]. Thus, we incubated CTL YFP-PrP* NRK cells, CTL parental NRK cells (parent) or siTMED9-treated YFP-PrP* with TG + BAF + AF647-αGFP Abs and collected time-lapse images (Supp. Figure S2B-D, Video 2). All of the CTL YFP-PrP* NRK cells internalized AF647-αGFP Ab (n=100) (Supp. Figure S2B). The untransfected CTL parental NRK cells did not internalize AF647-αGFP Ab (n=100) (Supp. Figure S2C). On the other hand, for the siTMED9-treated YFP-PrP* NRK cells, only 33% (n=100) internalized AF647-αGFP Abs (Supp. Figure S2D, Video 2). The inhibition of Ab-uptake in the majority of siTMED9-treated YFP-PrP* NRK cells demonstrated that TMED9-expression is important for efficient ER-export via the RESET pathway.

Third, we examined the impact of TMED9-depletion on RESET at the single-cell level by immunofluorescence. To assess the effect of TMED9-depletion on Ab-uptake by YFP-PrP* in individual cells, we first set baseline mean intensities for anti-TMED9 Ab immunofluorescence and AF647-αGFP Ab fluorescence inside of scrambled siRNA-treated control (CTL) parental NRK or CTL YFP-PrP* NRK cells that were incubated with TG+BAF+AF647-αGFP Ab for 90 min prior to fixation (Figure 3E,F). CTL parental NRK cells served as a negative control for AF647-αGFP Ab uptake because they do not express YFP-PrP* (Figure 3E,F, and [12]). CTL YFP-PrP* NRK cells served as positive control for antibody uptake (Figure 3E,F, and [12]). Both CTL parental NRK and CTL YFP-PrP* NRK cells serve as positive controls for TMED9 expression. Of the 33 randomly selected siTMED9-treated YFP-PrP* NRK cells that we analyzed for TMED9 expression, we found that 10 of them demonstrated perinuclear Golgi-pattern TMED9 staining, indicating incomplete knockdown, while the remaining 23 were depleted for TMED9 (Figure 3E,G). Importantly, in siTMED9-treated YFP-PrP* NRK cells, depletion of TMED9 to levels that were quantitatively below the TMED9 levels in CTL YFP-PrP* NRK cells resulted in visible and measurable inhibition of antibody-uptake (Figure 3E-G). This demonstrates the dependence of YFP-PrP* on TMED9 for ER-export to the cell surface on a single cell-by-cell basis.

Finally, we examined the effect of TMED9-depletion on RESET at the individual cellular level more directly by assessing ER-export of YFP-PrP*. YFP-PrP* NRK cells that were either treated with scrambled siRNA control (CTL) or siTMED9 were fixed at 0 min or 90 min timepoints after TG-treatment, and co-stained for TMED9 and calnexin. Calnexin-staining served as an ER marker. In CTL YFP-PrP* NRK cells, YFP-PrP* visibly localized to the ER at steady-state and was cleared out of the ER after TG-treatment (Figure 3H, Supp Figure S3A). Colocalization analysis of YFP-PrP* and calnexin revealed that this correlated with a drop in Pearson’s colocalization coefficients, r, from 0.834 +/-0.05 (n=14 cells) at steady-state to Pearson’s r value of 0.538 ± 0.067 (n=12 cells) after TG-treatment (Figure 3I). By contrast, for siTMED9 YFP-PrP* NRK cells in which TMED9-levels were depleted below wild type levels, YFP-PrP* appeared to remain localized to the calnexin-marked ER despite TG-treatment (Figure 3H, Supp Figure S3A). Accordingly, the Pearson’s r value was not significantly altered by TG-treatment. The Pearson’s r value between YFP-PrP* and calnexin in TMED9-depleted cells was 0.80 ± 0.11 (n=15 cells) at steady-state and 0.84 ± 0.10 (n=14 cells) in TG-treated cells (Figure 3I). This demonstrates that YFP-PrP* relies on TMED9 to leave the ER via the RESET pathway.

As with YFP-PrP* (Figure 3 and Supp. Figure S2), siTMED9-treatment appeared to inhibit RESET of the alternate misfolded GPI-AP, GFP-CD59 (C94S) (Supp Figure S3B-E). In comparison to control scrambled siRNA-treated cells, siTMED9 depleted GFP-CD59 (C94S) NRK cells exhibited (1) inefficient TG-induced clearance of GFP-CD59 (C94S), as shown through western blot analysis (Supp Figure S3B-C), (2) increased steady-state levels of GFP-CD59 (C94S), as shown by western blot (Supp Figure S3B-C), and (3) a significant decrease in the percentage of cells in which GFP-CD59 (C94S) underwent RESET upon TG treatment, as shown by live-cell imaging (Supp Figure S3D-E). The demonstration that TMED9-depletion inhibited ER-export of two unrelated misfolded GPI-APs suggest that TMED9 could play a general role in the ER-export step of the RESET pathway for misfolded GPI-APs.

### siRNA depletion of TMED9 does not prevent wild type GPI-APs from leaving the ER and trafficking to the plasma membrane

As a control to test for general effects of TMED9-depletion on ER-export, we used YFP-PrP WT. YFP-PrP WT folds efficiently and traffics via the secretory pathway from the ER to the plasma membrane where it stably resides and is predominantly localized at steady-state [12]. Between control and siTMED9-treated YFP-PrP WT NRK cells, we observed no obvious difference in YFP-PrP WT localization in the post-ER secretory pathway compartments and the plasma membrane. To confirm whether YFP-PrP WT was exposed on the cell surface regardless of TMED9-expression, we incubated live control and siTMED9-treated, unpermeabilized cells with AF594-αGFP antibodies (Abs) for 10 min prior to fixation. The AF594-αGFP Abs would only have access to surface exposed YFP-PrP WT. Then we fixed and stained the cells for TMED9 to detect siTMED9 knockdown efficiency (Supp Figure S3F-H). Additionally, we used CTL parental NRK cells as a negative control for AF594-αGFP Ab-binding because they do not express YFP-PrP WT. CTL YFP-PrP WT NRK cells served as positive control for AF594-αGFP Ab-binding (Supp Figure S3F-G). Both CTL parental NRK and CTL YFP-PrP WT NRK cells serve as positive controls for TMED9 expression. Of the 45 randomly selected siTMED9-treated YFP-PrP WT NRK cells that we analyzed for TMED9 expression, we found that 10 of them demonstrated a perinuclear Golgi-pattern for TMED9 staining, indicating no knockdown, while the remaining 35 were depleted for TMED9 (Supp Figure S3F, H). Importantly, regardless of siTMED9-treatment and TMED9-expression levels, the AF594-αGFP antibodies bound to the YFP-PrP WT NRK cells at levels above the CTL parental NRK cells (Supp Figure S3F-H). This experiment demonstrated that YFP-PrP WT is able to leave the ER and traffic to the cell surface independently of TMED9 expression.

As an alternate approach to analyze YFP-PrP WT localization in the presence or absence of TMED9, we fixed the cells and co-stained them for TMED9 expression and endogenous calnexin to mark the ER, which fills the cytoplasmic space. We counted the number of cells in which YFP-PrP WT localized to the plasma membrane, which surrounds the cytoplasm. In 100% of the TMED9-expressing cells (n=28), YFP-PrP WT visibly localized to the plasma membrane (Figure 3J). Similarly, in 100% of the TMED9-depleted cells (n=26), YFP-PrP WT visibly localized to the plasma membrane (Figure 3J), demonstrating that TMED9 is not required for ER-export of the wild type variant. Further support for the idea that TMED9 is not required for ER-export of wild type GPI-APs was provided by Pearson’s colocalization analysis of YFP-PrP WT with calnexin, which revealed similar r values of 0.325 ± 0.056 for n=22 control scrambled siRNA-treated (CTL) cells and 0.404 ± 0.100 for n=24 siTMED9-treated cells (Figure 3K). The slight increase in r value between YFP-PrP WT and calnexin in siTMED9-treated cells over CTL cells may indicate a delay in ER-export (discussed below). Critically, we show here that YFP-PrP WT localization does not require TMED9 for ER-export to post-ER compartments including the plasma membrane (Figure 3J-K). This is in marked contrast to the misfolded variant, YFP-PrP*, which relies on TMED9 expression for ER-export (Figure 3E-I).

Taken together, the results from the siTMED9 knockdown experiments demonstrate that TMED9-expression is required for the ER-export of misfolded GPI-APs. On the other hand, wild type GPI-APs traffic to the plasma membrane regardless of TMED9-expression or depletion. This indicates that the general mechanisms underlying ER-export and post-ER trafficking remained intact in the absence of TMED9, and that TMED9 is a key component of the machinery that is specifically required for the ER-export of misfolded GPI-APS.

### Chemical inhibitor of TMED9, BRD4780, inhibits RESET of misfolded GPI-APs

The siTMED9 depletion studies strongly suggest that TMED9 expression is required for RESET. However, TMED9-depletion by knock down or knock out was reported to be associated with a decrease in the levels of other p24/TMED-family proteins, including TMP21 and TMED2 [34, 35]. We replicated this phenomenon here with siTMED9-treated YFP-PrP* NRK cells (Figure 4A-B). As expected, siTMED9-depletion reduced TMED9 expression levels by 91.34% +/- 1.54% (Stdev.p, n=3). However, siTMED9-depletion also resulted in a 62.18±22.95% (Stdev.p, n=3) reduction in TMP21 and 48.26±16.89% (Stdev.p, n=3) reduction in TMED2 levels. The concomitant reduction of known RESET factors, TMP21 and TMED2, exposes the possibility that TMED9’s involvement in RESET pathway may be indirect through the stabilization of TMP21 or TMED2. Therefore, to gain insight into whether TMED9 plays a direct role in RESET, we opted to use the recently published chemical inhibitor of TMED9, BRD4780, that has been shown to acutely and specifically destabilize TMED9 [22].

**Figure 4:**
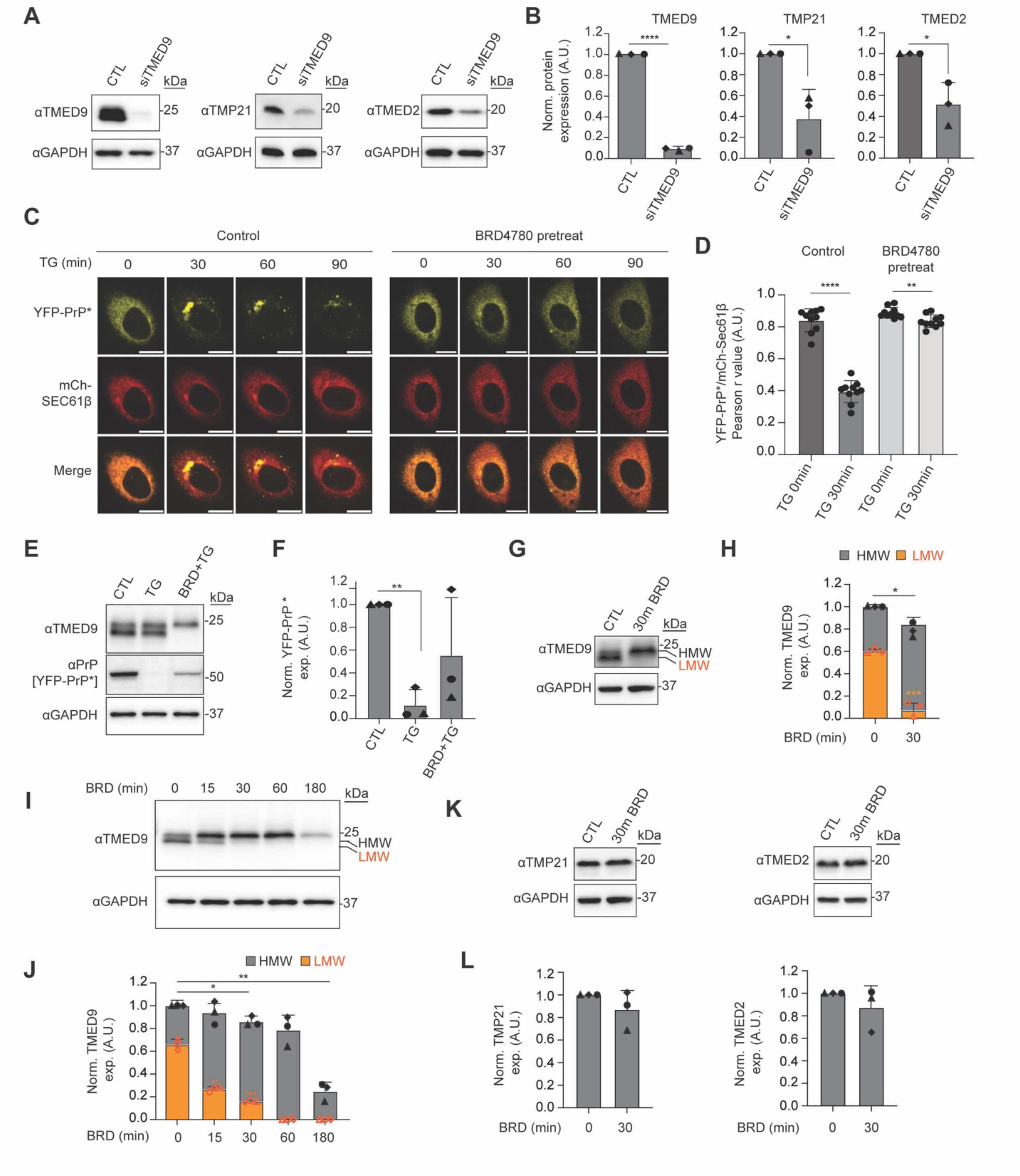
BRD4780 modifies TMED9 and inhibits RESET of misfolded GPI-APs. **(A)** Representative western blots of either non-targeting siRNA (CTL) or TMED9 siRNA (siTMED9)-treated YFP-PrP* NRK cells. Membranes were probed for TMP21 and GAPDH or TMED2 and GAPDH. (n=3 biological replicates). **(B)** Bar graph representing the mean band intensity for TMED9, TMP21 or TMED2 from 3 independently performed experiments as shown in (A). For each condition, TMED9. TMP21 or TMED2 band intensities were double normalized: first against the band intensity of GAPDH, second against the CTL band intensity. Error bars represent the standard deviation of the mean. Symbols were coded for each independently performed experiment. Statistics were calculated from unpaired t-test with Welch’s correction with * indicated p<0.05. **(C)** Time-lapse images of YFP-PrP* NRK cells that were co-transfected with mCh-Sec61β. Cells were either not pretreated (Control) or pretreated with BRD4780 (BRD pretreat) for 30 min. Image collection was started immediately after the addition of thapsigargin (TG). Scale bars represent 10 µm. **(D)** Plot of the average Pearson’s r values between YFP-PrP* and mCh-Sec61β. For Control vs. BRD4780-pretreated conditions, as described in (C), 10 time-lapses of individual cells were analyzed for the 0 and 30 min time points. Pearson’s colocalization coefficients, r, were measured between YFP-PrP* and mCh-Sec61β within the boundaries of the cell. For each data point, the boundaries of each cell at 0 and 30min time points were revealed by temporarily maximizing the gain for YFP-PrP*. **(E)** Representative western blots of YFP-PrP* NRK cells that were untreated (CTL), TG-treated for 90 min (TG) or BRD4780 (30 min)-pretreated and harvested after 90 min of TG treatment (BRD+TG) (n=3 biological replicates). Blots were probed for TMED9, PrP (YFP-PrP*) and GAPDH. **(F)** Average of normalized YFP-PrP* band intensity from 3 independently performed experiments as shown in (E). For each condition, YFP-PrP* band intensities were double normalized. First, they were normalized against the band intensity of GAPDH. Second, they were normalized against the CTL band intensity. Error bars represent standard deviations of the mean. Statistics were calculated from unpaired t-test with Welch’s correction with *** indicated p<0.001. **(G)** Representative western blots of YFP-PrP* NRK cells that were either untreated control (CTL) or treated with BRD4780 for 30 min (30’ BRD). Blots were probed for TMED9, TMP21, and TMED2 and their respective GAPDH (n=3 biological replicates). TMED9 protein profile shows 2 distinct bands, one lower band (orange arrow, LMW) and a higher band (black arrow, HMW). **(H)** Average of normalized High Molecular Weight (HMW) and Low Molecular Weight (LMW) TMED9 band intensity from 3 biological replicates as shown in (G). For each condition, TMED9 band intensities were double normalized. First, they were normalized against the band intensity of GAPDH. Second, they were normalized against the CTL intensity of each protein probed. Orange bars represent the quantification of the lower molecular weight (LMW) band intensity and gray bars represent the higher molecular weight (HMW) band intensity. Error bars represent standard deviations of the mean. Symbols were coded for each independently performed experiment. Statistics were calculated from unpaired t-test with Welch’s correction with * indicated p<0.05, and *** p<0.001. **(I)** Representative western blots of endogenous TMED9 from YFP-PrP* NRK cells after 180 min of BRD-treatment (n=3 biological replicates). Blots were probed for TMED9 and GAPDH. TMED9 protein profile shows 2 distinct bands, one LMW (orange arrow) and a HMW (black arrow). **(J)** Average of normalized HMW and LMW TMED9 band intensity from 3 biological replicates as shown i (I). For each time point TMED9 band intensities were double normalized: first by the band intensity of GAPDH and second against the BRD4780 0 min band intensity. Orange bars representing the measurement for the LMW of TMED9 and gray, the HMW. Error bars represent standard deviations of the mean. Symbols were coded for each independently performed experiment. Statistics were calculated from unpaired t-test with Welch’s correction with * indicated p<0.05 and ** p<0.01. **(K)** Representative western blots of TMP21, TMED2 and their representatives GAPDH either control (CTL) or BRD4780 (30 min)-treated YFP-PrP* NRK cells (n=3 biological replicates). **(L)** Average of normalized TMP21 and TMED2 band intensities from 3 biological replicates as shown in (K). For each condition, TMP21 or TMED2 band intensities were double normalized: first against the band intensity of GAPDH, second against the CTL band intensity. Error bar represents standard deviation of the mean. Symbols were coded for each independently performed experiment. Statistics were calculated from unpaired t-test with Welch’s correction with * indicated p<0.05.

To determine the optimal parameters for acute TMED9 inhibition by BRD4780, we used live cell imaging assays to identify the shortest pretreatment time and lowest concentration for BRD4780 that would inhibit TG-induced RESET in 100% of the YFP-PrP* NRK cells. To mark the ER, we co-transfected our YFP-PrP* NRK cells with the previously characterized ER-marker mCherry-Sec61β [36, 37]. We found that a 30 min pre-treatment of 100 µM BRD4780 was sufficient to completely block TG-induced RESET in 100% (n=34) of the cells. Instead of relocalizing to downstream secretory pathway compartments, YFP-PrP* remained in the ER of the YFP-PrP* NRK cells for at least 90 min after the addition of TG in BRD4780-pretreated cells (Figure 4C). Pearson’s colocalization analysis of YFP-PrP* with the ER marker, mCherry-Sec61β, revealed strong colocalization at steady-state that was reduced upon TG-treatment in control cells, but maintained in BRD4780-pretreated cells (Figure 4D). Similar experiments conducted with GFP-CD59 (C94S) NRK cells revealed that in 100% of the cells that were treated with TG alone, the ER-localized population of GFP-CD59 (C94S) underwent RESET (n=33), while in BRD4780-pretreated cells, the ER-population of GFP-CD59 (C94S) remained trapped in the ER in 100% of the cells (n=32) (Supp Figure S4A). As with YFP-PrP* (Figure 4D), colocalization analysis of GFP-CD59 (C94S) with mCherry-Sec61β supported the observation that GFP-CD59 (C94S) underwent TG-induced ER-export in the control cells, but remained in the ER of the BRD4780-pretreated cells (Supp Figure S4B).

Although BRD4780 pretreatment appeared to completely inhibit RESET, it did not prevent the loss of YFP-PrP*. After BRD4780 pretreatment, addition of TG induced a gradual fading of the YFP-fluorescence over the 90 min time course, without any obvious change in the localization (Figure 4C). This suggested that the YFP-PrP* was degraded from the ER by an alternate mechanism. Western blot analysis produced results that were consistent with the live-cell imaging. While TG alone induced degradation of YFP-PrP* within 90 min, the 30 min BRD4780-pretreatment partially inhibited this TG-induced clearance (Figure 4E-F). Altogether, our results demonstrate that although misfolded GPI-APs appear to get degraded by an alternate pathway, BRD4780 completely blocks the TG-induced clearance of YFP-PrP* via the RESET pathway.

Next, we sought to test whether the BRD4780-block on RESET was due to degradation of TMED9 in NRK cells. In untreated and TG-treated cells, our western blot results for TMED9 revealed a doublet of approximately 23.5 and 24 KDa just under the 25 KDa molecular weight marker (Figure 4E). For the cells pre-treated for BRD4780 for 30 min prior to the addition of TG for 90 min (BRD + TG), we observed that the lower molecular weight (LMW) band of the doublet disappeared (Figure 4E). To gain insight into the fate of the LMW band of TMED9, we treated cells with BRD4780 alone for 30 min and observed the overall expression level of TMED9 did not change (Figure 4G-H). Instead, there was a clear mobility shift from the approximately 23.5 KDa LMW band to the approximately 24 KDa higher molecular weight (HMW) band (Figure 4G-H). This mobility shift in TMED9 within 30 min of BRD4780-treatment suggested that the 30 min BRD4780-pretreatment induced block on RESET shown in Figure 4C was not due to degradation of TMED9. Instead the 30 min BRD4780-treatment appeared to acutely change the physical state of TMED9.

To gain more insight into BRD4780’s effect on TMED9, we performed a BRD4780-treatment time course and probed for TMED9 by western blot. We observed the gradual loss of the LMW band from 0 to 30 min. After 30 min of BRD4780-treatment, the LMW band decreased by 75.70±1.24 (Stdev.p, n=3) (Figure 4I-J), which corresponds with a complete inhibition of RESET (Figure 4C). By 60 min there was a complete loss of the LMW band, followed by an eventual loss of the HMW band over 180 min of BRD4780-treatment. Although we cannot rule out changes to TMP21 or TMED2 that are undetectable by western blot, we show that within 30 min of BRD4780-treatment when the LMW TMED9 band disappeared and RESET was blocked, TMP21 and TMED2 demonstrated no significant change in expression levels by western blot (Figure 4K-L). Thus, BRD4780 provides a useful tool to inhibit TMED9’s function as a RESET factor, while leaving the previously established RESET factors, TMP21 and TMED2, intact.

The results from these live-cell imaging and western blot based BRD4780 experiments are consistent with the idea that TMED9 plays an essential role in the clearance of YFP-PrP* via RESET. The rapid 30 min BRD4780-induced blockade of RESET, BRD4780-induced molecular weight shift of TMED9 and unaltered TMP21 and TMED2 bands spotlight TMED9 as RESET factor that plays a unique role beyond simply stabilizing the essential RESET factors, TMP21 and TMED2.

### BRD4780 alters TMED9 trafficking and glycosylation

The BRD4780-treatment experiments uncovered a connection between the loss of the LMW band of TMED9 and the loss of the cell’s ability to carry out RESET (Figure 4C, G-J). Intriguingly, the LMW band disappeared within 30 min of adding BRD4780, while the HMW band persisted for up to 180 min. We considered two explanations for this observation. First, the LMW band may be relatively unstable in comparison to the HMW band, and thus degrade more quickly upon BRD4780-treatment. Second, the LMW band may be converted to the HMW band as a step in TMED9’s path toward its eventual degradation. A major clue was presented in Figure 4I. Between 0 and 15 min of BRD4780-treatment, there was an evident and reproducible shift between the balance of the LMW and HMW TMED9 bands that comprised the total pool of TMED9 (Figure 4I). At time 0, total TMED9 was made up of 64.83±3.92% (Stdev.p, n=3) LMW and 34.17±3.92% (Stdev.p, n=3) HMW pools. However, within 15 min of BRD4780-treatment, the total TMED9 pool was comprised of 28.60±1.13% (Stdev.p, n=3) in the LMW and 71.40±1.13% (Stdev.p, n=3) in the HMW band. Given that the total levels of TMED9 remain steady over the first 15 min, this flip in the relative distribution of the TMED9 LMW and HMW populations negates the idea that the HMW band simply persists longer than the LMW band. Instead, this flip is consistent with the idea that the LMW population converts to the HMW population as TMED9 traffics along the BRD4780-induced degradation pathway for near complete degradation by 180 min (Figure 4I-J).

To delineate the BRD4780-induced degradation pathway for TMED9, we started by assessing which of the two major cellular degradatory systems, lysosomes and proteasomes, are involved. Western blot analysis revealed that in cells treated with BRD4780 for 180 min, there was a reduction in total TMED9 levels (Figure 5A-B, compare lanes 1 and 2). Pretreatment with the lysosomal degradation inhibitor, bafilomycin A1 (BafA1 + BRD4780) or with the proteasomal degradation inhibitor, MG132 (MG132 + BRD4780) revealed that bafilomycin A1 inhibited BRD4780-induced degradation of TMED9 HMW species (Figure 5A-B, compare lanes 2 and 3), while MG132 did not (Figure 5A-B, compare lanes 2 and 3). This suggests that BRD4780 induces lysosomal degradation of TMED9. We next assessed which of the two major categories of trafficking pathways between ER and lysosomes are involved: (1) the LC3-II-dependent ER-to-lysosome associated degradation pathways, including autophagy, that are inhibited by 3-methyladenine (3MA) [38], or (2) the ER-export to Golgi-dependent pathways that are inhibited with brefeldin A (BFA) [39–41]. We found that while 3-methyladenine did not affect BRD4780-induced degradation (Figure 5A-B, compare lanes 2 and 5), brefeldin A strongly inhibited BRD4780-induced degradation of both the LMW and HMW bands (Figure 5A-B, compare lanes 2 and 6). Brefeldin A causes the entire Golgi to fuse into the ER within 10 min [42]. Therefore, stabilization of both LMW and HMW bands by co-incubating cells with brefeldin A + BRD4780 indicates that the entire population of TMED9 must traffic through the Golgi prior to degradation. Note that for this drug panel bafilomycin A1 (BAF), MG132 and brefeldin A (BFA) stocks were each prepared in DMSO and diluted 1:4000, 1:2000 and 1:1000, respectively, into the medium. BRD4780 and 3-MA stocks were each prepared in water or directly included in the medium, as explained in the Materials and Methods. However for the drug panel in Figure 5A-B, separate DMSO controls for the BAF+BRD, MG132+BRD or BFA+BRD treatments were not included to simplify the experiment without changing our interpretation of the results.

**Figure 5:**
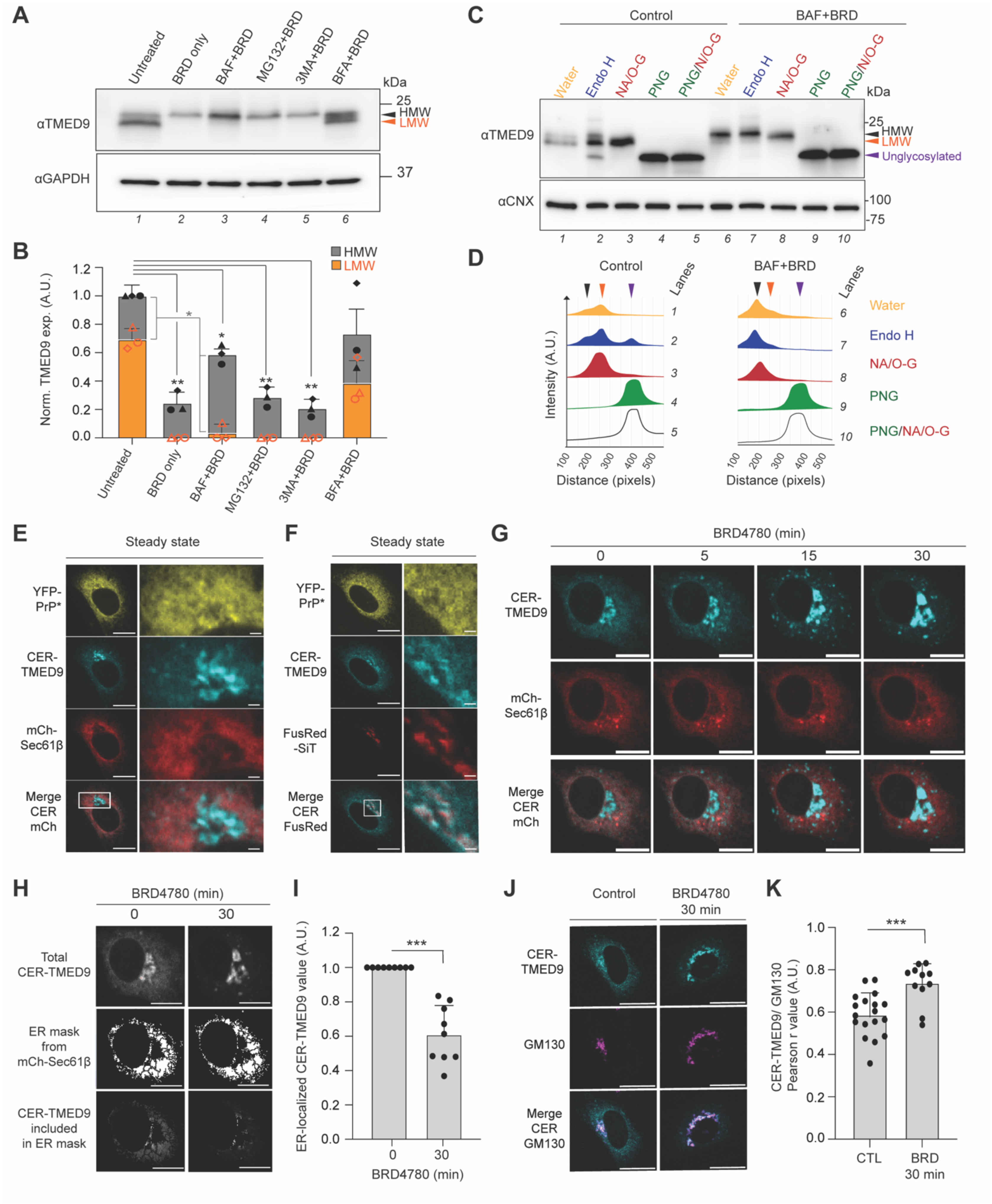
BRD4780 alters TMED9 trafficking and glycosylation. (A) Representative western blots of YFP-PrP* NRK cells that were treated as listed below (n=3 biological replicates). Untreated, treated with BRD4780 for 180 min (BRD), pretreated with bafilomycin A1 (BAF) for 180 min and co-treated with BRD for 180 min (BAF+BRD), pretreated with MG132 for 180 min and co-treated with BRD for 180 min (MG132+BRD), pretreated with 3-methyladenine (3MA) for 180 min and co-treated with BRD for 180 min (3MA+BRD) or pretreated with BFA for 180 min and co-treated with BRD for 180 min (BFA+BRD). Bafilomycin A1 (BAF), MG132 and brefeldin A (BFA) stocks were each prepared in DMSO and diluted 1:4000, 1:2000 and 1:1000 into the medium. For this panel, separate DMSO controls for the BAF+BRD, MG132+BRD or BFA+BRD treatments were not included. Blots are probed for endogenous TMED9 and GAPDH. **(B)** Bar graph representing the mean band intensity for TMED9 from 3 biological replicates as shown in (A). For each condition, TMED9 band intensities were double normalized first by the band intensity of GAPDH and second within individual experiments using control band intensity. Orange bars represent the measurement for the low molecular weight (LMW) band of TMED9 and gray bars represent the measurement of the high molecular weight (HMW) band of TMED9. Error bars represent standard deviations of the mean. Symbols were coded for individual experiments. Statistics were calculated from unpaired t-test with Welch’s correction with * indicated p<0.05 and ** p<0.01. **(C)** Western blots of digested protein lysates of stably transfected YFP-PrP* NRK cells either in control condition (Control) or after a pretreatment of 2 h of bafilomycin followed by co-incubation with 2 h of BRD4780 (BAF+BRD). Protein lysates undigested (water) or digested with Endoglycosidase H (Endo H), Neuraminidase+O-Glycosidase (NA+O-Glyc.), Peptide-N-glycosidase F (PNG) or NA+O-Glyc+PNGase (NA+O-G+PNG) for 3 h at 37°C (n=1 experiment). Blots were probed for TMED9 and calnexin (CNX). **(D)** Western blot band graphics obtained by first analyzing the plot lanes and second generating the line graphs of pre-selected regions of interest (ROI) from the western blot for control samples (left) or BAF+BRD samples (right). Black arrow represents the HMW form, orange arrow represents the LMW and purple, the unglycosylated form of TMED9. **(E)** (left) Fluorescence image of a typical untreated YFP-PrP* NRK cell that was co-transfected with CER-TMED9 and ER-marker, mCherry-Sec61β. Scale bar represents 10 µm. (right) Magnified panel of the stroked region on the right. Scale bar represents 1 µm. **(F)** (left) Fluorescence image of a typical untreated YFP-PrP* NRK cell that was co-transfected with CER-TMED9 and Golgi marker, FusionRed-SiT. Scale bar represents 10 µm. (right) Magnified panel of the stroked region on the right. Scale bar represents 1 µm. **(G)** Time-lapse imaging sequence of a NRK cell that was co-transfected with CER-TMED9 and ER marker, mCherry-Sec61β. Image-collection was started immediately after the addition of 100 µM BRD4780 treatment. Scale bar is 10 µm. **(H)** Example of an ER mask created from the ER-marker, mCherry-Sec61β. The ER mask was used to measure the CER-TMED9 fluorescence intensity within the ER at 0 or 30 min time points after the addition of BRD4780 from 9 individual time-lapses as shown in (G) and quantified in (I). **(I)** Bar graph representing the mean of ER-localized CER-TMED9 fluorescence-intensity measurements of 9 individual time-lapses of CER-TMED9 and mCherry-Sec61β-expressing cells, as shown in (G). The ER mask was generated based on and as analyzed in in (H). Statistics were calculated from unpaired t-test with Welch’s correction with ** indicated p<0.01 **(J)** Representative immunofluorescence images of YFP-PrP* NRK cells that were transfected with CER-TMED9 under control untreated (CTL) or 100 µM BRD4780 (30min)-treated (BRD) conditions and stained for GM130. Scale bar is 20 µm. **(K)** Bar graph representing the mean of Pearson’s correlation coefficient r to measure colocalization between CER-TMED9 and GM130 under untreated (CTL) and BRD4780 (30 min)-treated (BRD) conditions, as exemplified in (J). Colocalization for 18 cells were measured for control conditions and 11 cells were measured for BRD4780 (30min)-treated cells. The r values between CER-TMED9 and GM130 for 0 min and 30 min time points were 0.589 ± 0.099 and 0.739 ± 0.086, respectively. Statistics were calculated from unpaired t-test Welch’s correction with *** indicating p<0.001.

A possible explanation for the BRD4780-induced shift in TMED9’s molecular weight is that it reflects the maturation of glycosylation as TMED9 moves from the ER and ERGIC through the Golgi. In eukaryotic cells, the organization of glycosyltransferases across the ER and Golgi allow for predictable sequential glycosylation reactions to occur as secretory pathway proteins traffic through these compartments. Thus, glycosidase digests are a standard procedure to monitor glycoprotein movement through the secretory pathway [43, 44]. Upon entry into the ER, most secretory pathway proteins are N-linked glycosylated [45]. PNGase F cleaves all N-linked glycosylations. As glycoproteins move from the cis to the medial cisternae, they encounter Golgi-resident glycosyltransferases that modify the existing N-linked glycosylations to Endo H resistant forms [43, 44]. As they move through medial and trans cisternae, they may encounter O-linked glycosyltransferases and sialyltransferases, which conjugate sialic acids to either N- or O-linked glycosylations [43, 46, 47]. With this in mind, we hypothesized that the steady-state pool of LMW and HMW TMED9 bands may be a reflection of different glycosylation-states of TMED9 whose ratios are affected by BRD4780-induced of TMED9 ER-to-Golgi transport. We therefore took an open approach to test our hypothesis with a panel of glycosidase digests.

PNGase F and Endo H digests of the YFP-PrP* NRK lysates confirmed that at steady-state TMED9 is N-linked glycosylated with Endo H resistant and sensitive populations (Figure 5C-D, Supp Figure S5A-B). The Endo H resistant populations aligned with the LMW and HMW bands, and the Endo H sensitive population aligned with the PNGase F digested TMED9 core protein. The neuraminidase + O-glycosidase digest appeared to collapse the HMW and LMW doublet into single band with a similar mobility to the LMW band, indicating that the HMW band corresponded with the subset of TMED9 that accessed the trans cisternae. After BRD4780-treatment for 2 hrs in cells that were pretreated with bafilomycin to prevent TMED9 degradation, the entire TMED9 population ran as a single HMW band that was entirely Endo H resistant and neuraminidase + O-glycosidase sensitive (Figure 5C-D). The following two results suggest that TMED9 does not get O-linked glycosylated, but instead its N-linked glycosylation gets sialylated: (1) PNGase F and PNGase F + neuraminidase + O-glycosidase digests each produced a single band for TMED9 that ran at the exact same molecular weight as the core TMED9 protein (Figure 5C-D), and (2) neuraminidase and neuraminidase + O-glycosidase each produced a single band for TMED9 that ran at the exact same molecular weight as LMW TMED9 in untreated and BRD4780 (30 min)-treated cells (Supp Figure S5A-B). Our results match previously published reports that at steady-state TMED9 obtains an N-linked glycosylation that can be modified with Endo H resistance and neuraminidase sensitivity [17, 48, 49], but does not get O-linked glycosylated [17]. Taken together, these glycosidase digests demonstrate that TMED9 localization shifts from the ER to the Golgi upon BRD4780-treatment in YFP-PrP* NRK cells.

To test whether BRD4780 induces a shift in localization of TMED9 from the ER to the Golgi, we performed live-cell imaging in YFP-PrP* NRK cells transfected with CER-TMED9. This allowed us to image and measure CER-TMED9 distribution before and after BRD4780-treatment in the same cell. We co-transfected the cells with either the ER marker, mCherry-Sec61β or the Golgi marker, FusionRed-SiT. At steady-state, the CER-TMED9 appeared to colocalize with the ER marker, mCherry-Sec61β (Figure 5E) and the Golgi marker, FusionRed-SiT (Figure 5F). Upon BRD4780-treatment, the ER-localized population of CER-TMED9 appeared to move out of the ER and through the Golgi (Figure 5G-K). In YFP-PrP* NRK cells that were co-transfected with CER-TMED9 and mCherry-Sec61β, CER-TMED9 visibly moved out of the mCherry-Sec61β marked ER (Figure 5G). For the 0 and 30 min timepoints of these time-courses, we quantified the decrease of cumulative CER-TMED9 fluorescence intensity in the ER by establishing an ER mask by segmentation of the mCherry-Sec61β image (Figure 5H), and measuring the cumulative, background-subtracted pixel intensity of CER-TMED9 within the ER (Figure 5I). BRD4780 (30 min)-treatment induced a 55.17±14.26% (Stdev.p, n=5) decrease in the ER pool of CER-TMED9 (Figure 5I). The decrease in the ER-pool of TMED9 appeared to correlate with an increase in the perinuclear pool of TMED9 that formed the typical pattern of the Golgi (Figure 5G). To verify a shift in TMED9-localization from the ER to the Golgi within 30 min of BRD4780- treatment, we stained control or BRD4780-treated CER-TMED9-expressing cells for the Golgi-marker, GM130, and imaged them by immunofluorescence. GM130 is a cytosolic Golgi-matrix protein that is directly targeted to the Golgi membrane [50]. BRD4780-treated cells demonstrate a visible increase in CER-TMED9 in the Golgi, and an increase in the Pearson’s colocalization coefficient r between CER-TMED9 and GM130-stained compared to control (Figure 5J-K).

The results in Figure 5 suggest that BRD4780 changes TMED9’s distribution within the secretory pathway. The glycosidase digests panel for endogenous TMED9 and live-cell imaging results using exogenously expressed CER-TMED9 each represent TMED9 populations that span across the ER, Golgi and intermediate compartments at steady-state, but shift from the ER towards the Golgi and downstream compartments within 30 min of BRD4780-treatment. Altogether, the above results demonstrate that RESET inhibition by BRD4780 is associated with alterations in TMED9 trafficking and glycosylation.

### TMED9’s glycosylation state does not impact its function in RESET

The BRD4780 (30 min)-induced inhibition of RESET coincided with conversion of TMED9 from the LMW form to the fully Endo H resistant, sialylated HMW form and relocalization of CER-TMED9 from the ER to the Golgi (Figure 5). These observations could be explained by two possible scenarios: (1) Only the LMW population of TMED9 is able to carry out RESET or (2) TMED9 must be localized to the ER and/or immediately ER-accessible compartments to carry out RESET. If the first scenario were correct, then TMED9 that was converted to HMW would not be able to support RESET. If the second scenario were correct, then TMED9 that was relocated into compartments downstream of the cis-Golgi would not be able to carry out RESET.

To test the possible explanations, we performed a series of identical incubations in two separate but identical dishes of YFP-PrP* NRK cells. One dish culminated in the collection of the lysates to examine the LMW/HMW states of TMED9. The other dish was assessed by imaging to test for the cells’ ability to carry out TG-induced RESET of YFP-PrP* based on the relocalization of the YFP-PrP* signal from ER (cytoplasm-filling reticular pattern excluding the nucleus) to Golgi (perinuclear crescents). First, we replicated our previous finding shown in Figures 3G. In untreated cells, the majority of TMED9 was in the LMW state and 100% of the cells were able to undergo RESET (Figure 6A, lane 1 and Figure 6B, panel 1). Upon a BRD4780 (30 min)-treatment, TMED9 converted to the HMW population and all of the cells lost their ability to undergo RESET (Figure 6A, compare lanes 1 and 2, and Figure 6B, panels 1 and 2). However, when we introduced a 30 min wash-out step immediately after the BRD4780 (30 min)-treatment, TMED9 remained in the HMW form but the majority of the cells (72%, n=50) regained competency to carry out RESET (Figure 6A, lane 3 and Figure 6B, panel 3). Thus acute inhibition of RESET with BRD4780-treatment is reversible, despite the glycosylation state of TMED9. As shown previously (Figure 3I-J), long-term treatment with BRD4780 for 180 min induced near total TMED9-degradation (Figure 6A, lane 4) and blocked RESET (Figure 6B, panel 4), and was not reversible upon BRD4780 wash-out (Figure 6A, lane 5, Figure 6B, panel 5).

**Figure 6:**
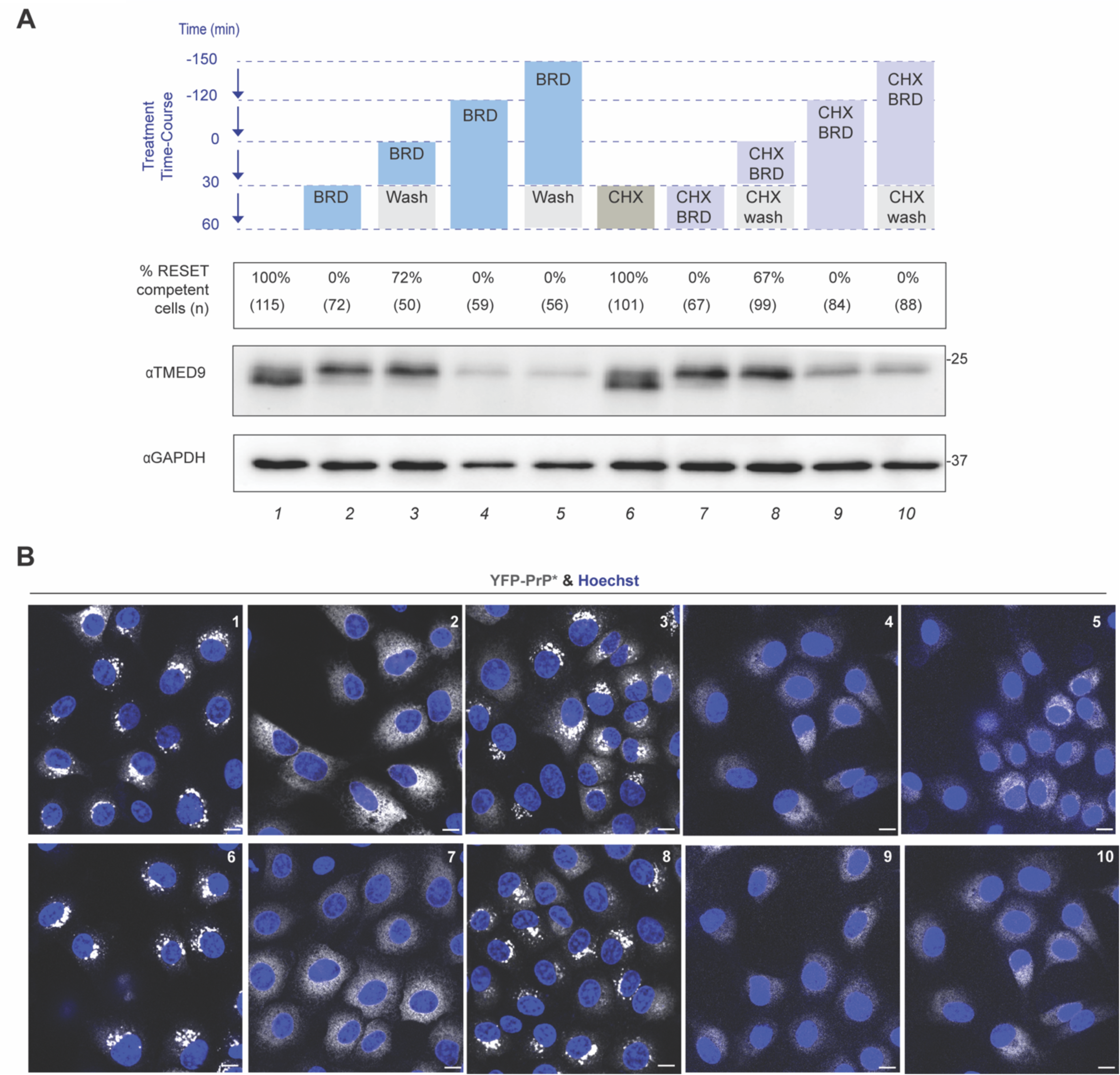
TMED9’s function in RESET is not related to its glycosylation state. **(A)** Table presenting the different treatments and their effects on TMED9’s molecular weight. As depicted in the treatment time-course, YFP-PrP* NRK cells were pre-treated with BRD4780 (BRD) or cycloheximide + BRD4780 (CHX BRD) for different periods of time with or without a 30 min BRD4780 wash-out (wash). “CHX wash” indicates that after pre-treatment with CHX + BRD, the BRD4780 wash-out included cycloheximide. The numbers below the treatment time-course diagram indicate the percentage of cells that were competent for RESET after each treatment course. The percentages of RESET competent cells were obtained by dividing the number of cells in which YFP-PrP* re-localized from the ER to the Golgi within 30 min of adding thapsigargin by the total number of cells counted. The n values (i.e. total number of cells counted per treatment course) were included directly under the percentage. After the pre-treatments, “ER” vs “Golgi” localization was assessed based on the pattern of YFP fluorescence as observed through live-cell confocal imaging. Cells were counted as RESET competent if the YFP-PrP* fluorescence pattern shifted from an ER pattern to a Golgi pattern. At steady-state, YFP-PrP* fluorescence is dispersed throughout the cell in a reticulo-tubular pattern with a sharp outline of nucleus that is consistent with ER and the nuclear envelope subdomain of the ER. During RESET, YFP-PrP* fluorescence changes from disperse to a perinuclear localization that is consistent with the Golgi. Aligned directly below the treatment courses and percentages, are the TMED9 and GAPDH western blots. Each lane encompassing the treatment, % RESET competent cells, TMED9 molecular weight, GAPDH loading control, was numbered on the bottom. There were 10 individual treatment conditions. **(B)** Representative live-cell images of YFP-PrP* NRK cells after each indicated treatment conditions (1-10). The brightness and contrast settings were increased in panels 4, 5, 9 and 10 with respect to the other panels in order to more easily assess ER vs. Golgi localization patterns. The nuclei were stained with Hoechst.

To eliminate the possibility that newly synthesized TMED9 could be contributing to the pool of TMED9 involved in RESET, we performed the above experiments in the presence of the protein translation inhibitor, cycloheximide (CHX). Again, in the presence of CHX, BRD4780 (30 min)-treatment induced the conversion of TMED9 to the HMW population and blocked RESET (Figure 6A, compare lanes 6 and 7, Figure 6B, compare panels 6 and 7). Following the CHX + BRD4780 (30 min)-co-treatment, BRD4780 wash-out using medium containing only CHX restored the cell’s ability to carry out RESET in the majority of the cells (67%, n=99), but did not alter the HMW state of TMED9 (Figure 6A, lane 8, Figure 6B, panel 8). The finding that simply washing out BRD4780 restored RESET in the presence of a protein translation inhibitor disconnects the glycosylation state of TMED9 from its ability to support RESET. Finally, near complete depletion of the existing pool of TMED9 with CHX + BRD4780 (180 min) co-treatment did not allow for the restoral of RESET with BRD4780 wash-out (Figure 6A, lanes 9 and 10, Figure 6B, panels 9 and 10).

Collectively, these experiments reiterate the requirement for TMED9’s presence to carry out RESET. These experiments demonstrate that TMED9’s glycosylation state does not impact its function in RESET.

### TMED9 must have access to misfolded GPI-APs in the ER to coordinate RESET

To gain insight into whether TMED9’s localization was involved in its ability to execute RESET, we again used exogenously expressed CER-TMED9 as a proxy for TMED9. We co-transfected NRK cells with YFP-PrP*, CER-TMED9 and mCherry-Sec61β. We co-incubated these cells with cycloheximide (CHX) + BRD4780 and then performed BRD4780 wash-out using medium with only CHX (Figure 7A). Live cell imaging revealed that CHX + BRD4780 (30min) induced export of CER-TMED9 out of the ER, as demonstrated earlier. However during the CHX-only wash-out step, CER-TMED9 appeared to move back into the ER and the cells regained their ability to carry out RESET (Figure 7A, Video 3).

**Figure 7:**
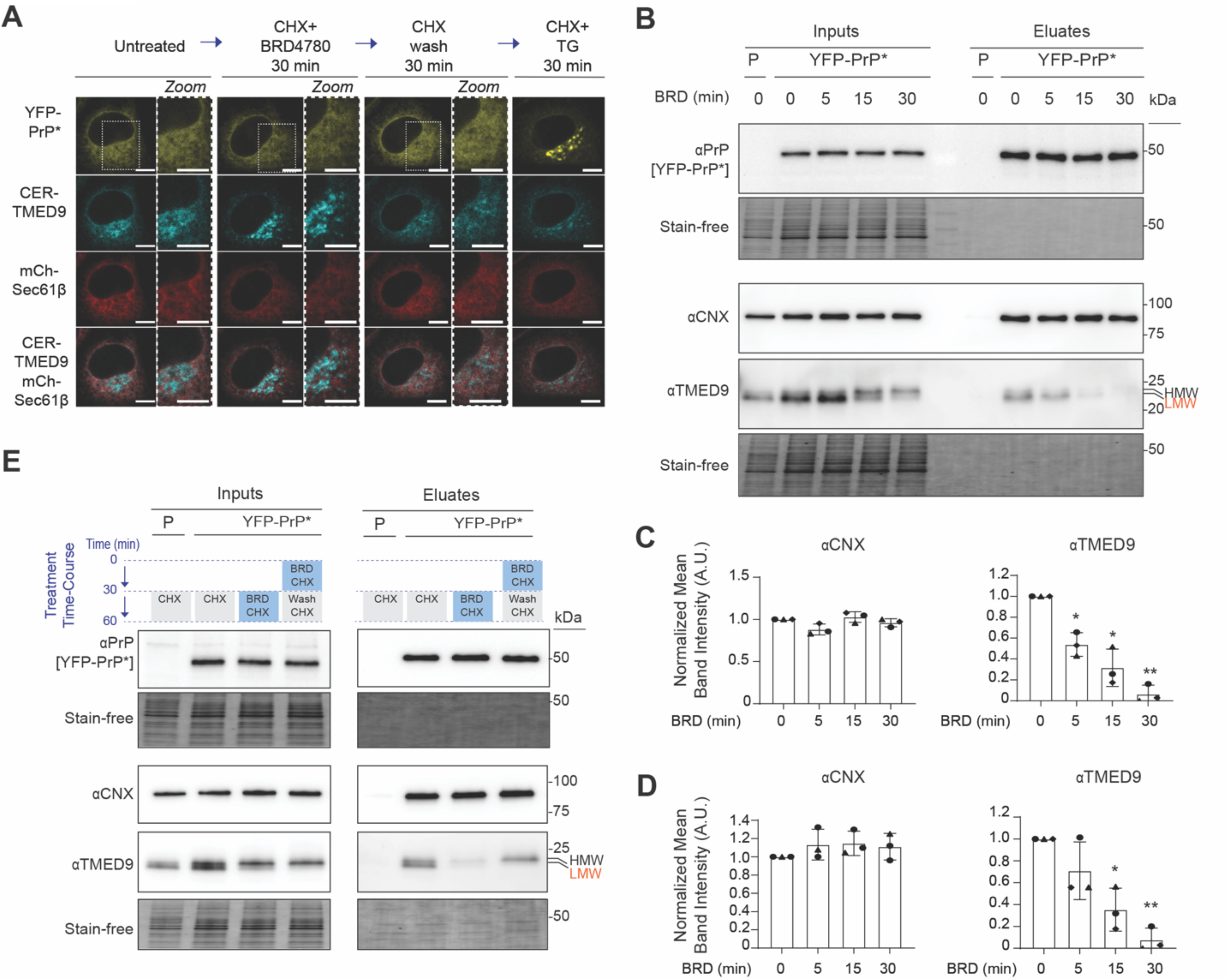
TMED9 must have access to misfolded GPI-APs in the ER to coordinate RESET. **(A)** Still frames of a typical YFP-PrP* NRK cell that was co-transfected with CER-TMED9 and mCherry-Sec61β and treated as follows. The first image is of the cell before treatment began (Untreated). The second image is of the cell after 30 min with 50 µg/ml of CHX + BRD4780 (CHX + BRD4780 30 min). The third image is of the cell 30 min after washing with CHX-only medium (CHX wash 30 min). The fourth image is of the cell 30 min after treatment with 50 µg/ml of CHX + 0.1 µM thapsigargin (CHX + TG 30 min). Juxtaposed with each image is an area zoom of the region within the stroked frame. Scale bar represents 10 µm in the larger field of view and in the magnified panel of the stroked region. These select images are also provided in Video 2. **(B)** Representative western blots of GFP-tag co-immunoprecipitations (co-IPs) from the parental untransfected NRK cells (P) or YFP-PrP* NRK cells (n=3 biological replicates). Quantified in (C). Cells were harvested at the indicated time points after BRD4780-treatment. Blots were probed for PrP to detect YFP-PrP*, or for endogenous calnexin (CNX) and TMED9. Under each western blot is depicted a “Stain-Free” image of the total protein in the gel. **(C)** Bar graph displaying the normalized mean band intensities for the CNX and TMED9 that co-eluted with YFP-PrP* from 3 biological replicates as shown in (B). For each time point, CNX and TMED9 eluate band intensities were double normalized. First, they were normalized by the band intensity of the eluted YFP-PrP* band. Second, they were normalized by the time 0 eluate band intensity for each protein probed. Error bars represent standard deviations of the mean. Symbols were coded for individual experiments. Statistics were calculated from unpaired t-test with Welch’s correction with and ** indicated p<0.01. **(D)** Bar graph displaying the normalized mean band intensities for the CNX and TMED9 that co-eluted with YFP-PrP* from 3 biological replicates as shown in (B). This graph represents the co-eluted CNX and TMED9 normalized against the amount of CNX and TMED9 in the inputs. Thus, for each time point, CNX and TMED9 eluate band intensities were triple normalized. First, they were normalized against the band intensities in the input. Second, they were normalized by the band intensity of the eluted YFP-PrP* band. Third, they were normalized by the time 0 eluate band intensity for each protein probed. Error bars represent standard deviations of the mean. Symbols were coded for individual experiments. Statistics were calculated from unpaired t-test with Welch’s correction with and ** indicated p<0.01. **(E)** Western blots of GFP-tag co-immunoprecipitations (co-IPs) from the parental untransfected NRK cells (P) or YFP-PrP* NRK cells (n=1 experiment). Cells were harvested after the indicated treatments. Blots were probed for PrP to detect YFP-PrP*, or for endogenous calnexin (CNX) and TMED9. Under each western blot is depicted a “Stain-Free” image of the total protein in the gel.

Based on our combined western blot and live-cell imaging results for siRNA depletion or BRD4780-inhibition and depletion of TMED9 (Figures 3, 4, 6, 7A, Supp Figures S2 and S3), we concluded that TMED9 expression was required for RESET, regardless of whether TMED9 was in the LMW or HMW form. The acute inhibition of RESET by BRD4780-treatment was concomitant with the relocalization of TMED9 out of the ER, while restoration of RESET was concomitant with the return of CER-TMED9 to the ER upon BRD4780 wash-out (Figures 4, 6, 7A). Importantly, we observed that YFP-PrP* did not co-traffic with CER-TMED9 out of the ER upon BRD4780-treatment (Figure 7A, Supp Figure S6). Instead, YFP-PrP* appeared to be left behind.

The above observations are consistent with the idea that in order to coordinate RESET, TMED9 requires access to the ER where RESET substrates are largely localized in association with calnexin (Supp Figure S1 and Figure 2, [12, 14]). To directly test this idea, we performed a co-immunoprecipitation time course, pulling down YFP-PrP* from untreated or BRD4780-treated YFP-PrP* NRK cells. In untreated cells, YFP-PrP* co-purified with calnexin and TMED9 (Figure 7B, t=0). However, upon BRD4780-treatment, we observed the rapid dissociation of TMED9 from YFP-PrP* between 5 and 30 min (Figure 7B). We quantified the results by two different methods (Figure 7C-D). In Figure 7C, we double normalized mean band intensities for the calnexin and TMED9 that co-eluted with YFP-PrP* from 3 biological replicates, as shown in Figure 7B, first by the band intensity of the eluted YFP-PrP* band and second by the time 0 eluate band intensity for each protein probed. This provided a measurement of the change in how much calnexin or TMED9 co-immunoprecipitated with YFP-PrP* over the BRD4780- treatment time course. In Figure 7D, we triple normalized the same data by additionally taking into account the amount of starting material in the inputs. First the eluate bands were normalized against the band intensities in the input. Second, they were normalized by the band intensity of the eluted YFP-PrP* band. Third, they were normalized by the time 0 eluate band intensity for each protein probed. Regardless of the quantification scheme, the data revealed that upon BRD4780-treatment, TMED9 nearly completely dissociated from YFP-PrP* (Figure 7C-D). Additionally, TMP21 and TMED2 appeared to dissociate from YFP-PrP* within the same time frame (Supp Figure S7). Intriguingly, YFP-PrP* steadily maintained association with calnexin, even while the p24-family members rapidly disengaged (Figure 7B-D), which is consistent with the imaging results showing that YFP-PrP* remained in the ER upon BRD4780- treatment (Figure 7A, Supp Figure S6).

Finally, to test our hypothesis that BRD4780 wash-out restores RESET by allowing TMED9 to return to the ER where it can interact with the RESET substrate, we performed co-immunoprecipitations of YFP-PrP* with anti-GFP antibody conjugated beads under the BRD4780-treatment and wash-out conditions (Figure 7D). We prevented the involvement of newly synthesized TMED9 by including cycloheximide (CHX) in all of the incubations. As a negative control for the co-immunoprecipitation, we included the parental cells with CHX. We treated YFP-PrP* NRK cells as follows: (1) CHX for 30 min, (2) CHX + BRD4780 for 30 min, and (3) CHX + BRD4780 for 30 min followed by a 30 min wash-out in medium containing only CHX. Again, we observed the TMED9 band shift from LMW to HMW upon the CHX + BRD4780 (30 min)-treatment in the input lanes. In the eluate lanes, the CHX (30 min) treatment allowed YFP-PrP* to pull down both calnexin and TMED9, while the CHX + BRD4780 (30 min)-treatment allowed YFP-PrP* to pull down calnexin but not TMED9. Excitingly, in the eluate lane for CHX + BRD4780 (30 min) followed by the CHX-only wash-out, pull-down of YFP-PrP* co-eluted calnexin and TMED9. After the wash-out, the TMED9 that co-eluted with YFP-PrP* ran at the higher molecular weight, indicating that it had trafficked to downstream compartments of the Golgi and returned to the ER where it could associate with YFP-PrP*.

Taken together, our results demonstrate that TMED9’s location in the ER is essential for it to carry out RESET. We showed that TMED9 relocalizes from the ER to the Golgi upon BRD4780 (30 min)-treatment. Upon relocalization out of the ER, TMED9 loses contact with the RESET substrate and its ability to conduct RESET. However if TMED9 is restored to the ER upon BRD4780 wash-out, then TMED9 is able to associate with the RESET substrate and perform its essential function in the RESET pathway. These results presented here are consistent with a model in which TMED9, in conjunction with other p24-family members, is a key constituent of the essential ER-export machinery that captures misfolded GPI-APs released by calnexin and conveys them from the ER to the Golgi (Supp Figure S8).

## Discussion

TMED9, along with its p24-family members, has been shown to associate with COPII and COPI coat proteins and constitutively traffic between the ER and Golgi within the early secretory pathway of yeast, plant and mammalian cells [16, 17, 51–54], independently of cargo-binding [55]. To elucidate TMED9’s role in RESET, we exploited acute thapsigargin-treatment as a useful tool to synchronize traffic through the RESET pathway [12, 14]. Within the first 20 min after thapsigargin-treatment, during the time that the misfolded GPI-APs were undergoing ER-export, they increased their association with TMED9 (Figures 1 and 2). Within 20 to 40 min after thapsigargin-treatment, misfolded GPI-APs dissociated from TMED9 as they departed from the Golgi *en masse* (Figures 1 and 2). Intriguingly, despite these dramatic changes in misfolded GPI-AP-TMED9 interactions during RESET, there were no obvious changes in the overall localization of CER-tagged TMED9 (Figure 1), nor in the endogenous TMED9 glycosylation patterns, which would signify a change in the balance of ER vs. Golgi-pools (Figure 2, input lanes), nor in the endogenous TMED9 immunofluorescence staining (Figure 3H, compare 0 min and 90 min CTL images). These observations suggest that TMED9 trafficking remained unperturbed despite the thapsigargin-induced flux of misfolded GPI-APs through RESET. This fits with the previously proposed idea that p24-family cycling is normally not directed by cargo binding [55]. However, more comprehensive studies examining TMED9’s molecular movements within the ER, Golgi and connecting compartments would be required to uncover the dynamics that may be hidden within this overall stasis.

To explain the data generated in this study in the context of the greater p24 literature, we propose the following model (Supp. Figure S8). In our model, TMED9 and associated p24-family members constitutively cycle between the ER and the Golgi (Supp. Figure S8A). These p24-family members include the demonstrated RESET factors TMP21 [12], TMED2 [20], and TMED9. Upon release from calnexin, the misfolded GPI-anchored proteins piggy-back on the cycling p24-family members to gain access to the Golgi (Supp. Figure S8B). Once in the Golgi, misfolded GPI-APs dissociate from the p24-family members and continue to move forward to downstream compartments of the endomembrane system. When TMED9 and its associated p24-family members are absent from the ER, misfolded GPI-APs are unable to access the ER-to-Golgi export sites. Instead, they remain associated with calnexin in the ER (Supp. Figure S8C). Manipulations with BRD4780 support this model presented in Supp. Figure S8 that TMED9, through its ER-Golgi-ER cycling, plays an essential role in forming and maintaining the RESET pathway for ER-export of misfolded GPI-APs. If TMED9 is removed from the ER, this pathway collapses (Supp. Figure S8C). By contrast, properly folded GPI-APs are able to leave the ER regardless of the presence or absence of TMED9 and associated p24-family members in the ER (Supp. Figure S8D). Since the exact relationships between WT GPI-APs and p24-family members remain to be dissected, we did not include the p24-family in the diagram in Supp. Figure S8D. Instead, we discuss the literature demonstrating that p24-family members bind with and influence the kinetics of WT GPI-APs ER-export [12, 16, 19, 20, 34, 56–62], and we speculate on their relationship below. Additional unknown RESET-specific and general ER-export factors remain to be identified (Supp. Figure S8).

A notable development from this and previous studies is that properly folded GPI-APs are able to undergo ER-export in the absence of TMED9 or TMP21 (Figure 3J and Supp. Figure S3F-H [12, 20]). This reinforces the idea that properly folded GPI-APs are able to access alternate ER-export pathways from misfolded GPI-APs [12]. We detected a small increase in r value between YFP-PrP WT and calnexin in siTMED9-treated cells over CTL cells (Figure 3K). This may indicate a delay in ER-export of GPI-APs to the plasma membrane as has been described previously in yeast where single p24-family members were depleted and mammalian cells where TMP21, TMED2 or TMED5 were individually depleted [16, 19, 58, 60–62]. Relatedly, a robust body of work demonstrates the protein-protein interactions between WT GPI-APs and p24-family members in yeast, plant and mammalian cells [12, 20, 34, 56–60]. This likely implicates p24-family members in direct and indirect roles for GPI-AP PQC, processing and trafficking beyond RESET. Indeed, p24-family members were shown to capture WT GPI-APs with incompletely processed GPI-anchors in the Golgi for retrieval back to the ER in yeast [56], influence the formation of membrane microdomains for WT GPI-AP ER-export sites [58, 63, 64], or facilitate trafficking of other cargo receptors or lipids between the ER and Golgi such as SURF4 or ERGIC53 that influence ER-exit site formation [54, 65–67]. Nonetheless, this and other demonstrations that WT GPI-APs are ultimately able to access the plasma membrane despite TMED9- or TMP21-depletion warrants future studies to discriminate between ER-export mechanisms for misfolded vs. properly folded secretory pathway substrates, which may open the door to targeted approaches in manipulating potentially harmful misfolded GPI-APs.

A useful insight revealed by this study is the facility of BRD4780 as a tool to manipulate and dissect the relationships between TMED9 and early secretory pathway traffic. As would be expected for the recently discovered specific inhibitor of TMED9 [22, 68], our results demonstrated that BRD4780-treatment phenocopied siTMED9 with respect to blocking RESET of misfolded GPI-APs. However, we uncovered the utility of BRD4780-treatment as a reversible modulator of TMED9. BRD4780-treatment induced the rapid re-localization of TMED9 within 15m that could be detected through glycosylation of endogenous TMED9 and imaging of CER-TMED9 (Figure 5). We showed that wash-out of BRD4780 within the first 30 min of treatment allowed the Golgi-localized TMED9 to be rapidly retrieved back to the ER. Critically, simply shifting TMED9 out of and back into the ER with BRD4780 and wash-out, respectively, caused TMED9 to dissociate and reassociate with RESET substrates and factors, and concomitantly disengage and re-engage in the RESET pathway (Figures 6-7, Supp. Figure S7). Extrapolating from our experience with BRD4780 as a reversible inhibitor, we propose that acute manipulations of TMED9 with BRD4780 will allow for a pinpointed investigations into TMED9’s functions, without triggering larger-scale irreversible changes to the endomembrane system.

We speculate that the BRD4780-induced relocalization of TMED9 indicates that BRD4780 imposes a reversible block on the retrograde transport of TMED9. Because TMED9 is normally constitutively and rapidly cycling between the ER and cis-Golgi, this block on retrograde transport results in a rapid relocalization of TMED9 out of the ER and into the Golgi. The observations presented here lead to many new questions for future studies. What is the mechanism by which TMED9 returns to the ER upon BRD4780 wash-out? Could BRD4780’s effect be due to a direct block in TMED9 binding to COPI coat proteins, or an indirect block of COPI recruitment by TMED9? Recently, Xiao *et al.* demonstrated that TMED9 is able to form homo-oligomers in the Golgi, and this oligomerized form of TMED9 has greater affinity for COPI than COPII [23]. This discovery invokes the possibility that BRD4780 blocks TMED9 homo-oligomerization, which in turn prevents retrograde transport of TMED9. Preventing the re-capture of TMED9 from the Golgi back to the ER would allow TMED9 to move forward from the Golgi to the trans-Golgi network. This idea fits with the results of our glycosidase assays of endogenous TMED9 and imaging experiments of CER-TMED9 in cells treatd with BRD4780. Conversely, BRD4780 wash-out would restore TMED9’s capacity to homo-oligomerize, recruit COPI and regain access to the ER where it is able to resume function in RESET. These ideas remain to be tested.

Intriguingly, upon BRD4780-induced relocalization of TMED9 from the ER to the Golgi, the misfolded GPI-APs remained in the ER in association with calnexin. This contrasted with misfolded luminal truncation mutant of mucin 1 (MUC1-fs). Upon BRD4780-treatment, MUC1-fs was shown to traffic out of the ER through the Golgi for subsequent bafilomycin-sensitive degradation [22]. Additionally, strong evidence for a direct interaction between MUC1-fs and TMED9 was recently published [23]. We speculate that the difference in the fates of the misfolded luminal MUC1-fs and misfolded membrane-bound GPI-APs may be based on their relative binding-affinities for the ER-resident chaperone, calnexin, and TMED9. In this scenario, TMED9 and calnexin may engage in a tug-of-war for PQC substrate-binding. The winner of this tug-of-war determines whether the PQC substrate remains in the ER with calnexin or cycles with TMED9. This idea remains to be tested.

A systematic effort involving a combination of biochemical protein-interaction studies, siRNA screening of the entire p24-family and associated molecules and RESET assays is required to dissect the precise composition of the TMED9-containing RESET cargo-receptor. This would build upon previous efforts to identify the critical p24-family components required for ER-export of misfolded GPI-APs [20]. A major confounding factor in knockdown or knockout screening of the p24-family members is that depletion of one p24-family member impacts the stability of other members (Figure 4A, [20, 34, 35, 69]). Getting around this issue would require the systematic identification of the co-depleted potential RESET factors and re-complementation, which is outside the scope of this study. Here, we attempted to address this issue with siTMED9 knock down experiments by including experiments using the small molecule inhibitor of TMED9, BRD4780. We show here that BRD4780 inhibits TMED9’s action in RESET in under 30 min by inducing TMED9 relocalization without impacting the stability of TMP21 or TMED2 (Figures 4-5). However, whether TMP21 and TMED2 co-traffic with TMED9 or are left behind in the ER remains to be discovered. Future efforts to elucidate the impact of BRD4780 on TMED9, TMED9-associated RESET factors, and ER-to-Golgi cycling of the p24-family members will be useful to better understand the relationships between TMED9, TMP21, TMED2 and other RESET factors and their substrates.

There are limitations to our study that must be overcome with improved tools. Although the Golgi-localized pool of TMED9 was clearly detectable by immunofluorescence, we were unable to unambiguously image the ER-pool of endogenous TMED9 using existing commercially available TMED9 antibodies. Possible explanations include that the antigen in the ER-pool of TMED9 was concealed through inter- or intra-protein interactions, the ER-pool of TMED9 was spread out sparsely along the relatively expansive surface area of the ER dropping the immunofluorescent TMED9 signal to near background levels, or the ER-pool of TMED9 was primarily localized to ER exit sites that were too closely apposed to the Golgi to distinguish from the Golgi-localized TMED9 signal using standard confocal-imaging. Here we found that inferring the localization of endogenous TMED9 using biochemical read-outs like glycosidase assays or mobility shifts on western blots predicted the same distribution between the ER and Golgi as was directly observed by live cell-imaging of transiently transfected CER-TMED9. Therefore, while our results demonstrate CER-TMED9’s utility as a proxy for the endogenous TMED9 population, the development and characterization of N-terminally fluorescent protein tagged CRISPR knock-ins will be helpful to overcome the limitations in imaging endogenous TMED9 dynamics.

Our study unveils new uncharted territories. In particular, given the advent of BRD4780 and the potential development of clinically beneficial derivatives [22, 68], this study revealed an important new direction of research: determine the fate of misfolded GPI-APs whose clearance via RESET has been blocked. As with siTMED9-treatment, BRD4780-treatment completely blocked RESET. However, the addition of TG in either siTMED9-treated or BRD4780-treated cells, results in a marked reduction in YFP-PrP* levels (Figures 3C, 3H and 4C), suggesting that YFP-PrP* could be degraded by alternate ER-localized degradation pathways. Similarly, Dvela-Levitt *et al.* previously demonstrated that BRD4780 induced the degradation of another misfolded GPI-AP protein, uromodulin (C126R), that accumulates intracellularly [22].

Blocking RESET and consequent access to the cell surface may be advantageous because cell surface populations of misfolding proteins have the opportunity to engage in various pathological activities. In the case of misfolded prion protein, cell surface populations of misfolding prion proteins have been shown to engage in (a) binding to and internalizing infectious prion aggregates into cells [70], (b) undergoing endocytosis and aggregating within the axonal endosomes of primary neurons and causing neuronal death [71], and (c) aggregating in extracellular deposits in the brain tissue of patients with prion disease [72, 73]. Preventing the extracellular access of misfolded GPI-APs by exploiting BRD4780 to reroute them for an internal route of degradation presents an exciting possible therapeutic intervention that may help ameliorate diseases, including prion diseases.

## Materials and Methods

### Cell lines

The isolation of the clonal line of Normal Rat Kidney (NRK) cells that we refer to here as parental NRK cells here was described previously [74]. YFP-PrP* NRK, YFP-PrP WT NRK, and GFP-CD59 (C94S) NRK cells are each derived from the NRK cells and their creation were described previously [12, 14]. In short, each of the cell lines were created by transiently transfecting a 10 cm dish with expression plasmids, expanding the transfected cells and selecting them for the stable integrants by using antibiotic resistance, and sorting them for by their fluorescence intensity. As described previously, YFP-PrP* NRK cells were sorted for expression of YFP-PrP* at similar levels to endogenous PrP in mouse brain lysate and YFP-PrP WT NRK and GFP-CD59 (C94S) NRK each express YFP-PrP WT and GFP-CD59 at similar levels to YFP-PrP* [12, 14].

### Plasmids

The construction of YFP-PrP WT, YFP-PrP* and GFP-CD59 (C94S) was described in detail [12]. Briefly, YFP-PrP WT includes the N-terminal prolactin signal sequence, which drives efficient translocation into the ER [25], and YFP-PrP* is derived from YFP-PrP WT, but contains a C179A mutation that disrupts the single disulfide bond in PrP. GFP-CD59 (C94S) includes the N-terminal rabbit lactase phlorizin-hydrolase “lactase” signal sequence, which drives efficient translocation into the ER [75–79].

Cerulean (CER)-TMED9 includes the N-terminal rabbit lactase phlorizin-hydrolase signal sequence and its construction was described in detail [24].

Golgi markers include CER-GalT (gift from Jennifer Lippincott-Schwartz, Addgene; plasmid# 11930 [29]) and FusionRed-SiT-15 (gift from Michael W. Davidson, Addgene; plasmid# 56133 [80]).

ER markers include mCherry (mCh)-Sec61β (gift from Jennifer Lippincott-Schwartz, Addgene; plasmid# 90994 [37]) and Cerulean (CER)-calnexin, which we constructed for this study. For CER-calnexin we used the same vector backbone as the SS(lactase)-CER-TMED9 described previously [24]. However, instead of the mature domain of TMED9, we PCR amplified the mature domain of calnexin (human) from a plasmid template obtained from DNASU (DNASU, HsCD00042644). “SS(lactase)” refers to rabbit lactase-phlorizin hydrolase signal sequence. “Mature domain” refers to the ORF that encodes the processed type I transmembrane protein after the N-terminal signal sequence has been cleaved by the signal peptidase. For assembly we used NEBuilder (NEB E5520S). The final ORF was verified to be the following through sequencing analysis with SS(lactase) in all capital letters, followed by linker and CER in lowercase, followed by calnexin mature domain in capital letters:

ATG GAG CTC TTT TGG AGT ATA GTC TTT ACT GTC CTC CTG AGT TTC TCC TGC CGG GGG TCA GAC TGG GAA TCT CTG CAG TCG ACG GTA CCG CGG GCC CGG GAT CCA CCG GTC GCC ACC atg gtg agc aag ggc gag gag ctg ttc acc ggg gtg gtg ccc atc ctg gtc gag ctg gac ggc gac gta aac ggc cac aag ttc agc gtg tcc ggc gag ggc gag ggc gat gcc acc tac ggc aag ctg acc ctg aag ttc atc tgc acc acc ggc aag ctg ccc gtg ccc tgg ccc acc ctc gtg acc acc ctg acc tgg ggc gtg cag tgc ttc gcc cgc tac ccc gac cac atg aag cag cac gac ttc ttc aag tcc gcc atg ccc gaa ggc tac gtc cag gag cgc acc atc ttc ttc aag gac gac ggc aac tac aag acc cgc gcc gag gtg aag ttc gag ggc gac acc ctg gtg aac cgc atc gag ctg aag ggc atc gac ttc aag gag gac ggc aac atc ctg ggg cac aag ctg gag tac aac gcc atc agc gac aac gtc tat atc acc gcc gac aag cag aag aac ggc atc aag gcc aac ttc aag atc cgc cac aac atc gag gac ggc agc gtg cag ctc gcc gac cac tac cag cag aac acc ccc atc ggc gac ggc ccc gtg ctg ctg ccc gac aac cac tac ctg agc acc cag tcc aag ctg agc aaa gac ccc aac gag aag cgc gat cac atg gtc ctg ctg gag ttc gtg acc gcc gcc ggg atc act ctc ggc atg gac gag ctg tac aag gca gga ggc agc CAT GAT GGA CAT GAT GAT GAT GTG ATT GAT ATT GAG GAT GAC CTT GAC GAT GTC ATT GAA GAG GTA GAA GAC TCA AAA CCA GAT ACC ACT GCT CCT CCT TCA TCT CCC AAG GTT ACT TAC AAA GCT CCA GTT CCA ACA GGG GAA GTA TAT TTT GCT GAT TCT TTT GAC AGA GGA ACT CTG TCA GGG TGG ATT TTA TCC AAA GCC AAG AAA GAC GAT ACC GAT GAT GAA ATT GCC AAA TAT GAT GGA AAG TGG GAG GTA GAG GAA ATG AAG GAG TCA AAG CTT CCA GGT GAT AAA GGA CTT GTG TTG ATG TCT CGG GCC AAG CAT CAT GCC ATC TCT GCT AAA CTG AAC AAG CCC TTC CTG TTT GAC ACC AAG CCT CTC ATT GTT CAG TAT GAG GTT AAT TTC CAA AAT GGA ATA GAA TGT GGT GGT GCC TAT GTG AAA CTG CTT TCT AAA ACA CCA GAA CTC AAC CTG GAT CAG TTC CAT GAC AAG ACC CCT TAT ACG ATT ATG TTT GGT CCA GAT AAA TGT GGA GAG GAC TAT AAA CTG CAC TTC ATC TTC CGA CAC AAA AAC CCC AAA ACG GGT ATC TAT GAA GAA AAA CAT GCT AAG AGG CCA GAT GCA GAT CTG AAG ACC TAT TTT ACT GAT AAG AAA ACA CAT CTT TAC ACA CTA ATC TTG AAT CCA GAT AAT AGT TTT GAA ATA CTG GTT GAC CAA TCT GTG GTG AAT AGT GGA AAT CTG CTC AAT GAC ATG ACT CCT CCT GTA AAT CCT TCA CGT GAA ATT GAG GAC CCA GAA GAC CGG AAG CCC GAG GAT TGG GAT GAA AGA CCA AAA ATC CCA GAT CCA GAA GCT GTC AAG CCA GAT GAC TGG GAT GAA GAT GCC CCT GCT AAG ATT CCA GAT GAA GAG GCC ACA AAA CCC GAA GGC TGG TTA GAT GAT GAG CCT GAG TAC GTA CCT GAT CCA GAC GCA GAG AAA CCT GAG GAT TGG GAT GAA GAC ATG GAT GGA GAA TGG GAG GCT CCT CAG ATT GCC AAC CCT AGA TGT GAG TCA GCT CCT GGA TGT GGT GTC TGG CAG CGA CCT GTG ATT GAC AAC CCC AAT TAT AAA GGC AAA TGG AAG CCT CCT ATG ATT GAC AAT CCC AGT TAC CAG GGA ATC TGG AAA CCC AGG AAA ATA CCA AAT CCA GAT TTC TTT GAA GAT CTG GAA CCT TTC AGA ATG ACT CCT TTT AGT GCT ATT GGT TTG GAG CTG TGG TCC ATG ACC TCT GAC ATT TTT TTT GAC AAC TTT ATC ATT TGT GCT GAT CGA AGA ATA GTT GAT GAT TGG GCC AAT GAT GGA TGG GGC CTG AAG AAA GCT GCT GAT GGG GCT GCT GAG CCA GGC GTT GTG GGG CAG ATG ATC GAG GCA GCT GAA GAG CGC CCG TGG CTG TGG GTA GTC TAT ATT CTA ACT GTA GCC CTT CCT GTG TTC CTG GTT ATC CTC TTC TGC TGT TCT GGA AAG AAA CAG ACC AGT GGT ATG GAG TAT AAG AAA ACT GAT GCA CCT CAA CCG GAT GTG AAG GAA GAG GAA GAA GAG AAG GAA GAG GAA AAG GAC AAG GGA GAT GAG GAG GAG GAA GGA GAA GAG AAA CTT GAA GAG AAA CAG AAA AGT GAT GCT GAA GAA GAT GGT GGC ACT GTC AGT CAA GAG GAG GAA GAC AGA AAA CCT AAA GCA GAG GAG GAT GAA ATT TTG AAC AGA TCA CCA AGA AAC AGA AAG CCA CGA AGA GAG TAG.

### Cell culture, transfection, siRNA knock down, chemical inhibitors

Cell culture conditions included incubation at 37°C, 5% CO2 and 90% humidity in Dulbecco’s Modified Eagle Medium (DMEM, Corning, 17-205-CV) containing 10% fetal bovine serum (FBS, Corning 35-011-CV), 1x L-glutamine (Corning, 25-005-CL). Cells used in experiments were never allowed to grow beyond 80% confluency and regularly split every 2 or 3 d using 0.05% Trypsin/0.53 mM EDTA in HBSS (Corning, 25-051-CL). Cells were typically cultured on tissue culture treated plastic (SCBT, sc-251460) and seeded onto #1.5 coverslip glass-bottom dishes (Cell Vis, D35C4-20-1.5-N) for imaging.

Transfections of YFP-PrP* NRK cells were performed at 70% confluency using PolyJet^TM^ In Vitro DNA Transfection Reagent (SignaGen Laboratories, SL100688). For all experiments involving transient transfections, we allowed time for cells that were overexpressing the plasmid-encoded protein to die. We split transiently transfected cells 24 h after transfection onto #1.5 glass-bottom coverslips (Cell Vis, D35C4-20-1.5-N) for imaging experiments. Imaging experiments were performed 48–72 h after transfection.

siRNA knockdown of TMED9 was carried out by two sequential transfections 48 h apart of the ON-TARGETplus SMARTpool Rat TMED9 siRNA (Dharmacon, L-092937-01-0005) with Dharmafect 2 transfection reagent (Dharmacon, T-2002-02) exactly as recommended by the manufacturer. For imaging experiments, immediately after the second siRNA transfection, cells were transferred to #1.5 coverslips (Cell Vis, D35C4-20-1.5-N), and 24 h later, the imaging experiments were performed. For western blots, immediately after the second siRNA transfection, cells were transferred to 6-well or 12-well tissue culture-treated dishes (Genesee, 25-105MP or 25-106MP).

Chemical inhibitors were added to cell culture in complete DMEM at the following working concentrations: 0.1 µM thapsigargin (diluted from 10mM stock prepared in DMSO; Calbiochem, 586006), 100 µM BRD4780 (also called AGN 192403 hydrochloride; diluted from 50mM stock prepared in water; Tocris, 1072), 10 µM of MG132 (diluted from 20mM stock prepared in DMSO; Calbiochem, 474790), 250 nM bafilomycin A1 (diluted from 1mM stock prepared in DMSO; SCBT, SC-201550B), 10 mM 3-methyladenine (3MA, diluted directly into cell culture medium at 10mM; Tocris, 3977), 2.5 µM brefeldin A (diluted from 25mM stock prepared in DMSO; LC Laboratories, B-8500) and 50 µg/ml cycloheximide (diluted from 10mg/ml stock prepared in water; Calbiochem, 23-97-64). As a control for either thapsigargin or bafilomycin-treatments, the equivalent concentrations of DMSO was added to cells. For BRD4780, 3MA, and CHX, since they were originally diluted in water, control conditions are simply the cell culture medium without the drug. The single exception to this was in the drug panel presented in Figure 5A, where as explicitly stated in the Figure Legend, DMSO controls were not included.

### Co-immunoprecipitation

Cells were cultured in 10 cm dishes to ∼70% confluency and treated as described. The entire procedure prior to elution was performed in a 4°C cold room and on ice to stabilize interactions. First, cells were washed twice with 5 mL of cold Low Salt Buffer (50 mM HEPES, pH7.4, 100 mM NaCl, 2 mM MgCl_2_). Cells were lysed with 1 mL of IP buffer (1% CHAPS, 50 mM HEPES, pH 7.4, 100 mM NaCl, and 2 mM CaCl2) and transferred to a microfuge tube. The lysate was pipetted up and down 20 times through a 200 µl pipet tip to disrupt the cell membranes. To clear the lysates of debris that may clog the column, lysates were subjected to two rounds of centrifugation for 5 min at 16,000 g at 4°C, and supernatants were transferred to new 1.5 mL microfuge tube after each centrifugation. 90 μL of the cleared lysate were set aside to load as “inputs” and the remaining 900 μL were incubated on ice for 15 min with 60 μL of super-paramagnetic eGFP µMACS MicroBead from the μMACS GFP Isolation Kit (Miltenyi, 130-091-288). In parallel, µColumns (Miltenyi, 130-042-701) were placed in the µMacs Separator (Miltenyi, 130-042-602) and equilibrated with 200 µL of IP buffer. At the end of the incubation period, the bead/lysate mixtures were loaded into the columns. Columns were then rinsed five times with 200 μL of IP buffer and washed once with 20 mM Tris Buffer, pH 7.5. After the washes, the experiment was transferred to room temperature and elution was performed by adding 40 μL of boiling elution buffer made of 2x Laemmli Sample Buffer (Bio-Rad, 1610737) and 20mM TCEP (tris(2-carboxyethyl)phosphine), Sigma, 646547). The input samples were denatured with 35 μL of 4X sample buffer (Bio-Rad, 1610747) with 40mM TCEP. The input and eluate samples were further analyzed by western blot.

### Western blots and antibodies used for western blots

After treatments, cells were washed with cold 1X phosphate-buffered saline (PBS), lysed in a small volume of cold radioimmunoprecipitation assay buffer (RIPA) buffer (50 mM Tris HCl, pH 7.6, 150 mM NaCl, 1% NP-40, 0.5% sodium deoxycholate and 0.1% SDS). Protein lysates were centrifuged at 4°C for 10 min at 15,000 rpm on a microcentrifuge. Supernatants were stored at −20°C and processed as follows. Protein lysate concentrations were determined with Pierce Micro BCA Protein Assay kit (ThermoFisher, 23235). Equal total quantities of protein were denatured using Laemmli sample buffer supplemented with tris(2-carboxyethyl)phosphine (TCEP) and boiled for 5 min before loading into a 12% or 4-20% acrylamide pre-cast SDS-PAGE gels (Bio-Rad, 4561045) for electrophoresis. Where indicated, we ran samples on Bio-Rad 4–20% Mini-PROTEAN® TGX Stain-Free™ Protein Gels (Bio-Rad, 4568094) to obtain images of total protein inputted into the gel using Bio-Rad’s Stain-Free™ Imaging system on the ChemiDoc™ Touch Imaging System (Bio-Rad, 17001401).

Proteins were transferred to a PVDF (polyvinylidene difluoride, Bio-Rad, 1704275) membrane using a Trans-Blot Turbo Transfer System (Bio-Rad, 1704150). Membranes were blocked with 5% non-fat milk in Tris-buffered saline with 0.1% Tween-20 (TBS-T + 5% Milk) for 1 hour at room temperature prior to probing with primary antibodies.

The following primary antibodies were added to the TBS-T + 2.5% milk solution at the following concentrations: mouse monoclonal αGFP (Proteintech, 66002-1-IG) at 1:2000, rabbit polyclonal αTMED9 (Proteintech, 21620-1-AP) at 1:2500, rabbit polyclonal αTMP21 (homemade and described previously [12]) at 1:2000, rabbit polyclonal αTMED2 (Proteintech, 11981-1-AP) at 1:2000, mouse monoclonal αCalnexin (Proteintech, 66903-1) at 1:2500, rabbit polyclonal αPrP (Proteintech, 12555-1-AP) at 1:2500, mouse monoclonal αGAPDH antibody (ThermoFisher, MA5-15738) at 1:4000, for 2 hours at room temperature or overnight at 4°C.

Membranes were washed three times with TBS-T and incubated in TBS-T + 5% milk for 1 hour at room temperature with either of the following secondary antibodies at 1:5000: HRP-conjugated Donkey anti-Rabbit (Sigma Aldrich, NA934-1ML) or HRP-conjugated Goat anti-Mouse (ThermoFisher, A28177). Membranes were washed three times, and briefly incubated in in Pierce™ ECL Western Blotting Substrate (ThermoFisher, 32209) or SuperSignal™ West Femto Maximum Sensitivity Substrate (ThermoFisher, 34096) prior to imaging the chemiluminescence on the ChemiDoc™ Touch Imaging System (Bio-Rad, 17001401).

FIJI (ImageJ 1.52p, NIH USA) was used to measure relative pixel densities of bands on unsaturated images collected as eight-bit .tif files from the Chemi-Doc (Bio-Rad). A rectangular ROI sized to enclose the band with the largest area was created and the same ROI was used to measure the integrated density of the rest of the bands. Integrated density values were background subtracted and normalized against “time 0,” “untreated” or “control” bands to obtain relative pixel densities. Relative pixel density values from three independently performed experiments were presented as mean ± SD (n = 3), as indicated in the figure legends. Means and SD were calculated and plotted using Microsoft Excel. Quantification details for specific experiments are provided in the adjoining figure legends.

### Glycosidase digests

Cell lysates were divided equally into multiple samples that were treated individually with water (no digested control), or with Endoglycosidase H (Endo H, New England Biolabs, P0702), with a2-3,6,8,9 Neuraminidase A (neuraminidase, New England Biolabs, P0722), with a combination of neuraminidase and O-Glycosidase (NA and O-G, New England Biolabs, P0733), with Peptide-N-glycosidase F (PNGase F, New England Biolabs, P0704) or with a combination of O-Glyc+PNGase (O-G + PNGase F) for 3 hrs at 37°C according to the manufacturer’s instructions.

To draw the band density histograms associated with the glycosidase digests, FIJI (ImageJ 1.52p, NIH USA) was used. Since the Endo H digest in the untreated lysate produced bands spanned the greatest distance from highest molecular weight band to lowest molecular weight band, a rectangular ROI sized to enclose the Endo H bands was created. This same ROI was dragged from lane to lane and the plot graph was generated by selecting the “Analyze,” “Gel” and “Plot lanes” menu items in Image J and graphs representing the densitometry intensity versus pixel distance were generated by selecting the “Analyze,” “Tools” menu items in Image J to run the Analyze Line Graph.

### Live-cell imaging, immunofluorescence and antibodies, nuclear stain and the microscope

Cells were seeded on a #1.5 glass bottom dish (Cell Vis, D35-20-1.5-N or D35C4-20-1.5-N) 24-48 hrs prior to the imaging experiment. Cells were imaged at 60-80% confluency. Treatment conditions (e.g. untreated, BRD4780 or TG-treatment) were specified in the figure legend. For live-cell experiments and time-lapse imaging, cells were imaged in their normal DMEM medium inside of a pre-equilibrated stage top incubator chamber adjusted to 37°C, 5% CO2 and 90% humidity. To stain nuclei for live cell imaging, cells were incubated for 10 min with NucBlue™ Live ReadyProbes™ Reagent Hoechst 33342 (Thermofisher, R37605).

For immunofluorescence against calnexin (CNX), TMED9, and GM130, cells were washed once with PBS and then fixed and permeabilized with methanol/acetone (3:1) solution for 10 min at −20°C. For immunofluorescence against cell surface GFP without additional staining, cells were washed once with PBS and then fixed with 4% paraformaldehyde (PFA) in PBS for 15 min at room temperature, but not permeabilized. After the fixation step, cells were washed three times with PBS and blocked for 30 min at room temperature with UltraCruz blocking reagent (SantaCruz; SC-516214) to stain GFP, calnexin, or GM130, or 5% normal goat serum (ThermoFisher, 31872) diluted into PBS to stain TMED9. Primary antibodies were diluted 1:200 in either UltraCruz blocking reagent or 2.5% normal goat serum in PBS buffer and added to the cells for 2 h at room temperature. The following primary antibodies were used: mouse monoclonal αGFP (Proteintech, 66002-1-IG), rabbit polyclonal αTMED9 (Novus, NBP1-31650, lot# 40121), mouse monoclonal αCalnexin (Proteintech, 66903-1), or rabbit polyclonal αGM130 (Novus Bio, NBP2-53420). Cells were washed 3 times with PBS. Secondary antibodies, Goat anti-Rabbit IgG, Alexa Fluor™ 660 (ThermoFisher Scientific, A21074) or Goat anti-Mouse IgG, Alexa Fluor™ 568 (ThermoFisher Scientific, A1103), were diluted at 1:2500 in either UltraCruz blocking reagent or 2.5% normal goat serum in PBS buffer and added to the cells for 1 h at room temperature. Cells were washed 3 times with PBS and incubated with fresh PBS with NucBlue nuclear stain (ThermoFisher, 33342) diluted at 1:20. Images were captured using the Nikon spinning disk (described below).

For immunofluorescence against cell surface GFP followed by immunofluorescence of TMED9, live unpermeabilized cells were first incubated with Alexa Fluor^TM^ 594 (AF594)-conjugated Rabbit αGFP antibody (Thermofisher, A-21312) for 10 min. The AF594-αGFP antibody was diluted at 1:200 directly into the cell culture medium. Next, the cells were fixed and permeabilized with methanol/acetone (3:1) solution for 10 min at −20°C. Cells were blocked with 5% normal goat serum in PBS (Thermofisher, 31872) and stained with mouse monoclonal αTMED9 (Novus, NBP3-33028) primary antibody at 1:500 followed by Goat anti-Mouse IgG, Alexa Fluor™ 660 (ThermoFisher Scientific, A-21055) secondary antibody diluted at 1:2000.

Images were acquired using a Nikon inverted spinning disk confocal microscope equipped with a Yokogawa CSU-X1 Spinning Disk, EMCCD camera, 60x Plan Apo 1.40 NA oil / 0.13mm WD, Tokai live-cell incubator, Perfect Focus system to prevent focus drift, lasers (405 nm, 445 nm, 488nm, 514nm, 561nm, 647 nm), emission filter sets (455/50, 480/40, 525/36, 540/30, 605/70, and 700/75), and dichroic mirror sets (405/488/561/640 and 440/514/561). This system is set up to prevent bleed-through between the following sets of fluorophores or fluorescent proteins: Cerulean, YFP, FusionRed/mCherry/AF568/AF594 and AF647/AF660, or Hoechst/DAPI, YFP, FusionRed/mCherry/AF568/AF594 and AF647/AF660, or Hoechst/DAPI, GFP and FusionRed/mCherry/AF568/AF594 and AF647/AF660. The Nikon microscope and image acquisition was controlled by the NIS-Elements software.

### Antibody Uptake

Cells were seeded in 4-well #1.5 glass bottom dish (Cell Vis D35C4-20-1.5-N) 24 hrs prior to the live-cell imaging or immunofluorescence experiment. The following were added to the cell culture medium: a 1:5000 dilution of Alexa Fluor^TM^ 647-conjugated Rabbit anti-GFP antibodies (Thermofisher, A-31852), 0.1 µM thapsigargin (Calbiochem, 586006) to induce RESET and 250nM balifomycin A1 (SCBT, SC-201550B) to inhibit lysosomal degradation. Cells were imaged live at 37°C, 5% CO2 and 90% humidity for 90 min or incubated in incubator for 90 min prior to fixation. Cells were fixed in methanol/acetone (3:1) solution for 10 min at −20°C and stained for TMED9 using a mouse monoclonal anti-TMED9 (Novus NBP3-33028).

### Image analysis

Pearson colocalization coefficients *r* were obtained by the following steps. In cells co-expressing or stained for two non-cross-talking fluorescent proteins or dyes, multichannel images were captured and cropped tightly around a single cell. The cell edge was identified by temporarily increasing the contrast of the image and ROIs were manually drawn around the edges of the cell and saved using ROI manager. Within the ROI for each set of 2 channels, Pearson’s correlation coefficient value *r* was measured using the Coloc2 plugin by selecting Analyze, Colocalization, and Coloc 2 commands in FIJI (ImageJ 1.52p, NIH USA).

To measure the fluorescence intensity of ER-localized CER-TMED9 within an imaging generate ER mask, cells were co-transfected with mCh-Sec61β to mark the ER. Multichannel images were collected before and after treatment in RFP (for mCh-Sec61β) and CFP (for CER-TMED9) channels and cropped tightly around a single cell. For each cell, the cell edge was identified by temporarily increasing the contrast of the image and ROIs were manually drawn around the edges of the cell and saved using ROI manager. ER masks were created from Sec61-mCherry images using Fiji (NIH Image J, 1.52p) by setting the threshold to cover a maximum of the ER compartment by selecting the following Image, Adjust from the menu, and selecting the Threshold command. The threshold setting was used to create and save the ER-mask by selecting Edit, Selection from the menu and selecting the Create Mask command. ER-localized CER-TMED9 fluorescence intensity was extrapolated using the ER mask by selecting Process from the menu and the Image Calculator command with Image 1 set as “ER mark”, operation set as “AND” and Image 2 set as “CER-TMED9”. The ER-localized CER-TMED9 fluorescence intensity was then measured using Measurement command.

### Statistics

Statistical analyses were performed using Prism 9 (GraphPad). Statistical significance was calculated from unpaired t-test with Welch’s correction on the mean value of at least 3 independent experiments. The p values that were equal or less than 0.05, were annotated with one star (*); equal or less than 0.01 were annotated with two stars (**); equal or less than 0.001, were annotated with three stars (***); equal or less than 0.0001, were annotated with three stars (****).

## Acknowledgements

We thank Alexandra Graninger for technical advice on the glycosidase digests and scientific discussions. We thank Xin Xiang and Yihong Ye for critical feedback on the study and manuscript. We are grateful to Rafael Villasmil of the NEI Flow Cytometry Core for cell sorting. We thank Meera Krishnan for technical help. This research was supported by National Institute of General Medical Sciences, National Institutes of Health grant no. R01 GM134327 and Uniformed Services University of the Health Sciences startup funds to P. Satpute-Krishnan.

## Abbreviations List

ER: Endoplasmic Reticulum
ERGIC: ER-Golgi Intermediate Compartment
PQC: Protein Quality Control
NRK: Normal Rat Kidney
GPI: Glycosylphosphatidylinositol
GPI-AP: Glycosylphosphatidylinositol-Anchored Protein
PrP: Prion Protein
YFP-PrP*: YFP-tagged Prion Protein C179A
YFP-PrP* NRK: stable transfectants of NRK expressing YFP-PrP*
YFP-PrP WT NRK: stable transfectants of NRK expressing YFP-PrP WT
GFP-CD59 C94S NRK: stable transfectants of NRK expressing GFP-CD59 C94S
ATZ: Alpha-1-antitrypsin Z variant
MUC1-fs: Mucin-1 frame shift mutant
CNX: Calnexin
RESET: Rapid ER Stress-induced Export
ERAD: ER-Associated Degradation
ERLAD: ER-to-Lysosome-Associated Degradation
siTMED9: TMED9 siRNA
CTL: Control
AF: Alexa Fluor^TM^ (in context of AF568, AF594, AF647 and AF660)
Ab: Antibody
LMW band of TMED9: Lower Molecular Weight band of TMED9
HMW band of TMED9: Higher Molecular Weight band of TMED9
YFP: Yellow Fluorescent Protein
GFP: Green Fluorescent Protein
CER: Cerulean
TG: Thapsigargin
BafA1: Bafilomycin A1
3MA: 3-Methyladenine
BFA: Brefeldin A
CHX: Cycloheximide
PNGase F or PNG: Peptide-N-Glycosidase F
Endo H: Endoglycosidase H
NA: Neuraminidase
O-G: O-Glycosidase

## Figure Legends and Figures listed in the order they appear in the text

**Supporting Figure S1:**
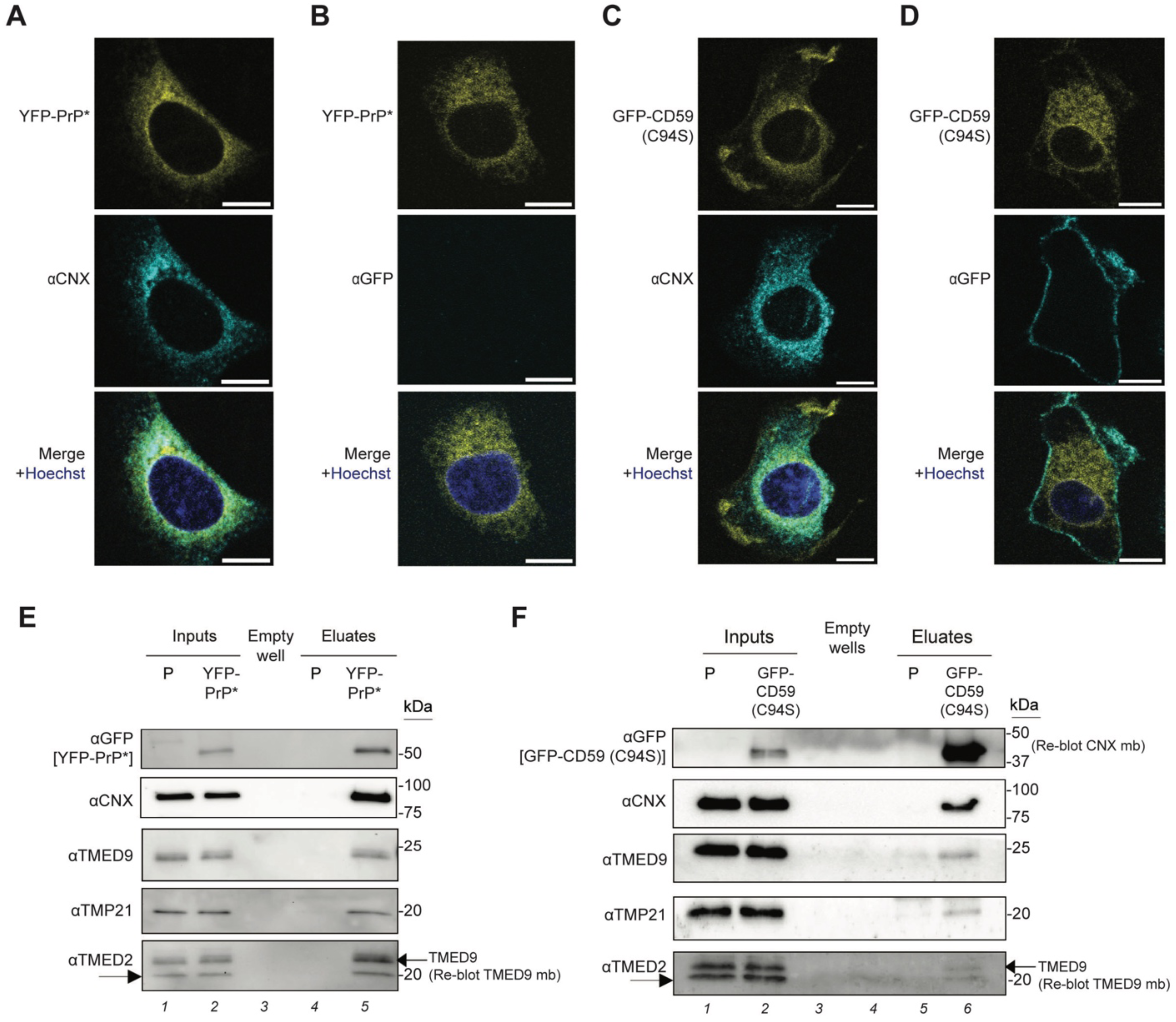
Misfolded GPI-APs localize to the ER and interact with calnexin, TMP21, TMED2, and TMED9. **(A-B)** Confocal images of YFP-PrP* NRK cells at steady state. Scale bar represents 10 µm. **(A)** Immunofluorescence image of endogenous calnexin (CNX) in a YFP-PrP* NRK cell. The nucleus was stained with Hoechst. **(B)** Immunofluorescence image of cell-surface YFP-PrP* using anti-GFP antibody on cells that were not permeabilized. The nucleus was stained with Hoechst. **(C-D)** Confocal images of GFP-CD59 (C94S) NRK cells at steady state. Scale bar represents 10 µm. **(C)** Immunofluorescence image against endogenous calnexin (CNX) in GFP-CD59 (C94S). The nucleus was stained with Hoechst. **(D)** Immunofluorescence of cell-surface GFP-CD59 (C94S) using anti-GFP antibody on cells that were not permeabilized. The nucleus was stained with Hoechst. **(E)** Western blots of GFP-tag co-immunoprecipitates from the parental untransfected NRK cells (P) or stably transfected YFP-PrP* NRK cells at steady state (n=1). Blots were probed for GFP for YFP-PrP*, and probed for endogenous calnexin (CNX), TMP21, TMED2 and TMED9. **(F)** Western blots of GFP-tag co-immunoprecipitates from parental untransfected NRK cells (P) or stably transfected GFP-CD59 (C94S) NRK cells (n=1). Blots were probed for GFP for GFP-CD59 (C94S), and probed for endogenous calnexin (CNX), TMP21, TMED2 and TMED9.

**Supplementary Figure S2:**
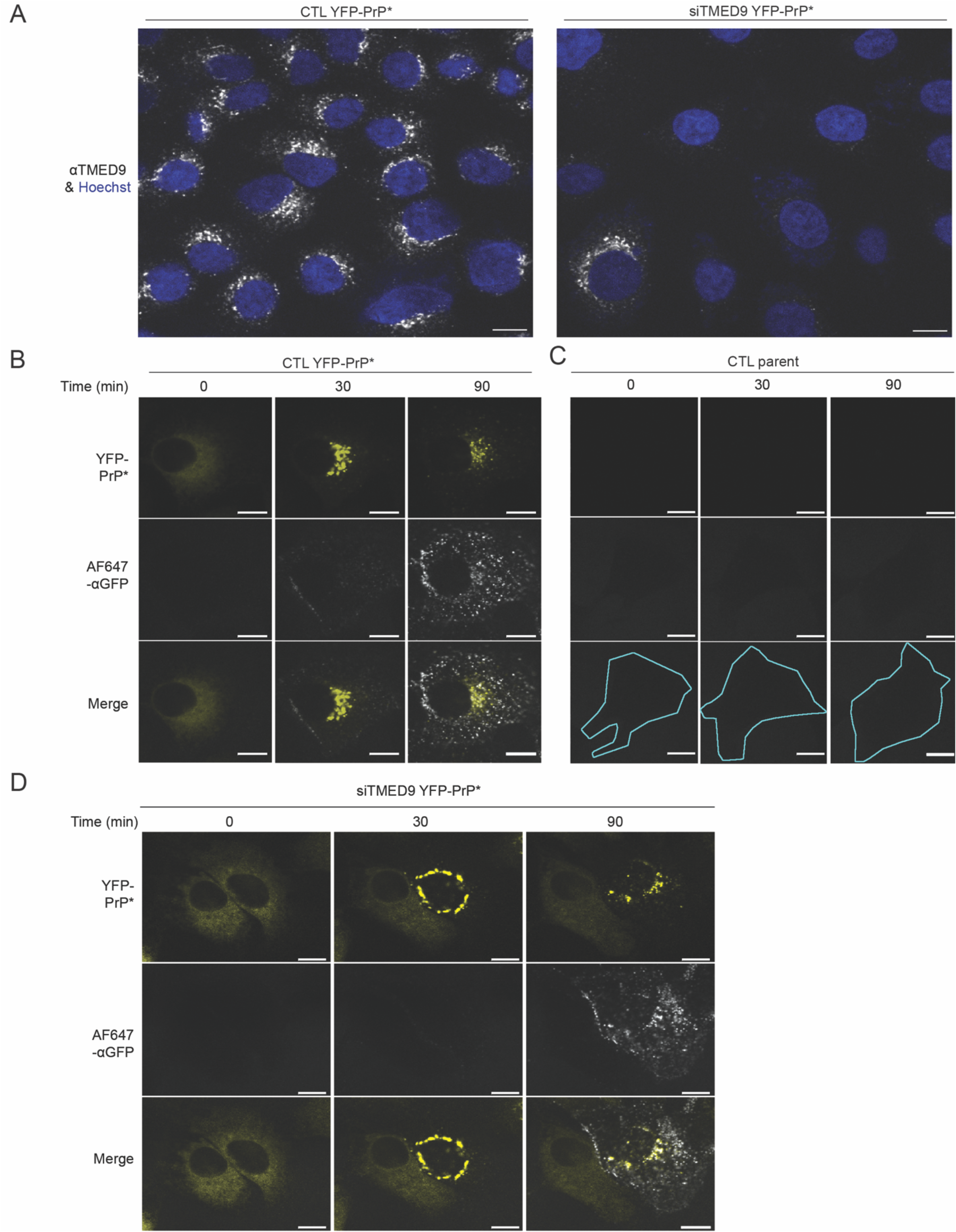
TMED9 knockdown blocks RESET and consequent anti-GFP-antibody uptake in YFP-PrP* NRK cells. **(A)** Immunofluorescence images of TMED9 staining in YFP-PrP* NRK cells that were treated with scrambled control siRNA (CTL) or siRNA against TMED9 (siTMED9). With our TMED9-knockdown protocol, the majority of siTMED9-treated YFP-PrP* NRK cells demonstrated TMED9 knock-down. A minor population (∼30-35%) of the siTMED9-treated cells did not demonstrate knock-down of TMED9. **(B-D)** Time-lapse images of cells collected immediately after the addition of thapsigargin, Alexa Fluor 647 (AF647)-conjugated rabbit anti-GFP antibodies, and bafilomycin A1. There is a faint ambient signal from the AF647-conjugated antibody in the medium that can be enhanced by increasing the gain. Scale bar represents 10 µm. **(B)** YFP-PrP* NRK cell that was treated with scrambled control siRNA (CTL). **(C)** Parental untransfected NRK cell that was treated with scrambled control siRNA (CTL). The outline of the cell has been hand-traced in cyan. **(D)** YFP-PrP* NRK cells that were treated with siRNA against TMED9 (siTMED9). A field of view that contained one cell that displayed the phenotype of complete TMED9 knockdown versus one cell with incomplete knockdown was selected for comparison. This panel is associated with Video 2.

**Supporting Figure S3:**
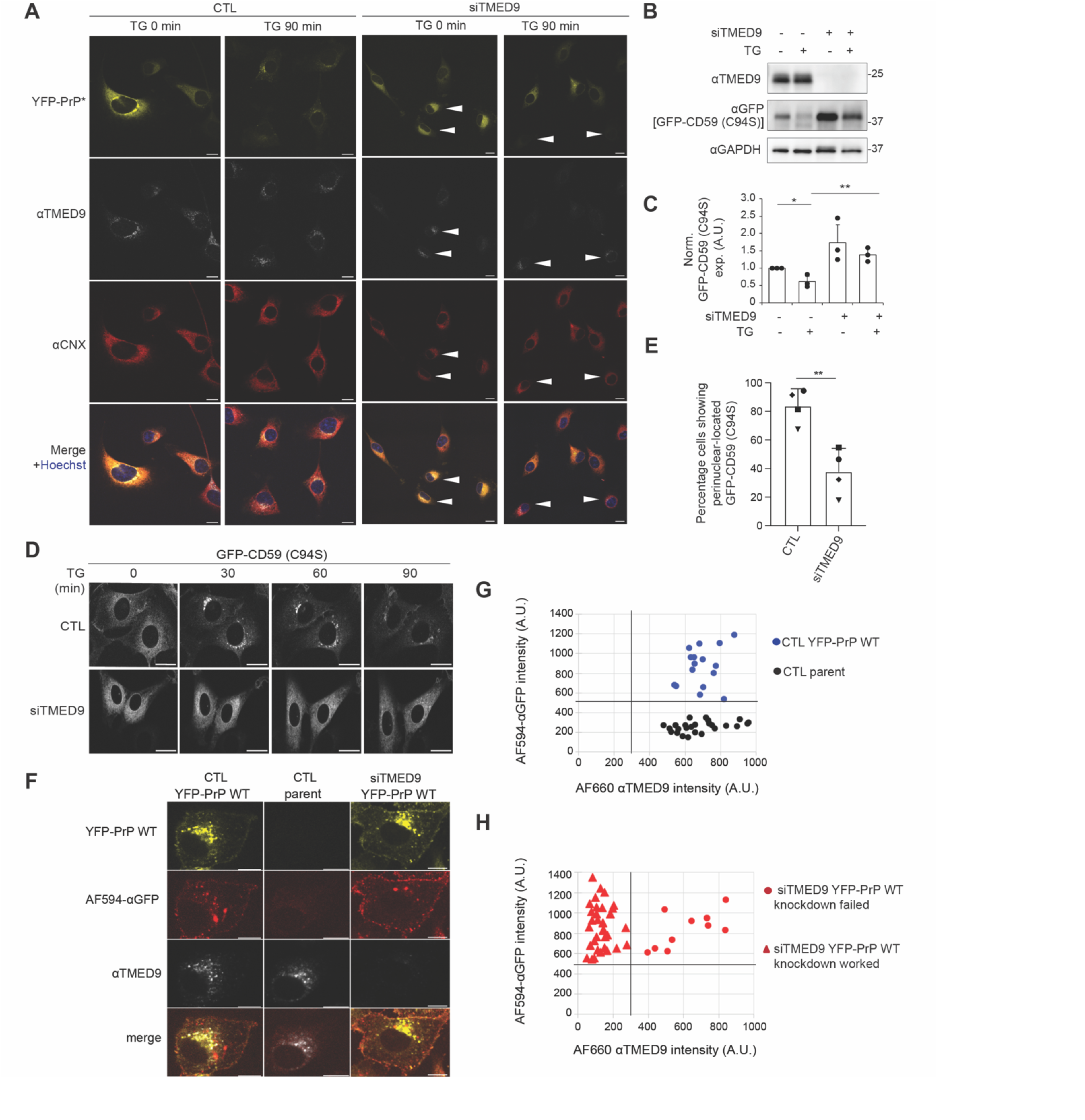
TMED9 is required for RESET of misfolded GPI-APs but is not required for ER-export of wild type GPI-APs. **(A)** Immunofluorescence image stained for endogenous TMED9 and CNX in control YFP-PrP* NRK (CTL) or siTMED9-treated YFP-PrP* NRK cells (siTMED9 cells) at steady state (TG 0min) after 90 min of TG-treatment (TG 90 min). Nuclei were stained with Hoechst. For the siTMED9-treated cells, large white arrows point to cells with only partial TMED9 knockdown. Scale bar represents 10 µm. **(B)** Representative western blots of either control (CTL) or TMED9 siRNA (siTMED9)-treated GFP-CD59 (C94S) NRK cells that were either untreated (TG –) or treated with 90 min TG (TG +) (n=3 biological replicates). **(C)** Bar graph representing the mean band intensity for GFP-CD59 (C94S) from 3 biological replicates as shown in (B). For each condition, GFP-CD59 (C94S) band intensities were double normalized. First, GFP band intensity was normalized against the band intensity of GAPDH. Second, the GFP band intensity was normalized against the untreated control (“siTMED9 – TG – “) band intensity. Error bars represent standard deviation of the mean. Symbols were coded for each independently performed experiment. Statistics were calculated from unpaired t-test with Welch’s correction with * indicated p<0.05 and ** indicated p<0.01. **(D)** Time-lapse images of control (CTL) or TMED9 siRNA (siTMED9) GFP-CD59 (C94S) NRK cells. Image-collection was started immediately after the addition of thapsigargin (TG). Scale bar represents 20 µm. **(E)** Percentage of cells showing a perinuclear Golgi-pattern for GFP-CD59 (C94S), indicative of Golgi-localization, after 30 min of TG-treatment. These data are derived from 4 independent experiments (n=4). Symbols were coded for each independently performed experiment. The number of cells analyzed for each biological replicate are as follows (CTL: triangle 37, diamond 59, circle 18, square 54; siTMED9: triangle 73, diamond 93, circle 28, square 64). Error bars represent standard deviation of the mean. Statistics were calculated from unpaired t-test with Welch’s correction with ** indicated p<0.01. The data underlying the graphs shown in Figure S3 are included in the S1_Data file. **(F)** Confocal images of control (CTL) scrambled siRNA treated YFP-PrP WT NRK cells or CTL parental NRK cells, or siTMED9-treated YFP-PrP WT NRK cells that were incubated with Alexa Fluor 594-conjugated anti-GFP antibodies for 10 minutes prior to permeabilization, then fixed, permeabilized and stained for TMED9. An Alexa Fluor^TM^ (AF) 660-conjugated secondary was used for the TMED9 immunofluorescence. All images were taken with identical imaging parameters. Scale bar represents 10 µm. **(G)** Plot of background-subtracted mean intensities measured in arbitrary units (A.U.) for cell-surface anti-GFP immunofluorescence and whole cell TMED9 immunofluorescence for scrambled siRNA-treated control (CTL) parental NRK (n=25) or CTL YFP-PrP WT NRK cells (n=16), as shown in (F). The cell surface YFP-PrP WT was stained with AF594-αGFP Ab. For the TMED9 immunofluorescence, an AF660-conjugated secondary was used. Images collected as shown and described in (F). For YFP-PrP WT cells, ROIs around the periphery of each cell was drawn by tracing the AF594-aGFP-staining. For CTL parent cells, ROIs were drawn around the periphery of the cells by temporarily increasing the gain in the 660 channel (TMED9-staining), because in addition to strong specific TMED9 binding, a small fraction of the primary antibody against TMED9 appeared to non-specifically bind the entire cytoplasm. Background was measured by creating an ROI within the same field of view where no cells were present. Once the cell peripheries were defined, the mean background-subtracted intensities for both AF660 (αTMED9) and AF594 (AF594-αGFP) were calculated. Both the scrambled siRNA control (CTL)-treated parental NRK cells and CTL YFP-PrP WT NRK cells served as positive controls for wild type TMED9 expression. A threshold (shown as bolded vertical grid line) for wild type TMED9 levels was arbitrarily placed to the left of the lowest TMED9-expressing scrambled siRNA-treated (CTL) cell. CTL parental NRK served as a negative control for AF594-αGFP Ab-binding. The CTL YFP-PrP WT NRK cells served as positive control for AF594-αGFP Ab-binding. A threshold (shown as bolded horizontal grid line) for AF594-αGFP Ab was arbitrarily placed below the lowest level of AF594-αGFP Ab mean fluorescence intensity of the positive control CTL YFP-PrP WT NRK, but above the highest level of AF594-αGFP Ab mean fluorescence intensity in the negative control CTL untransfected parental NRK. **(H)** Plot of background-subtracted mean intensities measured in arbitrary units (A.U.) for cell-surface anti-GFP immunofluorescence and whole cell TMED9 immunofluorescence for YFP-PrP WT NRK cells that were treated with siRNA against TMED9 (siTMED9) (n=45), as shown in (F). Cells were handled and images collected as described in (G), except that only one strategy was used to trace the periphery of the cells. Since every cell in this set stained for AF594-αGFP Ab, ROIs were drawn around the periphery of each cell by tracing the AF594 fluorescence. The mean intensities for both AF660 (αTMED9) and AF594 (AF594-αGFP) were measured within the periphery of the cell and background subtracted as described in (G). The data points were plotted on an identically proportioned graph to that in (G) so that direct comparisons of the data points could be made with the controls plotted in (G). Of the 45 randomly selected cells, 10 of the data points had mean TMED9 intensities within the CTL scrambled siRNA levels and were coded with a circle symbol to represent “siTMED9 YFP-PrP WT NRK knockdown failed.” These cells also displayed TMED9 fluorescence in the perinuclear Golgi pattern as observed for the CTL YFP-PrP WT shown in (F). 35 of the data points had mean TMED9 below the CTL levels and were coded with a triangle symbol to represent “siTMED9 YFP-PrP WT NRK knockdown worked.” As shown in (F), the siTMED9-treated cells continued to display YFP-PrP WT on the cell surface and bind to AF594-αGFP Ab even when TMED9 expression levels were below CTL levels.

**Supporting Figure S4:**
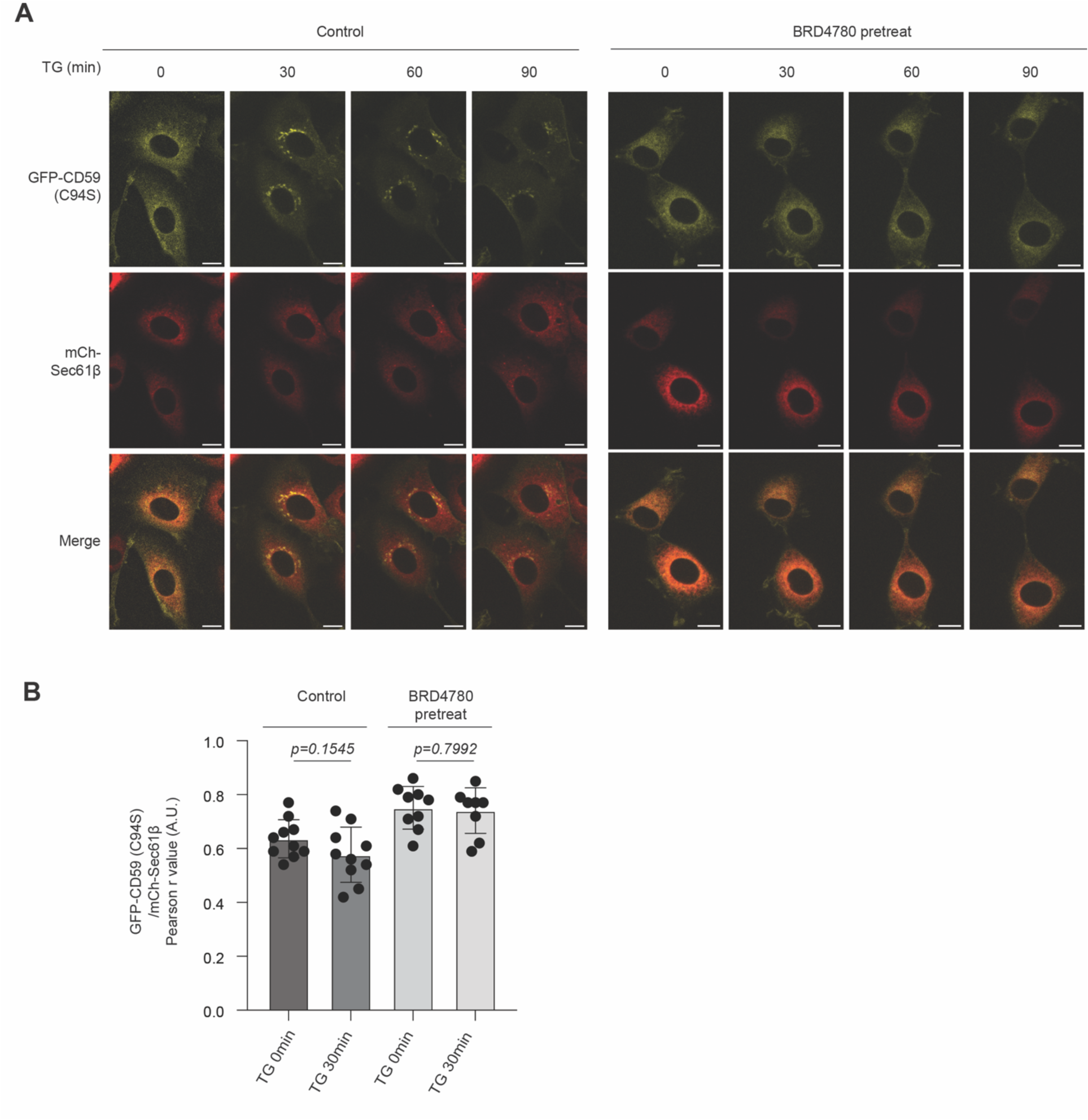
Chemical inhibitor of TMED9, BRD4780, inhibits RESET. **(A)** Representative time-lapse imaging of stably transfected GFP-CD59 C94S NRK cells in control condition (CTL) or pretreated with BRD4780 for 30 min (pretreated with BRD4780). Image-collection was started immediately after the addition of thapsigargin (TG). Scale bars represent 10 µm. **(B)** Plot of the average Pearson’s r values between GFP-CD59 (C94S) and mCh-Sec61β for Control vs. BRD4780-pretreated conditions, as described in (A). For Control and BRD4780-pretreated conditions, 10 and 9 time-lapses of individual cells, respectively, were analyzed for the 0 and 30 min time points. Pearson’s colocalization coefficients, r, were measured between GFP-CD59 (C94S) and mCh-Sec61β within the boundaries of the cell. For each data point, the boundaries of the cells were revealed by temporarily maximizing the gain for mCh-Sec61β. In CTL cells, the r values between GFP-CD59 (C94S) and mCh-Sec61β for 0 min and 30 min time points were 0.636 ± 0.067 and 0.577 ± 0.065, respectively. In BRD4780-treated cells, the r values between GFP-CD59 (C94S) and mCh-Sec61β for for 0 min and 30 min time points were 0.751 ± 0.075 and 0.741 ± 0.080, respectively.

**Supporting Figure S5:**
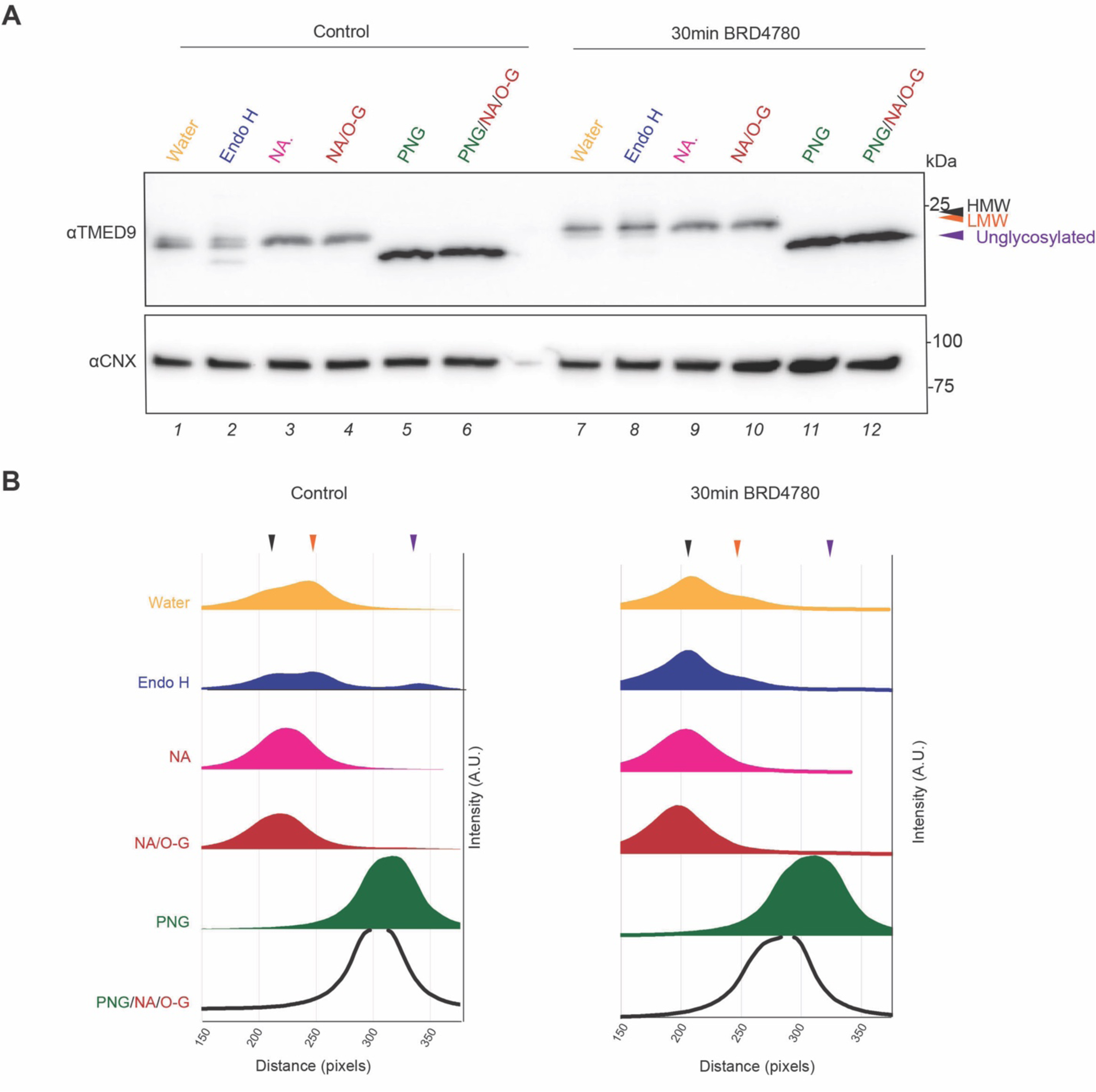
BRD4780-treatment for 30 min induces a shift in the TMED9 population towards endo H resistance and neuraminidase-sensitivity. (**A**) Western blots of digested lysates of YFP-PrP* NRK cells in control and BRD4780-treated conditions. Protein lysates were not digested (water) or digested with Endoglycosidase H (Endo H), Neuraminidase only (NA), Neuraminidase+O-Glycosidase (NA+O-G.), Peptide-N-glycosidase F (PNG) or Neuraminidase+O-Glycosidase + Peptide-N-glycosidase F (NA+O-G+PNG) for 3 h at 37°C (n=1 experiment). Blots were probed for TMED9 and calnexin (CNX). (**B**) Western blot band graphics were obtained by first analyzing the plot lanes and second by generating line graphs of pre-selected regions of interest (ROI) from the western blot for control samples (left) or BRD4780 (30 min)-treated samples (right). Black arrow represents the HMW form, orange arrow represents the LMW and purple, the unglycosylated form of TMED9.

**Supporting Figure S6:**
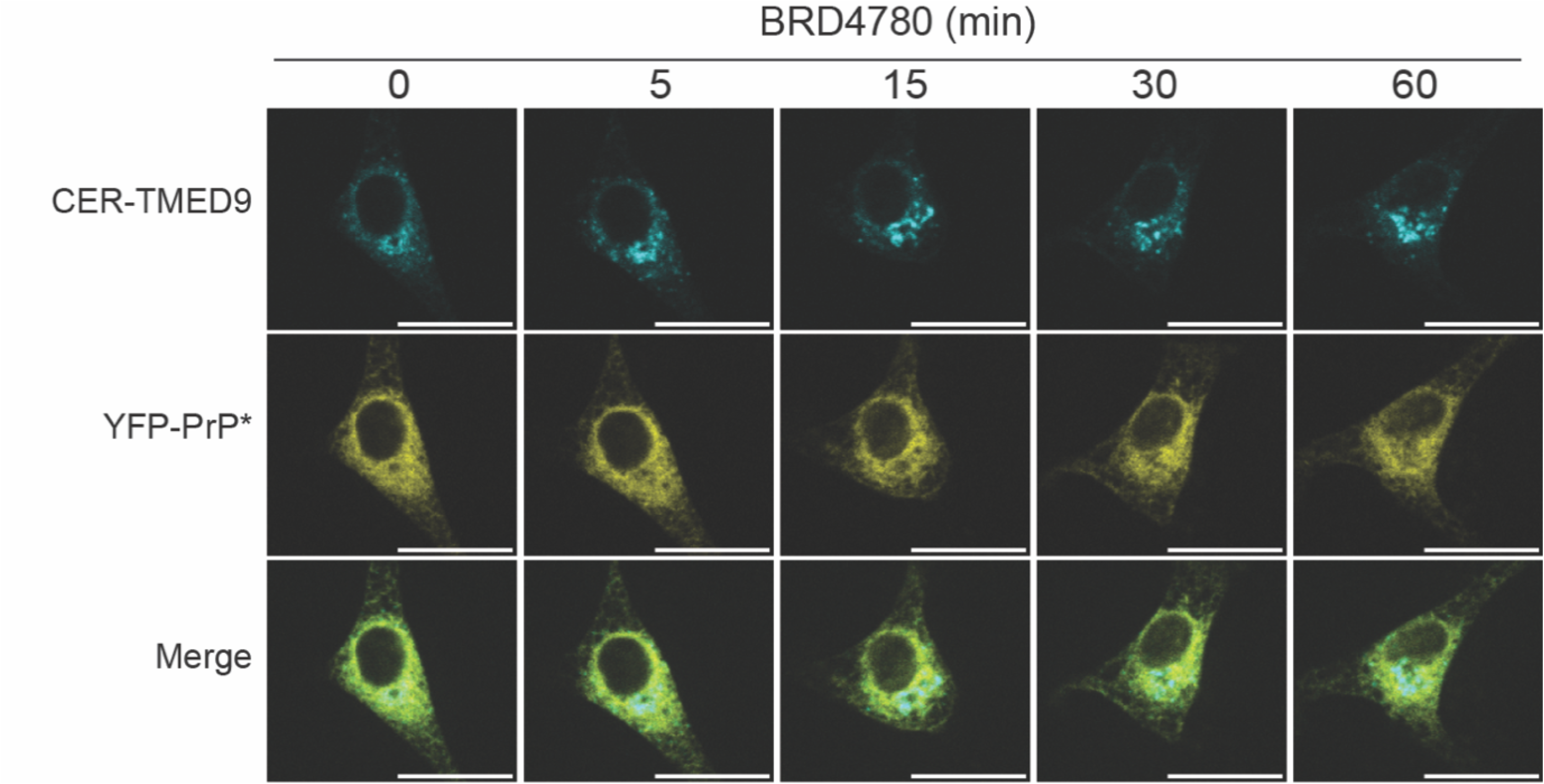
Cerulean-TMED9 relocalizes upon BRD4780-treatment leaving YFP-PrP* behind in the ER. Time-lapse images of a typical YFP-PrP* NRK cell that was transfected with CER-TMED9. Time-lapse image collection was started immediately after the addition of 100 µM BRD4780 treatment. Scale bar represents 20 µm.

**Supporting Figure S7:**
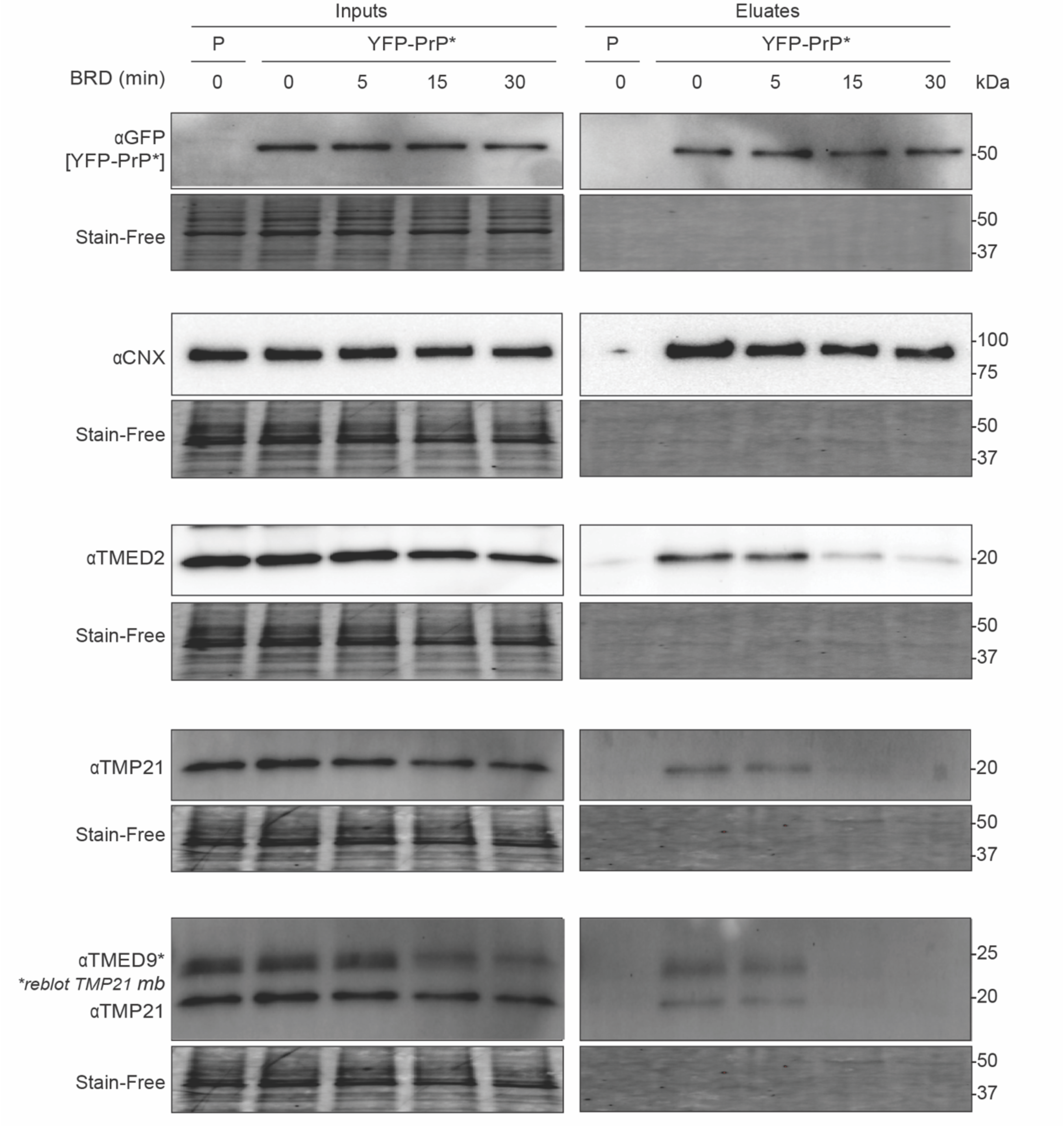
BRD4780 treatment dissociates TMED9, TMP21 and TMED2 from YFP-PrP*. Western blots of GFP-tag co-immunoprecipitations (co-IPs) from the parental untransfected NRK cells (P) or stably transfected YFP-PrP* NRK cells (n=1 experiment). YFP-PrP* NRK cells were treated with 100 µM BRD4780 (BRD) and collected for co-IP at the indicated time points. Cells were harvested at the indicated time points for co-immunoprecipitation of YFP-PrP* with anti-GFP antibody conjugated beads in addition to GFP to detect co-immunoprecipitation of YFP-PrP* constructs, blots were probed for endogenous calnexin (CNX), TMED2, TMP21, and TMED9. Under each western blot is depicted a “Stain-Free” image of the total protein in the gel.

**Supporting Figure S8:**
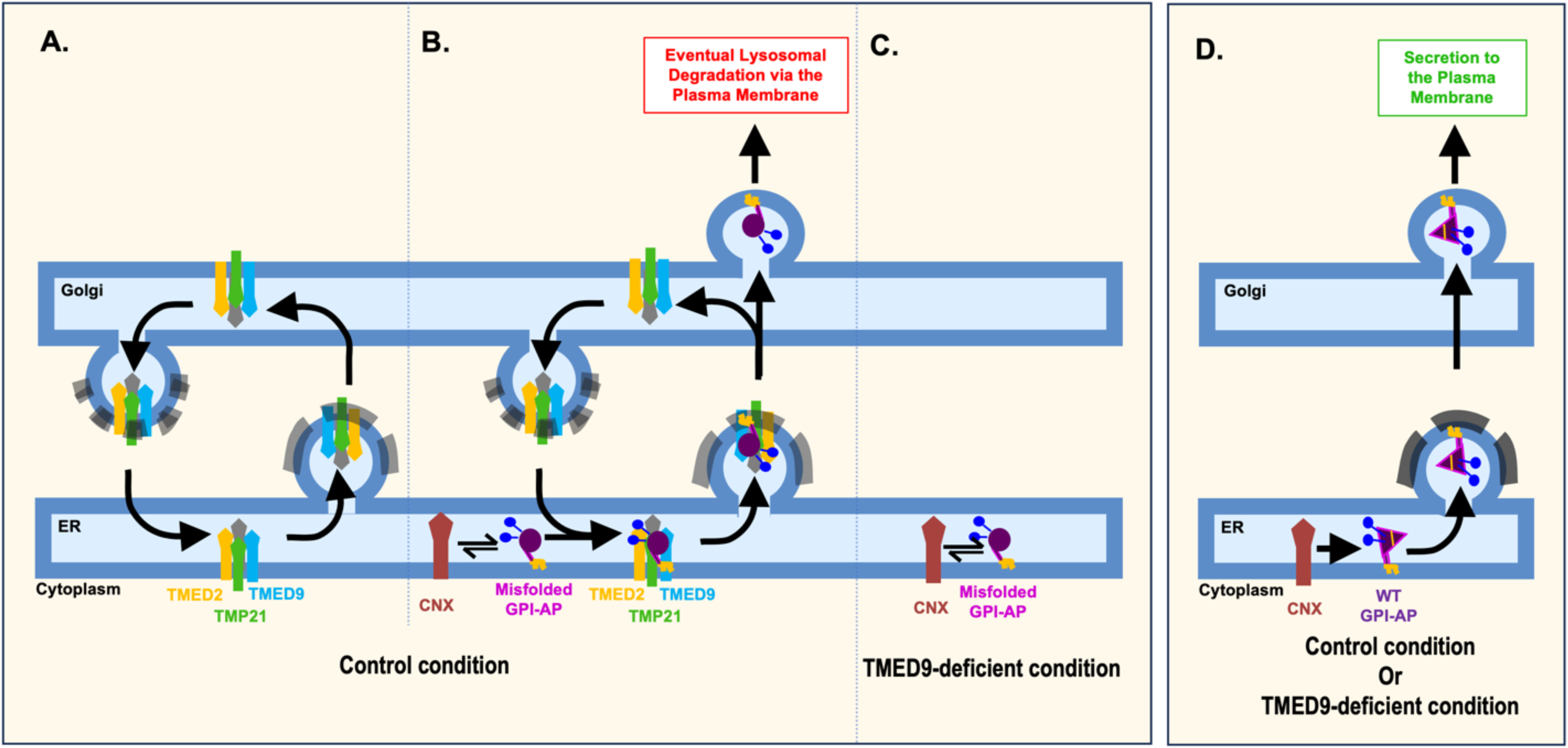
Model depicting the unique role of TMED9 in the ER-to-Golgi clearance of misfolded GPI-APs via the RESET pathway. (**A**) At steady-state, TMED9 (cyan), in conjunction with TMP21 (green), TMED2 (gold) and likely other export machinery (gray), such as other p24-family members, cargo receptors, COPII and COPI coat proteins (gray), traffic between the ER and Golgi, creating a pre-existing pathway for the ER-to-Golgi transport of select substrates. **(B)** Upon release from calnexin (red) during steady-state or ER-stress conditions, misfolded GPI-anchored proteins (GPI-APs) (depicted as purple circles) piggyback with p24-family members to access these p24-family populated ER-exit sites. **(C)** When TMED9 is depleted from the ER either by siRNA knockdown or by BRD4780-treatment, the TMED9 and p24-family populated ER-exit pathway collapses and misfolded GPI-APs are unable to exit the ER to the Golgi. Instead, they remain in association with calnexin. **(D)** By contrast to misfolded GPI-APs, properly folded GPI-APs are able to exit the ER to the Golgi for subsequent secretion to the plasma membrane regardless of the presence or absence of TMED9. Properly folded GPI-APs are depicted as purple triangles with a yellow stripe to indicate intact disulfide bonds. The p24-family members are not depicted in this panel.

**Video 1: Thapsigargin induces rapid ER-export of YFP-PrP* to the Golgi, but does not alter CER-TMED9 localization at the population level.** Time-lapse images of a YFP-PrP* NRK cell that is co-expressing CER-TMED9 and Golgi-marker, FusionRed (FusRed)-SiT. Images were collected at 5 min intervals starting after the addition of thapsigargin. A zoom box marks the region of the cell that was magnified on the right. Scale bar represents 10 µm. This Video is associated with Figure 1E.

**Video 2: TMED9 knockdown blocks RESET and consequent anti-GFP-antibody uptake in YFP-PrP* NRK cells.** Time-lapse images of YFP-PrP* NRK cells that were treated with siTMED9. Images were collected immediately after the addition of thapsigargin, Alexa Fluor 647 (AF647)-conjugated rabbit anti-GFP antibodies, and bafilomycin A1. A field of view that contained one cell with complete knockdown vs one cell with incomplete knockdown was selected for comparison. Scale bar represents 10 µm. This Video is associated with Supp. Figure 2C.

**Video 3: BRD4780-induced ER-export of TMED9 and block on RESET are reversible within 30 min of treatment.** Time-lapse images collected at 5 min intervals of a YFP-PrP* NRK cell that was co-transfected with CER-TMED9 and mCherry-Sec61β. As specified in text in the video, the cell was treated with CHX + BRD4780 for 30min, then washed with medium containing CHX-only for 30 min, and then treated with CHX + thapsigargin (TG). Scale bar represents 10 µm. This Video is associated with Figure 7A.

## Notes

### Competing Interest Statement

The authors have declared no competing interest.

### Summary of Updates

(1) edited for clarity (2) new experiment added to Supp. Figure S3 (3) updated model in Supp. Figure S8

## References

1. Princiotta MF, Finzi D, Qian SB, Gibbs J, Schuchmann S, Buttgereit F, et al. Quantitating protein synthesis, degradation, and endogenous antigen processing. Immunity. 2003;18(3):343–54. doi: 10.1016/s1074-7613(03)00051-7. PubMed PMID: 12648452.

2. Dobson CM. Protein folding and misfolding. Nature. 2003;426(6968):884–90. doi: 10.1038/nature02261. PubMed PMID: 14685248.

3. Hartl FU, Bracher A, Hayer-Hartl M. Molecular chaperones in protein folding and proteostasis. Nature. 2011;475(7356):324–32. Epub 20110720. doi: 10.1038/nature10317. PubMed PMID: 21776078.

4. Braakman I, Hebert DN. Protein folding in the endoplasmic reticulum. Cold Spring Harb Perspect Biol. 2013;5(5):a013201. Epub 20130501. doi: 10.1101/cshperspect.a013201. PubMed PMID: 23637286; PubMed Central PMCID: PMCPMC3632058.

5. Anelli T, Sitia R. Protein quality control in the early secretory pathway. EMBO J. 2008;27(2):315–27. doi: 10.1038/sj.emboj.7601974. PubMed PMID: 18216874; PubMed Central PMCID: PMCPMC2234347.

6. Phillips BP, Gomez-Navarro N, Miller EA. Protein quality control in the endoplasmic reticulum. Curr Opin Cell Biol. 2020;65:96–102. Epub 20200511. doi: 10.1016/j.ceb.2020.04.002. PubMed PMID: 32408120; PubMed Central PMCID: PMCPMC7588826.

7. Ruggiano A, Foresti O, Carvalho P. Quality control: ER-associated degradation: protein quality control and beyond. J Cell Biol. 2014;204(6):869–79. doi: 10.1083/jcb.201312042. PubMed PMID: 24637321; PubMed Central PMCID: PMCPMC3998802.

8. Sun Z, Brodsky JL. Protein quality control in the secretory pathway. J Cell Biol. 2019;218(10):3171–87. Epub 20190919. doi: 10.1083/jcb.201906047. PubMed PMID: 31537714; PubMed Central PMCID: PMCPMC6781448.

9. Molinari M. ER-phagy responses in yeast, plants, and mammalian cells and their crosstalk with UPR and ERAD. Dev Cell. 2021;56(7):949–66. Epub 20210324. doi: 10.1016/j.devcel.2021.03.005. PubMed PMID: 33765438.

10. De Leonibus C, Cinque L, Settembre C. Emerging lysosomal pathways for quality control at the endoplasmic reticulum. FEBS Lett. 2019;593(17):2319–29. Epub 20190813. doi: 10.1002/1873-3468.13571. PubMed PMID: 31388984.

11. Wilkinson S. Emerging Principles of Selective ER Autophagy. J Mol Biol. 2020;432(1):185–205. Epub 20190514. doi: 10.1016/j.jmb.2019.05.012. PubMed PMID: 31100386; PubMed Central PMCID: PMCPMC6971691.

12. Satpute-Krishnan P, Ajinkya M, Bhat S, Itakura E, Hegde RS, Lippincott-Schwartz J. ER stress-induced clearance of misfolded GPI-anchored proteins via the secretory pathway. Cell. 2014;158(3):522–33. doi: 10.1016/j.cell.2014.06.026. PubMed PMID: 25083867; PubMed Central PMCID: PMCPMC4121523.

13. Ashok A, Hegde RS. Selective processing and metabolism of disease-causing mutant prion proteins. PLoS Pathog. 2009;5(6):e1000479. Epub 2009/06/23. doi: 10.1371/journal.ppat.1000479. PubMed PMID: 19543376; PubMed Central PMCID: PMCPMC2691595.

14. Cheatham AM, Sharma NR, Satpute-Krishnan P. Competition for calnexin binding regulates secretion and turnover of misfolded GPI-anchored proteins. J Cell Biol. 2023;222(10). Epub 20230913. doi: 10.1083/jcb.202108160. PubMed PMID: 37702712; PubMed Central PMCID: PMCPMC10499038.

15. Guo XY, Liu YS, Gao XD, Kinoshita T, Fujita M. Calnexin mediates the maturation of GPI-anchors through ER retention. J Biol Chem. 2020;295(48):16393–410. Epub 2020/09/25. doi: 10.1074/jbc.RA120.015577. PubMed PMID: 32967966; PubMed Central PMCID: PMCPMC7705322.

16. Belden WJ, Barlowe C. Erv25p, a component of COPII-coated vesicles, forms a complex with Emp24p that is required for efficient endoplasmic reticulum to Golgi transport. J Biol Chem. 1996;271(43):26939–46. doi: 10.1074/jbc.271.43.26939. PubMed PMID: 8900179.

17. Dominguez M, Dejgaard K, Fullekrug J, Dahan S, Fazel A, Paccaud JP, et al. gp25L/emp24/p24 protein family members of the cis-Golgi network bind both COP I and II coatomer. J Cell Biol. 1998;140(4):751–65. doi: 10.1083/jcb.140.4.751. PubMed PMID: 9472029; PubMed Central PMCID: PMCPMC2141742.

18. Strating JR, van Bakel NH, Leunissen JA, Martens GJ. A comprehensive overview of the vertebrate p24 family: identification of a novel tissue-specifically expressed member. Mol Biol Evol. 2009;26(8):1707–14. Epub 20090508. doi: 10.1093/molbev/msp099. PubMed PMID: 19429673.

19. Schimmoller F, Singer-Kruger B, Schroder S, Kruger U, Barlowe C, Riezman H. The absence of Emp24p, a component of ER-derived COPII-coated vesicles, causes a defect in transport of selected proteins to the Golgi. EMBO J. 1995;14(7):1329–39. doi: 10.1002/j.1460-2075.1995.tb07119.x. PubMed PMID: 7729411; PubMed Central PMCID: PMCPMC398218.

20. Zavodszky E, Hegde RS. Misfolded GPI-anchored proteins are escorted through the secretory pathway by ER-derived factors. Elife. 2019;8. Epub 20190516. doi: 10.7554/eLife.46740. PubMed PMID: 31094677; PubMed Central PMCID: PMCPMC6541436.

21. Roberts BS, Satpute-Krishnan P. The many hats of transmembrane emp24 domain protein TMED9 in secretory pathway homeostasis. Front Cell Dev Biol. 2022;10:1096899. Epub 20230116. doi: 10.3389/fcell.2022.1096899. PubMed PMID: 36733337; PubMed Central PMCID: PMCPMC9888432.

22. Dvela-Levitt M, Kost-Alimova M, Emani M, Kohnert E, Thompson R, Sidhom EH, et al. Small Molecule Targets TMED9 and Promotes Lysosomal Degradation to Reverse Proteinopathy. Cell. 2019;178(3):521–35 e23. doi: 10.1016/j.cell.2019.07.002. PubMed PMID: 31348885.

23. Xiao L, Pi X, Goss AC, El-Baba T, Ehrmann JF, Grinkevich E, et al. Molecular basis of TMED9 oligomerization and entrapment of misfolded protein cargo in the early secretory pathway. Sci Adv. 2024;10(38):eadp2221. Epub 20240920. doi: 10.1126/sciadv.adp2221. PubMed PMID: 39303030; PubMed Central PMCID: PMCPMC11414720.

24. Roberts BS, Mitra D, Abishek S, Beher R, Satpute-Krishnan P. The p24-family and COPII subunit SEC24C facilitate the clearance of alpha1-antitrypsin Z from the endoplasmic reticulum to lysosomes. Mol Biol Cell. 2024;35(3):ar45. Epub 20240131. doi: 10.1091/mbc.E23-06-0257. PubMed PMID: 38294851.

25. Rane NS, Yonkovich JL, Hegde RS. Protection from cytosolic prion protein toxicity by modulation of protein translocation. EMBO J. 2004;23(23):4550–9. Epub 20041104. doi: 10.1038/sj.emboj.7600462. PubMed PMID: 15526034; PubMed Central PMCID: PMCPMC533048.

26. Rane NS, Chakrabarti O, Feigenbaum L, Hegde RS. Signal sequence insufficiency contributes to neurodegeneration caused by transmembrane prion protein. J Cell Biol. 2010;188(4):515–26. Epub 20100215. doi: 10.1083/jcb.200911115. PubMed PMID: 20156965; PubMed Central PMCID: PMCPMC2828915.

27. Liu YS, Guo XY, Hirata T, Rong Y, Motooka D, Kitajima T, et al. N-Glycan-dependent protein folding and endoplasmic reticulum retention regulate GPI-anchor processing. J Cell Biol. 2018;217(2):585–99. Epub 2017/12/20. doi: 10.1083/jcb.201706135. PubMed PMID: 29255114; PubMed Central PMCID: PMCPMC5800811.

28. Voeltz GK, Rolls MM, Rapoport TA. Structural organization of the endoplasmic reticulum. EMBO Rep. 2002;3(10):944–50. doi: 10.1093/embo-reports/kvf202. PubMed PMID: 12370207; PubMed Central PMCID: PMCPMC1307613.

29. Cole NB, Smith CL, Sciaky N, Terasaki M, Edidin M, Lippincott-Schwartz J. Diffusional mobility of Golgi proteins in membranes of living cells. Science. 1996;273(5276):797-801. doi: 10.1126/science.273.5276.797. PubMed PMID: 8670420.

30. Presley JF, Cole NB, Schroer TA, Hirschberg K, Zaal KJ, Lippincott-Schwartz J. ER-to-Golgi transport visualized in living cells. Nature. 1997;389(6646):81-5. doi: 10.1038/38001. PubMed PMID: 9288971.

31. Wei JH, Seemann J. Unraveling the Golgi ribbon. Traffic. 2010;11(11):1391–400. doi: 10.1111/j.1600-0854.2010.01114.x. PubMed PMID: 21040294; PubMed Central PMCID: PMCPMC4221251.

32. Ladinsky MS, Mastronarde DN, McIntosh JR, Howell KE, Staehelin LA. Golgi structure in three dimensions: functional insights from the normal rat kidney cell. J Cell Biol. 1999;144(6):1135–49. doi: 10.1083/jcb.144.6.1135. PubMed PMID: 10087259; PubMed Central PMCID: PMCPMC2150572.

33. Yoshimori T, Yamamoto A, Moriyama Y, Futai M, Tashiro Y. Bafilomycin A1, a specific inhibitor of vacuolar-type H(+)-ATPase, inhibits acidification and protein degradation in lysosomes of cultured cells. J Biol Chem. 1991;266(26):17707–12. PubMed PMID: 1832676.

34. Tashima Y, Hirata T, Maeda Y, Murakami Y, Kinoshita T. Differential use of p24 family members as cargo receptors for the transport of glycosylphosphatidylinositol-anchored proteins and Wnt1. J Biochem. 2022;171(1):75–83. doi: 10.1093/jb/mvab108. PubMed PMID: 34647572.

35. Zhang M, Liu L, Lin X, Wang Y, Li Y, Guo Q, et al. A Translocation Pathway for Vesicle-Mediated Unconventional Protein Secretion. Cell. 2020;181(3):637–52 e15. Epub 20200408. doi: 10.1016/j.cell.2020.03.031. PubMed PMID: 32272059.

36. Obara CJ, Moore AS, Lippincott-Schwartz J. Structural Diversity within the Endoplasmic Reticulum-From the Microscale to the Nanoscale. Cold Spring Harb Perspect Biol. 2023;15(6). Epub 20230601. doi: 10.1101/cshperspect.a041259. PubMed PMID: 36123032; PubMed Central PMCID: PMCPMC10394098.

37. Nixon-Abell J, Obara CJ, Weigel AV, Li D, Legant WR, Xu CS, et al. Increased spatiotemporal resolution reveals highly dynamic dense tubular matrices in the peripheral ER. Science. 2016;354(6311). Epub 20161027. doi: 10.1126/science.aaf3928. PubMed PMID: 27789813; PubMed Central PMCID: PMCPMC6528812.

38. Fregno I, Fasana E, Bergmann TJ, Raimondi A, Loi M, Solda T, et al. ER-to-lysosome-associated degradation of proteasome-resistant ATZ polymers occurs via receptor-mediated vesicular transport. EMBO J. 2018;37(17). Epub 20180803. doi: 10.15252/embj.201899259. PubMed PMID: 30076131; PubMed Central PMCID: PMCPMC6120659.

39. Doms RW, Russ G, Yewdell JW. Brefeldin A redistributes resident and itinerant Golgi proteins to the endoplasmic reticulum. J Cell Biol. 1989;109(1):61–72. doi: 10.1083/jcb.109.1.61. PubMed PMID: 2745557; PubMed Central PMCID: PMCPMC2115463.

40. Lippincott-Schwartz J, Yuan LC, Bonifacino JS, Klausner RD. Rapid redistribution of Golgi proteins into the ER in cells treated with brefeldin A: evidence for membrane cycling from Golgi to ER. Cell. 1989;56(5):801–13. doi: 10.1016/0092-8674(89)90685-5. PubMed PMID: 2647301; PubMed Central PMCID: PMCPMC7173269.

41. Fujiwara T, Oda K, Yokota S, Takatsuki A, Ikehara Y. Brefeldin A causes disassembly of the Golgi complex and accumulation of secretory proteins in the endoplasmic reticulum. J Biol Chem. 1988;263(34):18545–52. PubMed PMID: 3192548.

42. Sciaky N, Presley J, Smith C, Zaal KJ, Cole N, Moreira JE, et al. Golgi tubule traffic and the effects of brefeldin A visualized in living cells. J Cell Biol. 1997;139(5):1137–55. doi: 10.1083/jcb.139.5.1137. PubMed PMID: 9382862; PubMed Central PMCID: PMCPMC2140213.

43. Freeze HH, Kranz C. Endoglycosidase and glycoamidase release of N-linked glycans. Curr Protoc Mol Biol. 2010;Chapter 17:Unit 17 3A. doi: 10.1002/0471142727.mb1713as89. PubMed PMID: 20069534; PubMed Central PMCID: PMCPMC3869378.

44. Varki A CR, Esko JD, Stanley P, Hart GW, Aebi M, Mohnen D, Kinoshita T, Packer NH, Prestegard JH, Schnaar RL, Seeberger PH Essentials of Glycobiology [Internet]. In: Varki A, Cummings RD, Esko JD Stanley P, Hart GW, Aebi M, et al., editors. Essentials of Glycobiology. 4th ed. Cold Spring Harbor (NY): Cold Spring Harbor Laboratory Press; 2022.

45. Hammond C, Braakman I, Helenius A. Role of N-linked oligosaccharide recognition, glucose trimming, and calnexin in glycoprotein folding and quality control. Proc Natl Acad Sci U S A. 1994;91(3):913–7. doi: 10.1073/pnas.91.3.913. PubMed PMID: 8302866; PubMed Central PMCID: PMCPMC521423.

46. Bhide GP, Colley KJ. Sialylation of N-glycans: mechanism, cellular compartmentalization and function. Histochem Cell Biol. 2017;147(2):149–74. Epub 20161214. doi: 10.1007/s00418-016-1520-x. PubMed PMID: 27975143; PubMed Central PMCID: PMCPMC7088086.

47. Rabouille C, Hui N, Hunte F, Kieckbusch R, Berger EG, Warren G, et al. Mapping the distribution of Golgi enzymes involved in the construction of complex oligosaccharides. J Cell Sci. 1995;108 ( Pt 4):1617–27. doi: 10.1242/jcs.108.4.1617. PubMed PMID: 7615680.

48. Fullekrug J, Suganuma T, Tang BL, Hong W, Storrie B, Nilsson T. Localization and recycling of gp27 (hp24gamma3): complex formation with other p24 family members. Mol Biol Cell. 1999;10(6):1939–55. doi: 10.1091/mbc.10.6.1939. PubMed PMID: 10359607; PubMed Central PMCID: PMCPMC25391.

49. Lavoie C, Paiement J, Dominguez M, Roy L, Dahan S, Gushue JN, et al. Roles for alpha(2)p24 and COPI in endoplasmic reticulum cargo exit site formation. J Cell Biol. 1999;146(2):285–99. doi: 10.1083/jcb.146.2.285. PubMed PMID: 10427085; PubMed Central PMCID: PMCPMC3206572.

50. Yoshimura SI, Nakamura N, Barr FA, Misumi Y, Ikehara Y, Ohno H, et al. Direct targeting of cis-Golgi matrix proteins to the Golgi apparatus. J Cell Sci. 2001;114(Pt 22):4105–15. doi: 10.1242/jcs.114.22.4105. PubMed PMID: 11739642.

51. Montesinos JC, Sturm S, Langhans M, Hillmer S, Marcote MJ, Robinson DG, et al. Coupled transport of Arabidopsis p24 proteins at the ER-Golgi interface. J Exp Bot. 2012;63(11):4243–61. Epub 20120510. doi: 10.1093/jxb/ers112. PubMed PMID: 22577184; PubMed Central PMCID: PMCPMC3398454.

52. Fiedler K, Veit M, Stamnes MA, Rothman JE. Bimodal interaction of coatomer with the p24 family of putative cargo receptors. Science. 1996;273(5280):1396-9. doi: 10.1126/science.273.5280.1396. PubMed PMID: 8703076.

53. Strating JR, Martens GJ. The p24 family and selective transport processes at the ER-Golgi interface. Biol Cell. 2009;101(9):495–509. doi: 10.1042/BC20080233. PubMed PMID: 19566487.

54. Mitrovic S, Ben-Tekaya H, Koegler E, Gruenberg J, Hauri HP. The cargo receptors Surf4, endoplasmic reticulum-Golgi intermediate compartment (ERGIC)-53, and p25 are required to maintain the architecture of ERGIC and Golgi. Mol Biol Cell. 2008;19(5):1976–90. Epub 20080220. doi: 10.1091/mbc.e07-10-0989. PubMed PMID: 18287528; PubMed Central PMCID: PMCPMC2366877.

55. Gomez-Navarro N, Melero A, Li XH, Boulanger J, Kukulski W, Miller EA. Cargo crowding contributes to sorting stringency in COPII vesicles. J Cell Biol. 2020;219(7). doi: 10.1083/jcb.201806038. PubMed PMID: 32406500; PubMed Central PMCID: PMCPMC7300426.

56. Castillon GA, Aguilera-Romero A, Manzano-Lopez J, Epstein S, Kajiwara K, Funato K, et al. The yeast p24 complex regulates GPI-anchored protein transport and quality control by monitoring anchor remodeling. Mol Biol Cell. 2011;22(16):2924–36. Epub 20110616. doi: 10.1091/mbc.E11-04-0294. PubMed PMID: 21680708; PubMed Central PMCID: PMCPMC3154887.

57. Muniz M, Nuoffer C, Hauri HP, Riezman H. The Emp24 complex recruits a specific cargo molecule into endoplasmic reticulum-derived vesicles. J Cell Biol. 2000;148(5):925–30. doi: 10.1083/jcb.148.5.925. PubMed PMID: 10704443; PubMed Central PMCID: PMCPMC2174538.

58. Bonnon C, Wendeler MW, Paccaud JP, Hauri HP. Selective export of human GPI-anchored proteins from the endoplasmic reticulum. J Cell Sci. 2010;123(Pt 10):1705–15. Epub 20100427. doi: 10.1242/jcs.062950. PubMed PMID: 20427317.

59. Bernat-Silvestre C, De Sousa Vieira V, Sanchez-Simarro J, Pastor-Cantizano N, Hawes C, Marcote MJ, et al. p24 Family Proteins Are Involved in Transport to the Plasma Membrane of GPI-Anchored Proteins in Plants. Plant Physiol. 2020;184(3):1333–47. Epub 20200908. doi: 10.1104/pp.20.00880. PubMed PMID: 32900981; PubMed Central PMCID: PMCPMC7608175.

60. Theiler R, Fujita M, Nagae M, Yamaguchi Y, Maeda Y, Kinoshita T. The alpha-helical region in p24gamma2 subunit of p24 protein cargo receptor is pivotal for the recognition and transport of glycosylphosphatidylinositol-anchored proteins. J Biol Chem. 2014;289(24):16835–43. Epub 20140428. doi: 10.1074/jbc.M114.568311. PubMed PMID: 24778190; PubMed Central PMCID: PMCPMC4059126.

61. Takida S, Maeda Y, Kinoshita T. Mammalian GPI-anchored proteins require p24 proteins for their efficient transport from the ER to the plasma membrane. Biochem J. 2008;409(2):555–62. doi: 10.1042/BJ20070234. PubMed PMID: 17927562.

62. D’Arcangelo JG, Crissman J, Pagant S, Copic A, Latham CF, Snapp EL, et al. Traffic of p24 Proteins and COPII Coat Composition Mutually Influence Membrane Scaffolding. Curr Biol. 2015;25(10):1296–305. Epub 20150430. doi: 10.1016/j.cub.2015.03.029. PubMed PMID: 25936552; PubMed Central PMCID: PMCPMC4439346.

63. Emery G, Parton RG, Rojo M, Gruenberg J. The trans-membrane protein p25 forms highly specialized domains that regulate membrane composition and dynamics. J Cell Sci. 2003;116(Pt 23):4821–32. doi: 10.1242/jcs.00802. PubMed PMID: 14600267.

64. Loizides-Mangold U, David FP, Nesatyy VJ, Kinoshita T, Riezman H. Glycosylphosphatidylinositol anchors regulate glycosphingolipid levels. J Lipid Res. 2012;53(8):1522–34. Epub 20120524. doi: 10.1194/jlr.M025692. PubMed PMID: 22628614; PubMed Central PMCID: PMCPMC3540855.

65. Maldutyte J, Li XH, Gomez-Navarro N, Robertson EG, Miller EA. ER export via SURF4 uses diverse mechanisms of both client and coat engagement. J Cell Biol. 2025;224(1). Epub 20241112. doi: 10.1083/jcb.202406103. PubMed PMID: 39531033; PubMed Central PMCID: PMCPMC11557686.

66. Contreras FX, Ernst AM, Haberkant P, Bjorkholm P, Lindahl E, Gonen B, et al. Molecular recognition of a single sphingolipid species by a protein’s transmembrane domain. Nature. 2012;481(7382):525-9. Epub 20120109. doi: 10.1038/nature10742. PubMed PMID: 22230960.

67. Anwar MU, Sergeeva OA, Abrami L, Mesquita FS, Lukonin I, Amen T, et al. ER-Golgi-localized proteins TMED2 and TMED10 control the formation of plasma membrane lipid nanodomains. Dev Cell. 2022;57(19):2334–46 e8. Epub 20220928. doi: 10.1016/j.devcel.2022.09.004. PubMed PMID: 36174556.

68. Dvela-Levitt M, Shaw JL, Greka A. A Rare Kidney Disease To Cure Them All? Towards Mechanism-Based Therapies for Proteinopathies. Trends Mol Med. 2021;27(4):394–409. Epub 20201216. doi: 10.1016/j.molmed.2020.11.008. PubMed PMID: 33341352.

69. Fujita M, Watanabe R, Jaensch N, Romanova-Michaelides M, Satoh T, Kato M, et al. Sorting of GPI-anchored proteins into ER exit sites by p24 proteins is dependent on remodeled GPI. J Cell Biol. 2011;194(1):61–75. Epub 20110704. doi: 10.1083/jcb.201012074. PubMed PMID: 21727194; PubMed Central PMCID: PMCPMC3135397.

70. Fehlinger A, Wolf H, Hossinger A, Duernberger Y, Pleschka C, Riemschoss K, et al. Prion strains depend on different endocytic routes for productive infection. Sci Rep. 2017;7(1):6923. Epub 20170731. doi: 10.1038/s41598-017-07260-2. PubMed PMID: 28761068; PubMed Central PMCID: PMCPMC5537368.

71. Chassefeyre R, Chaiamarit T, Verhelle A, Novak SW, Andrade LR, Leitao ADG, et al. Endosomal sorting drives the formation of axonal prion protein endoggresomes. Sci Adv. 2021;7(52):eabg3693. Epub 20211222. doi: 10.1126/sciadv.abg3693. PubMed PMID: 34936461; PubMed Central PMCID: PMCPMC8694590.

72. Jeffrey M, Goodsir CM, Bruce M, McBride PA, Scott JR, Halliday WG. Correlative light and electron microscopy studies of PrP localisation in 87V scrapie. Brain Res. 1994;656(2):329–43. Epub 1994/09/12. doi: 10.1016/0006-8993(94)91477-x. PubMed PMID: 7820594.

73. Scheckel C, Aguzzi A. Prions, prionoids and protein misfolding disorders. Nat Rev Genet. 2018;19(7):405–18. Epub 2018/05/02. doi: 10.1038/s41576-018-0011-4. PubMed PMID: 29713012.

74. Hailey DW, Rambold AS, Satpute-Krishnan P, Mitra K, Sougrat R, Kim PK, et al. Mitochondria supply membranes for autophagosome biogenesis during starvation. Cell. 2010;141(4):656–67. doi: 10.1016/j.cell.2010.04.009. PubMed PMID: 20478256; PubMed Central PMCID: PMCPMC3059894.

75. Mantei N, Villa M, Enzler T, Wacker H, Boll W, James P, et al. Complete primary structure of human and rabbit lactase-phlorizin hydrolase: implications for biosynthesis, membrane anchoring and evolution of the enzyme. EMBO J. 1988;7(9):2705–13. doi: 10.1002/j.1460-2075.1988.tb03124.x. PubMed PMID: 2460343; PubMed Central PMCID: PMCPMC457059.

76. Hauri HP, Sterchi EE, Bienz D, Fransen JA, Marxer A. Expression and intracellular transport of microvillus membrane hydrolases in human intestinal epithelial cells. J Cell Biol. 1985;101(3):838–51. doi: 10.1083/jcb.101.3.838. PubMed PMID: 3897250; PubMed Central PMCID: PMCPMC2113743.

77. Naim HY, Sterchi EE, Lentze MJ. Biosynthesis and maturation of lactase-phlorizin hydrolase in the human small intestinal epithelial cells. Biochem J. 1987;241(2):427–34. doi: 10.1042/bj2410427. PubMed PMID: 3109375; PubMed Central PMCID: PMCPMC1147578.

78. Ouwendijk J, Peters WJ, van de Vorstenbosch RA, Ginsel LA, Naim HY, Fransen JA. Routing and processing of lactase-phlorizin hydrolase in transfected Caco-2 cells. J Biol Chem. 1998;273(12):6650–5. doi: 10.1074/jbc.273.12.6650. PubMed PMID: 9506961.

79. Keller P, Toomre D, Diaz E, White J, Simons K. Multicolour imaging of post-Golgi sorting and trafficking in live cells. Nat Cell Biol. 2001;3(2):140–9. doi: 10.1038/35055042. PubMed PMID: 11175746.

80. Shemiakina, II, Ermakova GV, Cranfill PJ, Baird MA, Evans RA, Souslova EA, et al. A monomeric red fluorescent protein with low cytotoxicity. Nat Commun. 2012;3:1204. doi: 10.1038/ncomms2208. PubMed PMID: 23149748.

